# Cell-Free Biosynthesis to Evaluate Lasso Peptide Formation and Enzyme-Substrate Tolerance

**DOI:** 10.1101/2020.11.25.399105

**Authors:** Yuanyuan Si, Ashley M. Kretsch, Laura M. Daigh, Mark J. Burk, Douglas A. Mitchell

## Abstract

Lasso peptides are ribosomally synthesized and post-translationally modified peptide (RiPP) natural products that display a unique lariat-like structure. Owing to a rigid topology, lasso peptides are unusually stable towards heat and proteolytic degradation. Some lasso peptides have been shown to bind human cell-surface receptors and exhibit anticancer properties, while others display antibacterial or antiviral activities. Known lasso peptides are produced by bacteria and genome-mining studies indicate that lasso peptides are a relatively prevalent RiPP class; however, the discovery, isolation, and characterization of lasso peptides are constrained by the lack of an efficient production system. In this study, we employ a cell-free biosynthesis (CFB) strategy to address the longstanding challenges associated with lasso peptide production. We report the successful formation of a diverse array of lasso peptides that include known examples as well as a new predicted lasso peptide from *Thermobifida halotolerans*. We further demonstrate the utility of CFB to rapidly generate and characterize multisite precursor peptide variants in order to evaluate the substrate tolerance of the biosynthetic pathway. We show that the lasso-forming cyclase from the fusilassin pathway can produce millions of sequence-diverse lasso peptides via CFB with an extraordinary level of sequence permissiveness within the ring region of the lasso peptide. These data lay a firm foundation for the creation of large lasso peptide libraries using CFB to identify new variants with unique properties.

## Introduction

Lasso peptides were first identified in the early 1990s and today comprise a large and growing class of ribosomally synthesized and post-translationally modified peptides (RiPP).^1,2^ Similar to other RiPP classes, lasso peptides are formed by conversion of precursor peptides that consist of a N-terminal leader region and a C-terminal core region.^3^ After translation of the linear precursor peptide, lasso peptide biosynthesis begins with recognition of the leader region by a RiPP precursor recognition element (RRE) which guides the biosynthetic enzymes to the substrate.^4^ The leader region is then proteolytically removed by a peptidase homologous to transglutaminase (Protein Family PF13471). Subsequently, an ATP-dependent lasso cyclase, homologous to asparagine synthetase (PF00733), catalyzes macrolactam formation between the N-terminus and the carboxyl side chain of a Glu or Asp residue within the core region.^1,2,5^ To date, all characterized, naturally occurring lasso peptides contain 7–9 residues within the macrolactam ring region, although an unnatural 10-mer variant has recently been reported.^6^ During lasso peptide biosynthesis, the substrate peptide must be “pre-folded” with the C-terminal tail passing through the plane of the incipient ring, resulting in a lariat-like tertiary structure after macrolactam formation.^7^ The threaded lasso topology is stabilized either by a large, steric locking residue located below the ring (tail), or occasionally by disulfide bonds. An additional conformational constraint is sometimes provided by a large residue above the plane of the ring (loop). Further chemical complexity can be introduced into the lasso peptide scaffold through post-translational modifications, including glycosylation,^8^ phosphorylation,^9^ acetylation,^10^ deimination,^11^ methylation,^12^ epimerization,^13^ and hydroxylation.^14^

The distinctive and rigid globular shape of lasso peptides typically imparts extraordinary resistance to thermal and proteolytic degradation,^2,15^ which are unusual traits for a peptide. Although nearly 80 lasso peptides have been described, less than 30 have reported biological activity.^16^ Many of the known activities involve antibacterial properties that entail binding a variety of distinct targets, including RNA polymerase,^17–21^ prolyl endopeptidase,^22^ lipid II,^23^ mycobacterial ClpC1 ATPase,^24^ and blocking *fsr* quorum-sensing.^25^ Other lasso peptides, discovered by targeted screening, antagonize various human GPCRs, such as the glucagon^26^ and endothelin B receptors.^27,28^ Additionally, anti-HIV^29,30^ and anticancer^31^ activities have also been described.

Recent studies have explored the substrate tolerance of several lasso peptide biosynthetic enzymes to access greater structural diversity and evaluate the potential of lasso peptides for medical applications. For example, multiple amino acid substitutions and systematic structure-activity analysis of microcin J25 found that the biosynthetic enzymes tolerated amino acid substitutions at over 80% of the lasso peptide residues.^32–34^ Epitope grafting within the loop region endowed microcin J25 with integrin receptor-binding activity, whereas the natural scaffold was inactive.^35^ Furthermore, a previous study found that the core region of the lasso peptide citrulassin A (D8E) could be fused to the FusA leader region to create a chimeric precursor peptide. The fusilassin leader peptidase (FusB) and lasso cyclase (FusC) successfully produced the citrulassin A variant (D8E), which suggested the fusilassin pathway may be broadly tolerant of alternative substrates, although wildtype citrulassin A (containing Asp8 for macrolactam formation) was not tolerated.^6^ Finally, unnatural amino acids have been incorporated into several lasso peptides, including capistruin^36^ and microcin J25.^37^

The increase in publicly available genomes and availability of the open-access bioinformatic tool RODEO (Rapid ORF Description & Evaluation Online)^11^ enabled the prediction of over 3,000 high-confidence lasso biosynthetic gene clusters (BGCs) in the NCBI database (as of 2018).^6^ Despite the thousands of lasso peptides residing in databases, the isolation and characterization of new lasso peptides have been hindered by the lack of an efficient production method. Historically, new lasso peptides were isolated from the native producer,^27,38^ which can suffer from slow cellular growth and low yields. More recently, lasso peptides have been produced through heterologous expression of the BGC.^39,40^ However, heterologous expression requires time-consuming molecular cloning and growth steps and can be hampered by pathway-host incompatibilities, giving rise to poor gene expression, pathway enzyme inactivity, and product toxicity.

Cell-free biosynthesis (CFB) methods potentially offer an orthogonal, rapid, and high-throughput strategy to overcome several limitations of existing approaches for lasso peptide production. CFB reactions include three components: cell extract, energy mix, and a DNA template. The crude cell extract provides the cellular machinery for *in vitro* transcription and translation of the DNA templates.^41^ CFB eliminates the need for transformation, expression, and protein purification, which has been historically challenging for lasso peptide enzymes^6,42^. Thus, CFB can shorten the time requirement to advance from gene sequence to chemical product. Also, the use of modular DNA templates in CFB allows rapid modification of precursor sequences,^43^ streamlining the generation of lasso peptide variant libraries. Armed with an efficient production system, understanding the principles that govern lasso peptide formation and substrate tolerance could be more readily discerned, potentially allowing for the design of lasso peptides with specific biological properties.

Cell-free protein synthesis has been well-established for single protein production, with numerous examples of high yielding proteins.^44^ In contrast, there are fewer examples of CFB applied to small molecules and natural products, which require the production of multiple active enzymes to deliver a single product of interest. Some examples include 2,3-butanediol,^45^ diverse glycan motifs,^46^ and a cyclic dipeptide produced from two non-ribosomal peptide synthetases.^47^ Several mature RiPPs have been produced using cell-free methods, such as goadsporin,^48^ nisin,^49^ thiocillin,^50^ lactazole^51^. To the best of our knowledge, lasso peptides have not previously been produced in the manner.

In this study, we evaluated CFB methodology to produce a diverse array of lasso peptides, including known examples and novel lasso peptides predicted from genomic data. Furthermore, we expanded the utility of CFB to rapidly interrogate the substrate tolerance of the fusilassin biosynthetic enzymes. To enable this study, we created a randomized, multisite replacement library of the fusilassin precursor peptide by CFB. The ensuing screening and sequencing results indicated that nearly 2 million variants of FusA can be tolerated by FusC with the ring region of fusilassin being especially tolerant to substitution. These data considerably expand the sequence-structure space accessible to lasso peptides and raises their potential as customizable ligands for biotechnological applications.

## Results and Discussion

### Cell-free biosynthesis of known lasso peptides

In this study, we evaluated cell-free biosynthesis (CFB) as a potential solution to overcome longstanding challenges in lasso peptide production. We first tested four previously reported lasso peptides: burhizin (*Burkholderia rhizoxinica*),^40^ capistruin (*Burkholderia thailandensis*),^18^ fusilassin (*Thermobifida fusca*),^6,42^ and cellulassin (*Thermobifida cellulosilytica*). For burhizin and capistruin, a plasmid bearing the complete BGC was used as the DNA template. This included genes encoding the precursor peptides (*burA/capA*), RRE-leader peptidase fusion proteins (*burB/capB*), lasso cyclases (*burC/capC*), and ABC transporters (*burD/capD*, **Figures 1, S1)**. For fusilassin, a plasmid that fuses maltose-binding protein (MBP) to the N-terminus of the precursor peptide (*fusA*) was used in conjunction with a second plasmid encoding the lasso cyclase (*fusC*), RRE (*fusE*), and leader peptidase (*fusB*). For cellulassin, the BGC with the native gene organization: MBP-tagged precursor peptide (*celA*), followed by the lasso cyclase (*celC*), RRE (*celE*), and leader peptidase (*celB*) (**Figures 1, S1**) was used as the DNA template. Each plasmid was purified by maxi-prep to achieve high DNA purity and concentration. Extracts for CFB reactions were prepared from *E. coli* BL21(DE3) cells (**Supplemental Methods**). After addition of energy mix and appropriate plasmids, the CFB reactions were allowed to progress for 16 h prior to analysis by matrix-assisted laser desorption/ionization time-of-flight mass spectrometry (MALDI-TOF-MS). In each case, we readily observed the expected product mass (**Figure 1**). Peptide identity was corroborated by high-resolution and tandem MS (HRMS/MS, **Figures S2–4**), and the positions of the macrolactam linkages of burhizin, capistruin, and fusilassin were consistent with previous reports (Gly1-Glu8, Gly1-Asp9, and Trp1-Glu9, respectively).^6,40,52^ Tandem MS analysis of cellulassin has not been previously reported; however, our data confirmed the expected macrolactam linkage between Trp1-Glu9 (**Figure S5**).

**Figure 1.**
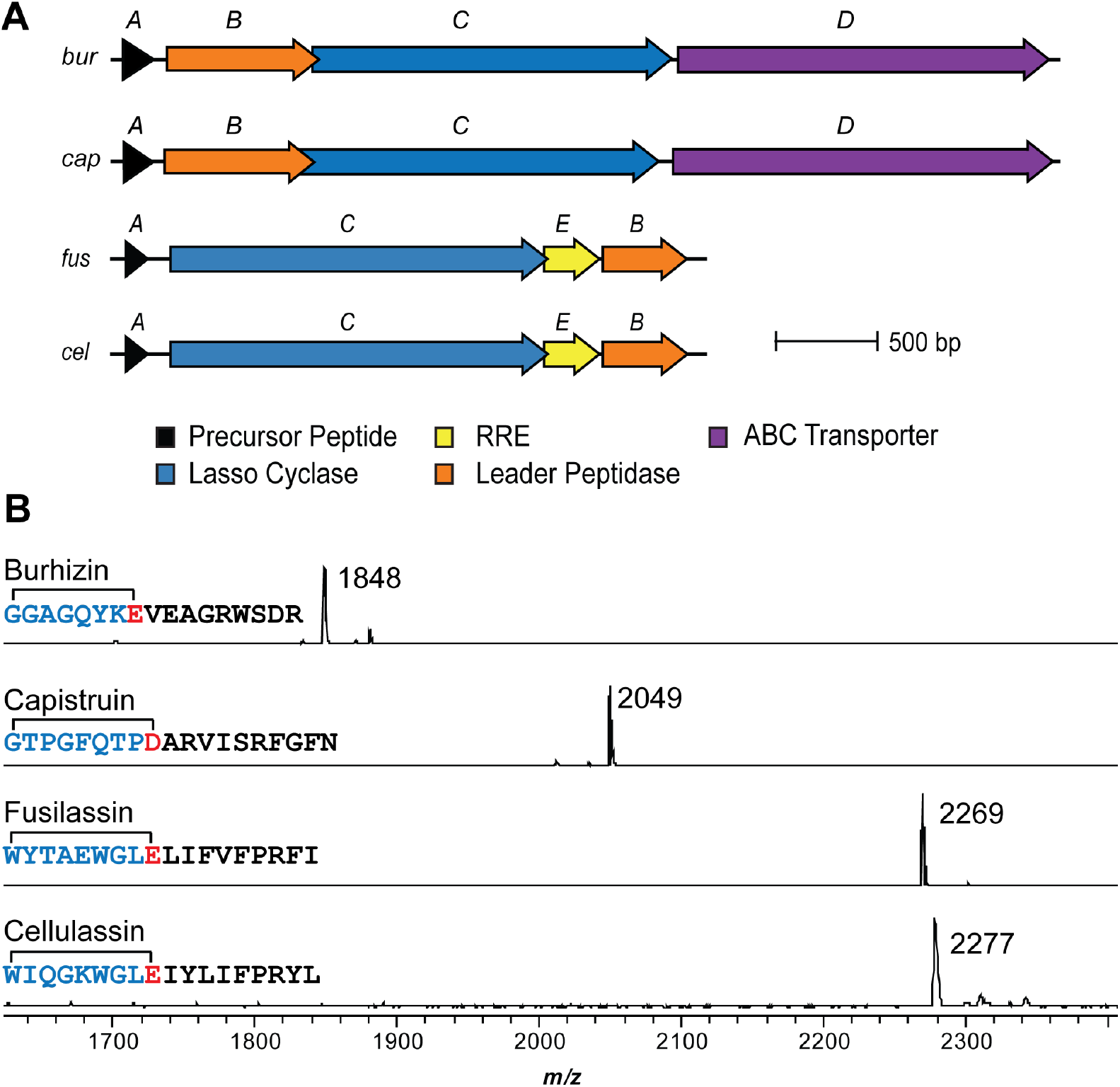
*In vitro* lasso peptide production through CFB. **(A)** The biosynthetic gene clusters provided in the CFB reaction for the production of burhizin, capistruin, fusilassin, and cellulassin respectively. RRE: RiPP leader peptide Recognition Element. **(B)** Endpoint MALDI-TOF-MS assay of burhizin (*m/z* 1848), capistruin (*m/z* 2046), fusilassin (*m/z* 2269), and cellulassin (*m/z* 2277) produced from CFB. Blue, macrolactam ring residues; Red, acceptor site. The mass label corresponds to the [M+H]^+^ ion of the lasso peptides.

Encouraged by the production of four distinct lasso peptides by CFB, we sought to determine if multiple lasso peptides could be produced in a single CFB reaction, as this would render the method more attractive for library generation. We therefore carried out CFB to simultaneously produce capistruin, burhizin, and fusilassin (from a total of four plasmids). All three lasso peptides were detected by MALDI-TOF-MS with the relative production level of burhizin correlating to amount of plasmid supplied (**Figure S6)**.

To evaluate if lasso peptides produced by CFB were in the native threaded topology, as opposed to the unthreaded, “branched-cyclic” state, we treated burhizin, fusilassin, and cellulassin with carboxypeptidase Y. All three were highly resistant to digestion, showing either no C-terminal trimming (burhizin) or low-efficiency removal of the last one or two residues, consistent with a lasso-like conformation (**Figure S7)**. After prolonged heating (2 h at 95 °C), burhizin (eight-residue ring) was still heat-resistant^40^ while fusilassin and cellulassin (both nine-residue rings) appear to unthread. Previous work has shown that fusilassin adopts a branched-cyclic form after extended exposure to organic solvent.^42^

### Scalability of CFB-produced lasso peptides

To assess the scalability of lasso peptides produced by CFB, we scaled the capistruin production reaction by 36-fold (0.75 mL). The yield of capistruin was determined using high-performance liquid chromatography (HPLC) against a standard curve of purified capistruin.^53^ Compared to the reported post-HPLC yield of 0.2 μg/mL culture from heterologous expression in *E. coli*,^18^ CFB resulted in ~200-fold higher yield (40 μg/mL, **Figure S8**). HPLC retention time analysis and HRMS/MS data yielded indistinguishable data for authentic and CFB-produced capistruin (**Figure S8**). CFB conducted at this scale permitted bioactivity evaluation using a modified RNA-polymerase inhibition assay. CFB was conducted using a plasmid encoding the fluorescent protein mCherry as a DNA template.^54^ Due to the inhibitory activity against RNA polymerase,^55^ CFB-produced capistruin supplied at 10 μM yielded a 50% reduction in mCherry production compared to a control omitting capistruin (**Figure S8**). This convenient assay could be used to evaluate any compound as a potential inhibitor of transcription and/or translation.

### CFB to produce sequence-diverse lasso peptides

The above examples demonstrated the ability of CFB to produce a diverse panel of known lasso peptides using an operationally simple procedure with native BGCs expressed from highly pure, concentrated plasmids. Our previous report demonstrated that the fusilassin biosynthetic pathway is somewhat unusual in that it tolerates substitution at the first position of the core region (naturally Trp) and ring size variants (±1 residue).^6^ This result was obtained from enzymatic reconstitution studies that required molecular cloning, expression, and purification steps for each FusA variant. By simply adding the biosynthetic genes to the CFB mixture, these results were rapidly replicated without the need for any protein purification (**Figure S9**). Besides single-site variants, the fusilassin enzymes are known to tolerate more divergent sequences. As described in previous reports, new RiPPs can be obtained by using a chimeric substrate strategy where the leader region from one precursor peptide is retained while the core region derives from a second precursor peptide.^56^ Using this approach, citrulassin A (D8E) was successfully produced by the fusilassin enzymes using a fusilassin leader-citrulassin core chimeric substrate.^6^ Formation of the citrulassin variant by the non-cognate fusilassin enzymes suggested that one robust set of substrate-tolerant lasso peptide biosynthetic enzymes could allow the production of novel lasso peptides by simply replacing the core region of the precursor peptide.

We next sought to determine if CFB and the chimeric approach could be combined to produce lasso peptides predicted from genome-mining. We first conducted a BLASTP search of the non-redundant NCBI database using FusC as the query. All proteins with >40% identity to FusC (n = 90) were re-analyzed by RODEO, confirming all were within a high-confidence lasso peptide BGC (i.e., homologs of FusB and FusE were present as well as a high-scoring precursor peptide). From the resulting list of BGCs containing FusC-like cyclases, we chose ten distinct core sequences to assess the tolerability of FusB/E and FusC for accepting chimeric substrates (**Table S1**). The genes for the chimeric peptides (FusA leader peptide, FusA_LP_, fused to a non-cognate core peptide, XxxA_CP_) were commercially synthesized with the native Glu acceptor residue retained (**Table S2**). The chimeric precursor peptides were then individually cloned into the pET28-MBP vector and purified by maxi-prep. As before, the precursor peptides were produced by CFB using highly pure plasmid DNA; however, for these reactions, the fusilassin biosynthetic enzymes (FusB, FusC, and FusE) were supplied as purified proteins (**Figure S10**). This had an anticipated two-fold positive effect: *(i)* rather than being expressed by CFB, energy resources could be preserved to elevate the production level of the precursor peptide, and *(ii)* the CFB extract contains endogenous proteases will compete for the precursor peptide. By direct addition of the biosynthetic proteins at a higher, researcher-defined concentration, the flux towards lasso peptide formation could be maximized relative to substrate degradation.

Masses corresponding to the cyclized lasso peptides of two of the ten chimeric substrates, from *Thermobifida halotolerans* and *Thermobifida cellulosilytica* (FusA_LP_-HalA_CP_, halolassin *m/z* 2047; FusA_LP_-CelA_CP_, cellulassin *m/z* 2277), were detected by MALDI-TOF-MS (**Figure 2**). Both cyclized sequences derive from the same genus as fusilassin, and their corresponding lasso cyclases show >70% similarity to FusC (**Table S1**). The lasso peptide molecular formulas and sequence identity were confirmed by HRMS/MS and consistent with a Tyr1-Glu8 and Trp1-Glu9 macrolactam for halolassin and cellulassin, respectively (**Figures S11-12**). A threaded topology for halolassin and cellulassin was supported by resistance to carboxypeptidase Y digestion. Consistent with the cellulassin produced from the native BGC, resistance to carboxypeptidase Y digestion was abolished after prolonged heating at 95 °C. Conversely, halolassin maintained a folded state after prolonged heating, as indicated by resistance to carboxypeptidase Y treatment before and after heat treatment. The enhanced thermal stability of halolassin relative to cellulassin may be attributable to having a smaller ring (**Figure S13**).^15^

**Figure 2.**
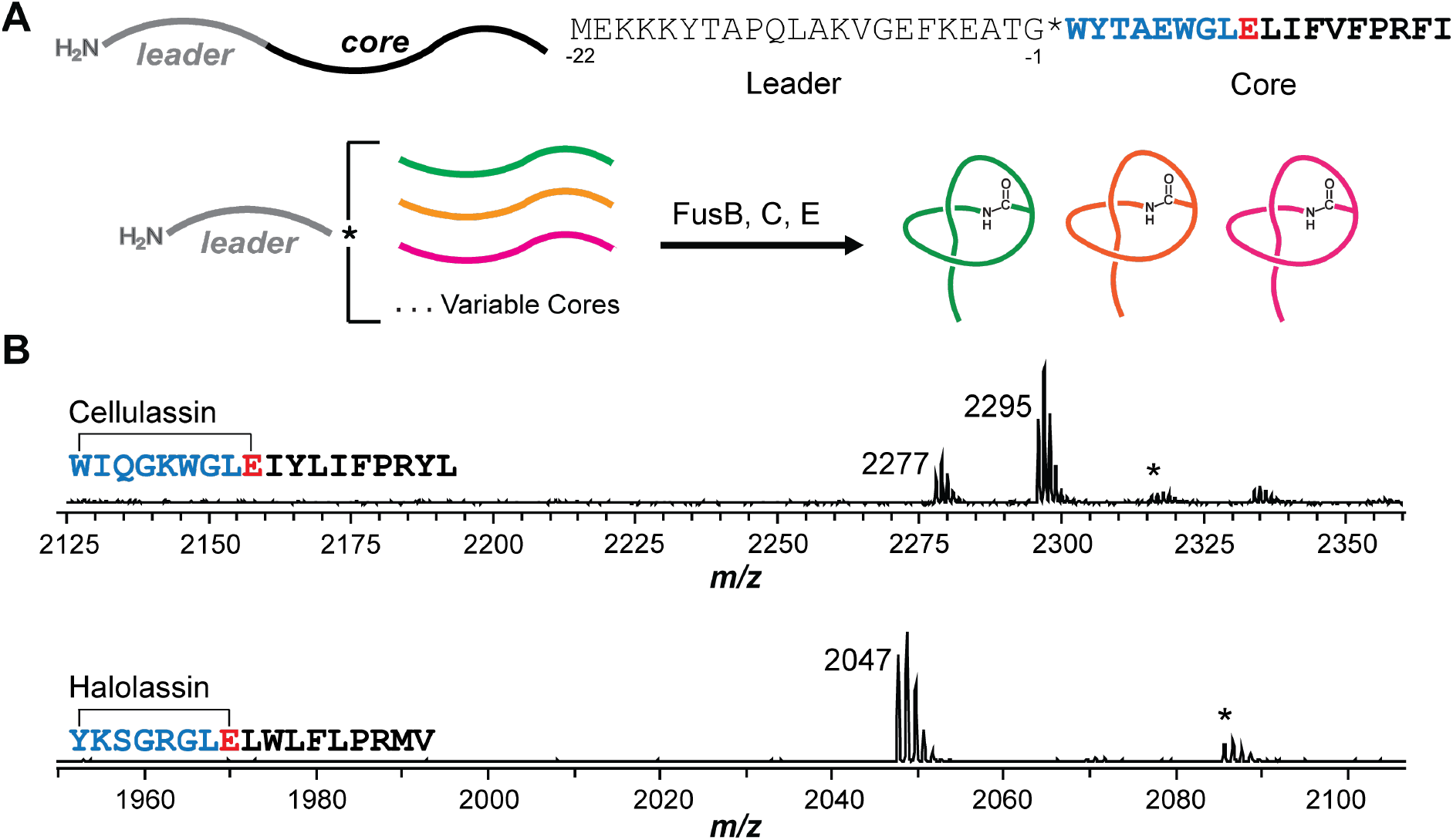
Chimeric substrate strategy to generate diverse lasso peptides. **(A)** The chimeric substrate principle fuses diverse (non-cognate) core peptides to the leader peptide sequence of FusA (residues −1 to −22). This permits a single set of biosynthetic proteins (FusB, C, and E) to produce various lasso peptides. **(B)** Endpoint MALDI-TOF-MS assay of CFB reactions with FusA_LP_-CelA_CP_ (cellulassin, 2277 Da; uncyclized core 2295 Da) and FusA_LP_-HalA_CP_ (halolassin, 2047 Da) as the chimeric substrates. Blue, macrolactam ring residues; Red, acceptor site. The mass label corresponds to the [M+H]^+^ ion of the lasso peptide. * indicates the [M+K]^+^ ion for the lasso peptide.

For the eight chimeric substrates that failed to be processed by the fusilassin biosynthetic proteins, we sought to determine whether a lack of cyclization by FusC or leader peptidolysis by FusB/FusE was responsible. Due to the presence of endogenous proteases in the *E. coli* cell extract, unprocessed precursor peptides and uncyclized, linear core peptides were expected to undergo proteolytic degradation. To monitor linear core peptide formation, we used the PURExpress *in vitro* Protein Synthesis Kit (NEB), which provides the minimal components necessary for transcription and translation (i.e., endogenous proteases are absent).^57^ As a positive control, PURExpress reactions were first carried out to produce the FusA and FusA_LP_-HalA_CP_ precursor peptides. Purified FusB, FusE, FusC, and ATP were then added prior to analysis by MALDI-TOF-MS (**Supplemental Methods**). Fusilassin and halolassin were readily detected, although a significant amount of linear core (uncyclized) was observed in the halolassin sample (**Figure S14**). These data suggest that the fusilassin biosynthetic proteins tolerate HalA_CP_, although the processing efficiency is lower relative to the native substrate. These results demonstrate that the endogenous proteases present in the *E. coli* extract will degrade short, uncyclized peptides, as the linear core of HalA was not detected during standard CFB (**Figure 2**). With the PURExpress system validated, we next tested the eight chimeric precursor peptides that were undetected using the earlier CFB procedure. After precursor peptide production by PURExpress, exogenous addition of biosynthetic proteins, and MALDI-TOF-MS analysis, linear core peptide was detected for 8/8 chimeric substrates and no evidence of cyclization was observed (**Figure S14**). Therefore, FusB and FusE properly removed FusA_LP_ from the chimeric substrates but FusC did not tolerate the alternative core sequences.

The core region of the two successful chimeric substrates (HalA_CP_ and CelA_CP_) encode FusA-like hydrophobic residues in the loop region (**Table S1**). In contrast, the ring regions of these substrates are more variable, suggesting that FusC might be more tolerant of sequence variation in the ring relative to the loop. To test this hypothesis, the loop regions of four failed chimeric substrates derived from *Marinactinospora thermotolerans*, *Nonomuraea candida*, *Streptomyces mobaraensis*, and *Streptomyces* sp. NBS 14/10 were replaced with the FusA loop sequence (_10_LIFVFP_15_, **Figure S15**). The latter two hybrid lasso peptides (i.e., FusA_LP_-MobA_CP_:LIFVFP and FusA_LP_-NbsA_CP_:LIFVFP) were successfully produced by CFB using exogenously supplied fusilassin biosynthetic proteins, suggesting that a native-like loop sequence was more critical than a native-like ring for FusC substrate tolerance. To evaluate the role of the ring in the successful maturation of these chimeric substrates, an additional four hybrid precursor peptides were derived from the same strains. While these sequences retained their native (non-FusA) loop sequences, the ring regions were replaced with the FusA sequence (_2_YTAEWG_7_). No cyclized product was observed from any of these hybrid precursor peptides, in agreement that a native-like loop region was critical for FusC substrate tolerance (**Figure S15**).

### Evaluating FusC substrate tolerance through Ala-substitution

The core sequences of cellulassin and halolassin share the highest identity to fusilassin (8/18 and 6/18 residues, respectively). Taken together with citrulassin A (3/18 allowing for a gap and non-wildtype replacement of Asp8 with Glu, **Table S1**),^6^ the available data demonstrate that FusC can accept certain non-cognate chimeric substrates that differ considerably from FusA. To determine whether any single residue was necessary for cyclization by FusC, we prepared 16 Ala variants of FusA that individually replaced each core residue, except for Ala4 and the Glu9 macrolactam acceptor residue. Given that maxi-prep plasmid preparation is laborious, we developed a time- and cost-efficient method to obtain DNA templates for CFB. Instead of using plasmids, we PCR-amplified the linear DNA products comprising the T7 promoter, lac operon, *fusA* mutant, and T7 terminator for use as the precursor template (**Figure S16**). To prevent the degradation of linear DNA, CFB reactions were supplemented with the nuclease inhibitor GamS (NCBI accession CAA23978.1, **Figure S10)**.^43^ Similar to the chimeric strategy illustrated above, FusB, FusC, and FusE were supplied as purified proteins. The convenience of this optimized CFB strategy was demonstrated by the production of wildtype fusilassin and every Ala-substituted variant except for FusA-R16A (**Figure 3**). The lack of detection of FusA-R16A is reminiscent of our previous observation that conversion of FusA-R16G into the corresponding fusilassin variant by in vitro reconstitution using all purified components was significantly impaired.^6^ We attribute the lack of detection by CFB to inefficient cyclization of FusA-R16A with concomitant degradation by endogenous proteases. Furthermore, this variant removes the only basic residue in the peptide, making any potential product more difficult to detect by positive mode MALDI-TOF-MS.^58^ To evaluate if FusC could tolerate the R16A variant, we created a double variant that introduced Arg into the ring. The W6R/R16A variant of fusilassin was readily detected after CFB-based production, showing that Glu9 is the only position of fusilassin intolerant of Ala-substitution (**Figure 3**). Therefore, when assessing substrate tolerance by CFB, a number of factors must be considered, including the limit of detection and the relative rates of lasso cyclization and proteolytic degradation. To probe more deeply into the substrate tolerance of FusC, a more comprehensive analysis with multi-site replacement was thus warranted.

**Figure 3.**
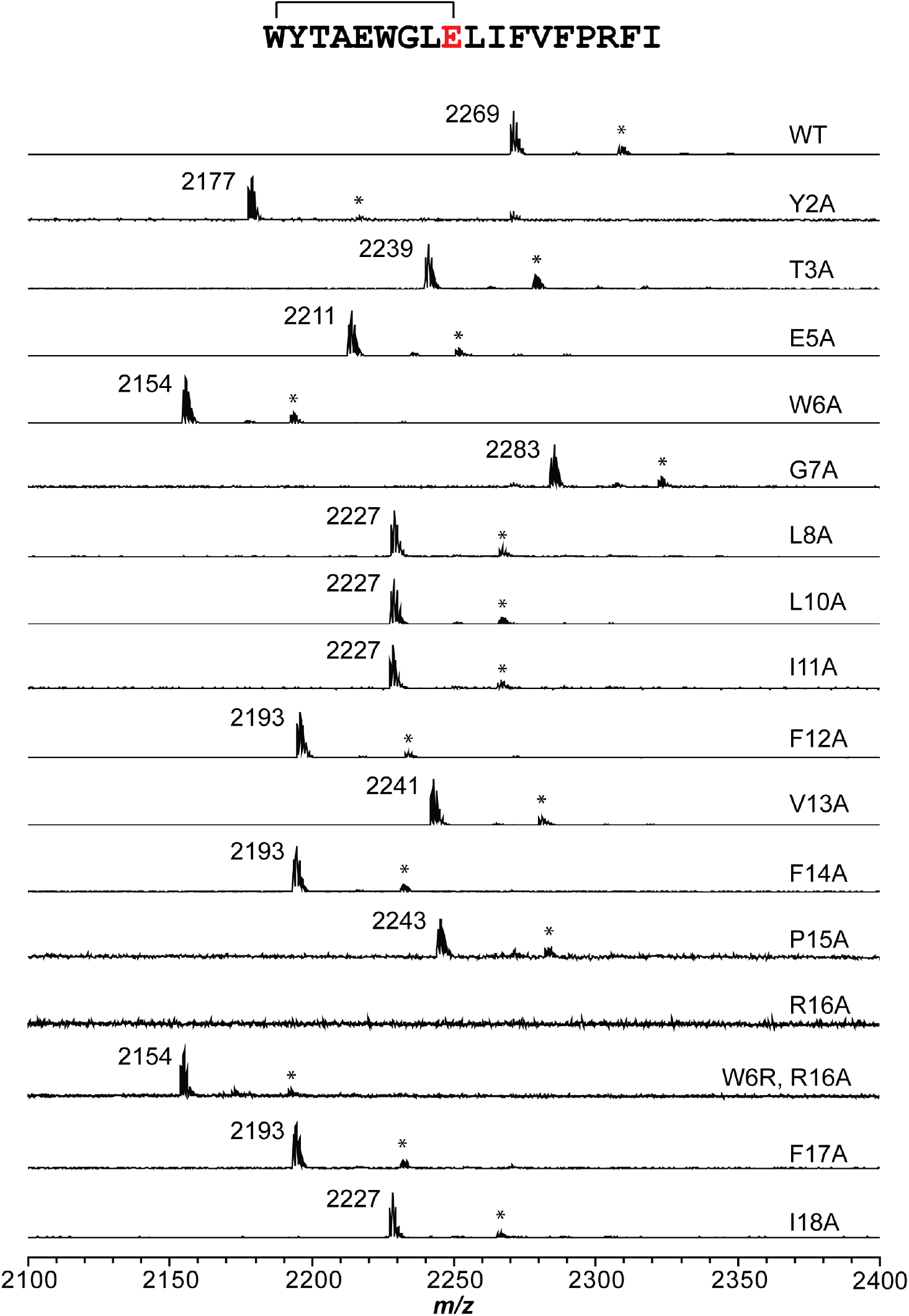
Evaluating FusC substrate tolerance through Ala-substitution. The sequence of wildtype fusilassin is provided at the top with endpoint MALDI-TOF-MS assay data for CFB-produced fusilassin variants shown below. The Glu9 acceptor site is intolerant to substitution.^6^ The mass label denotes the lasso peptide [M+H]^+^ ion. * indicates the lasso peptide [M+K]^+^ ion.

### CFB to probe lasso peptide substrate tolerance through multisite replacement

Based on the results of the chimeric substrate study, the data suggested that FusC is more tolerant to variation in the ring region compared to the loop region. To investigate the tolerance of FusC in more depth, we prepared two libraries that replaced five contiguous residues in these regions (ring positions 2–6 and loop positions 10–14, **Table S3**). These positions avoid altering the macrolactam-forming and steric plug residues. Each library was prepared using the degenerate NNK codon (comprises 32 codons that provides all 20 common amino acids and one stop codon) for a theoretical size of 3.2 million variants. In this design, ~16% of the sequences will contain a premature stop codon in the varied region.^59^

The ring and loop libraries were cloned into pET28, and multiple transformations were performed with chemically competent *E. coli*. The transformed cells were then diluted and plated to obtain ~1,000 colonies per 10 cm plate with each colony presumed to encode a unique FusA variant. Colony PCR was performed on individual colonies in 96-well plate format to amplify linear DNA. After ethanol precipitation, the purified linear DNA was used in CFB reactions as before (**Figures 4–5**). The expected masses of the FusA variants range from 1923−2569 Da (ring library) and 1954–2599 Da (loop library) with the lowest and highest masses possessing Gly_5_ and Trp_5_ variable regions, respectively. A control CFB reaction lacking template DNA was conducted and confirmed that this mass range was clear of any potentially confounding entities (**Figure S17**). Owing to endogenous proteases present in the CFB extract, we anticipated that the majority of detected masses would correspond to FusC-tolerated cyclic products, although we also expected to occasionally detect linear, uncyclized core peptides (i.e., processed by the leader peptidase but not the lasso cyclase)

**Figure 4.**
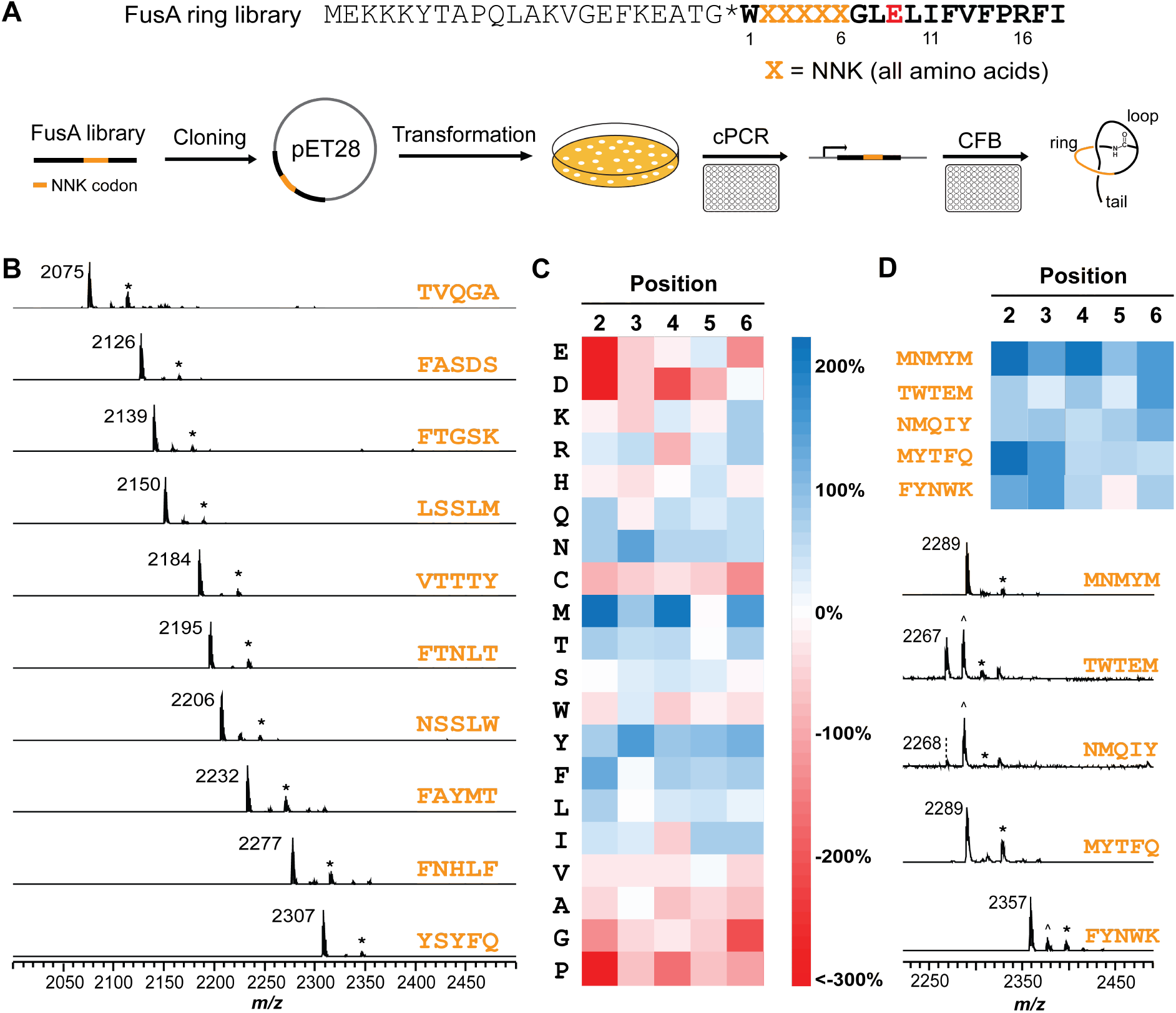
Substrate compatibility evaluation of FusA ring. **(A)** Experimental workflow for fusilassin ring library screening. Positions 2-6 of the FusA core region were diversified using 5 NNK codons (orange, library size of 3.2 M unique peptide sequences). After library cloning and transformation into *E. coli*, colony PCR was performed in 96-well plates. The linear DNA products were then used in CFB reactions to produce lasso peptides. **(B)** MALDI-TOF mass spectra of 10 representative fusilassin ring variants. The sequences of the varied region are orange. **(C)** Heatmap analysis on confirmed FusC substrates (lasso-compatible sequences) showing the percent difference between the occurrence of a residue by core position and the expected frequency based on the NNK codon. The analysis was conducted from 280 substrates. **(D)** MALDI-TOF mass spectra showing the lasso peptide formation of five predicted FusC substrates from the heatmap. At least four residues in designed sequences are predicted to be favored in FusC substrates. The precursor peptides were synthesized through CFB and reacted with heterologously expressed and purified FusB, C, and E. The mass labels denote the [M+H]^+^ ion of the lasso peptides. ^ indicates the uncyclized linear core peptide. * indicates the [M+K]^+^ ion for the lasso peptide.

Individual colonies from the ring and loop NNK libraries were subjected to the above-described workflow and subsequent MALDI-TOF-MS analysis. Ions within the expected mass range were detected for 513/984 (~52%) ring variants and 3/57 (~5%) loop variants. All three successful loop variants were sequenced, confirming the observed mass corresponds to the cyclized lasso peptide, instead of the linear core peptide or other truncated precursor peptides. Among the three fusilassin loop variants, no polar or charged residues were present, despite a high probability of occurrence from five NNK codons (**Figure 5**).

**Figure 5.**
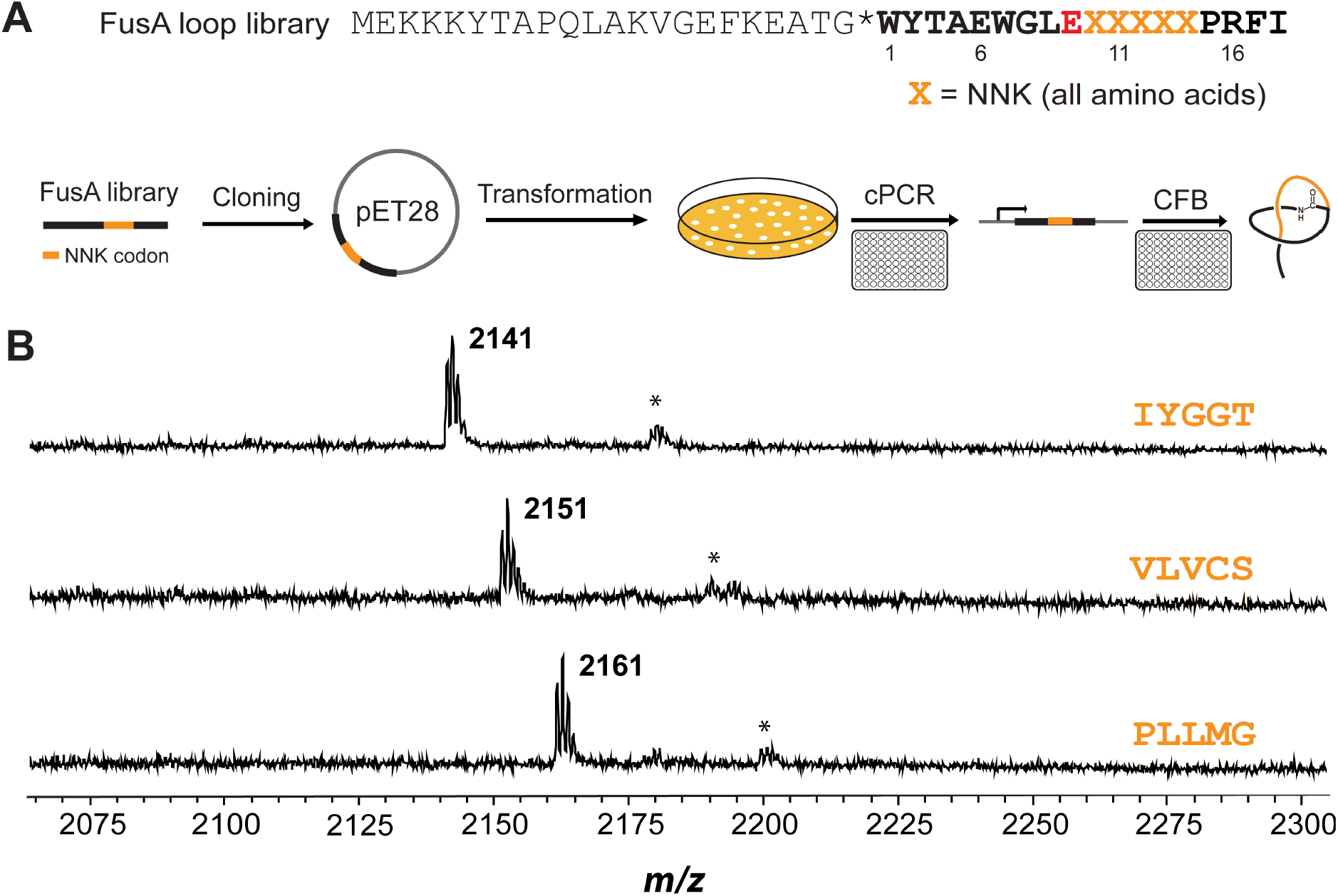
Substrate compatibility evaluation of FusA loop. **(A)** Experimental workflow for fusilassin loop library evaluation. Positions 10-14 of the FusA core region were diversified using 5 contiguous NNK codons (orange, theoretical library size of 3.2 M unique peptide sequences). After library cloning and transformation into *E. coli*, colony PCR (cPCR) was performed in 96-well plates. The linear DNA products were then used in CFB reactions to produce lasso peptides. **(B)** From 57 randomly chosen clones, lasso peptide formation was only detected for three loop variants. Orange depicts the varied sequence at positions 10-14 of the FusA loop region. The mass labels denote to the [M+H]^+^ ion of the lasso peptide. * indicates [M+K]^+^ ion for the lasso peptide.

Given the higher success rate, we carried out a more thorough analysis on the ring library. Adjusting for the expected frequency of stop codons within the library, the probability of a random ring sequence (core positions 2–6) being tolerated by FusC increases to ~61%. Therefore, the fusilassin biosynthetic proteins can generate nearly 2 million distinct lasso peptides from a penta-substituted ring library. Any future lasso peptide library that is restricted to the ring region still represents a large fraction of the exposed surface area (**Table S4**), indicating the ring region of lasso peptides could be engineered to display customized binding activities. Among the 513 sequences expected to be converted to mature lasso peptides, we sequenced the DNA from more than half (*n* = 280), confirming that the observed mass precisely corresponded to the expected peptide sequence (**Figures 4, S18**). In a minority of cases, masses corresponding to the linear core peptides (FusB/E product) were detected, which likely arose from slower rates of lasso peptide formation and proteolytic degradation. The available data support a sequence-dependent rate of lasso peptide formation, which will be the subject of future study.

Ten successfully produced members of the ring library were selected for structural and stability assessment. HRMS/MS analysis supported a Trp1-Glu9 ring composition for each variant (**Figure S19-28**). Nine out of ten variants were resistant to carboxypeptidase Y digestion, suggesting a threaded topology (**Figure S29**). The LSSLM ring variant was susceptible to carboxypeptidase Y even prior to heating, suggesting this lasso peptide was especially prone to unfolding. Among the nine carboxypeptidase Y-resistant variants, a mass consistent with removal of Ile-18 was observed at a ratio that varied dramatically with the mass intensity of the full-length lasso peptide. Also, a range of heat stabilities was noted, which illustrates that the stability of the lasso fold is sequence-dependent (**Figure S29**).

A sequence logo, generated from the 280 sequence-confirmed FusC substrates, showed little to no conservation at any of the five varied ring positions, indicating broad tolerance at each position (**Figure S30**).^60^ We further analyzed the preference for specific amino acids by comparing the observed frequency at each position with the expected frequency based on the NNK codon (**Figure 4**). Overall, FusC displays a modest preference at positions 2, 4, and 6 compared to positions 3 and 5. There is a subtle bias towards hydrophobic residues at the even-numbered positions, such as Met, Phe, and Tyr, while charged residues like Glu, Asp, and Lys were disfavored. These results indicate that the FusC binding pocket for FusA is likely to be hydrophobic, and with the side chains of residues 2, 4, and 6 likely orienting on one face of the ring (odd-numbered residues oriented in the opposite direction), even-numbered ring positions may be more likely engaged by the enzyme. Several other residues show site-specific preferences. For example, Glu is favored at position 5 over all other positions while Lys is favored at position 6. To determine whether we could predict FusC tolerated sequences, we designed five predicted substrates according to the heatmap (core positions 2–6 varied with: MNMYM, TWTEM, NMQIY, MYTFQ, and FYNWK). Our predictions were correct for 5/5 substrates by MALDI-TOF-MS (**Figure 4**). Further, these FusA variants were readily produced from a single CFB reaction (**Figure S31**).

We next conducted a parallel analysis for FusC non-substrate sequences. We randomly sequenced 154 non-substrates and confirmed the formation of linear core peptide using PURExpress (**Figure S32**), indicating that FusB/E universally processed the leader region of all FusA variants. We then analyzed the preference at each position and generated a second heatmap (**Figure S33**). Analogous to the observations on true substrates, the presence of charged residues (e.g., Glu, Lys, Arg) were enriched in the non-substrate set. To test whether our analysis could predict FusC non-substrates, we designed an additional five FusA ring variants (IKEVT, QDWFM, FLRCL, IDRSY, and LKNFT). Our predictions were correct for 5/5 non-substrates. Similarly, PURExpress reactions confirmed FusB/E produced linear core in every case but FusC failed to cyclize these peptides (**Figure S33**).

### Bioinformatics predicts extensive diversity of naturally occurring lasso peptides

Given the unprecedented substrate tolerance in the ring region, we sought to explore whether wide sequence variability of the lasso peptide ring is a characteristic of known and predicted lasso peptides. We first conducted a survey to update the list of all lasso peptides predicted from the NCBI database. This latest dataset used 11 known lasso leader peptidases as queries for Position-Specific Iterative (PSI)-BLAST (**Table S5**),^61^ with each class and major phylum known to produce lasso peptides represented. Proteins homologous to the leader peptidase (*n* = 9,571, as of Nov-2020) were subjected to RODEO analysis.^11^ This yielded 7,701 non-redundant, high-scoring lasso precursor peptides, and more than doubles the number previously reported.^6^ Each entry was verified to locally encode the requisite RRE, leader peptidase, and lasso cyclase (**Supplemental Dataset 1**).

Among the 7,701 lasso precursor peptides, 4,485 unique core sequences remained after removal of identical entries (if two distinct organisms encode the same core sequence, only one was retained). To describe the residue variability at each core position, the Shannon entropy,^62^ relative entropy,^63^ and ConSurf values^64^ were calculated (**Table S6**). To ascertain the amino acid specificity among naturally encoded lasso peptides, we compared the observed, position-dependent frequency of each amino acid to the expected frequency. The latter was calculated from the weighted average of codon usage at the genus-level (**Table S7, Figure S34**). For the set of predicted, 9-residue ring (same size as fusilassin) lasso peptides core sequences with all replicate entries removed, the Shannon entropy value at each position, was >3 (values >2 are considered variable). The only exception was the macrolactam acceptor site, which gave a Shannon entropy value <1. Within the ring region, positions 3−7 show the highest variability and the propensity to observe a particular amino acid in this region is near the expected frequency. Core position 7 was the most variable by all three metrics (Shannon and relative entropies and ConSurf score). This analysis further shows that, excluding the macrolactam-forming positions 1 and 9, the ring variability of natural lasso peptides is slightly higher than the loop-tail regions (**Table S6)**. Notably, there are enrichments of specific amino acids, such as Gly/Ser within the ring, but not in the loop-tail region. Phe/Val are significantly enriched at position 11 while Gln/Phe are overrepresented at position 12 (**Figure S34**). Extensive sequence diversity is evident across all regions of natural lasso peptides; however, certain locations appear to be under evolutionary selection. This result could have been anticipated, as not every possible peptide sequence should be compatible with lasso peptide formation, nor will every possible sequence perform a beneficial function. The naturally encoded sequence space observed today is thus logically confined by substrate-enzyme co-evolution and conditional selection pressure that defines the breadth of the biological activities exhibited by lasso peptides in nature.

### Summary and Conclusion

The data presented herein demonstrate the first application of cell-free biosynthesis (CFB) for the production of lasso peptides and shows the versatility of this method for enhancing understanding of biosynthetic tolerance. We first produced four known lasso peptides, capistruin, burhizin, fusilassin, and cellulassin using CFB. Capistruin was used as an example to demonstrate that the CFB reactions are scalable, as we produced 200-fold more capistruin by CFB relative to heterologous expression in live *E. coli*. Capistruin was then subjected to a CFB-based transcription/translation inhibition assay that monitored the production of the fluorescent protein mCherry. Considering the speed, ease of manipulation, and serviceable yields achieved for lasso peptides, CFB provides a new route to overcome production issues often encountered when attempting to isolate mature products from the native producer or through heterologous expression.

This work further advances the chimeric strategy of RiPP production by fusing variable core regions to a native leader peptide to permit substrate recognition by the employed biosynthetic proteins. We produced a new, genomically predicted lasso peptide by this method in addition to confirming wide substrate tolerance for the fusilassin biosynthetic proteins. The efficiency of CFB was leveraged to investigate the substrate scope of the fusilassin pathway by screening a diverse, penta-substituted NNK library of FusA variants. Over 1,000 FusA variants were rapidly assayed, showing FusC is widely permissive of ring substitution but less tolerant in the loop region. Our analysis of natural lasso peptide core sequences suggests that other lasso peptide pathways may display a wider substrate tolerability in the loop region. Thus, when used together with the fusilassin biosynthetic proteins, library coverage of a large majority of the exposed surface area of a lasso peptide could be achievable.

With less than 2% of unique lasso peptide sequences being characterized to date, this molecular class of natural product remains critically underexplored. CFB is a promising tool to access these molecules, particularly from difficult to cultivate or otherwise inaccessible organisms. Our data also provide a solid foundation to eventually permit the customization of lasso peptides for specific biotechnological applications.

## Acknowledgment

We thank Julian Hegemann and Adam J. DiCaprio for providing expression constructs and Xiao Rui Guo for assistance with acquiring high-resolution mass spectral data (Univ. of Illinois at Urbana-Champaign). We thank Jozsef Kukolya (Agro-Environmental Research Institute) for providing *Thermobifida cellulosilytica* and Alessandra Eustáquio (Univ. of Illinois at Chicago) for providing capistruin-containing culture extracts. This work was supported in part by the Ving and May Lee Chemistry Discovery Fund (Department of Chemistry, Univ. of Illinois at Urbana-Champaign) and the National Institutes of Health (R41 AT010852 to M.J.B. and D.A.M.).

## Author Contributions

Y.S., A.M.K., and L.M.D. performed the experiments. All authors designed experiments, analyzed data, and assisted in the writing and editorial process. D.A.M. and M.J.B. conceived of and supervised the project.

---

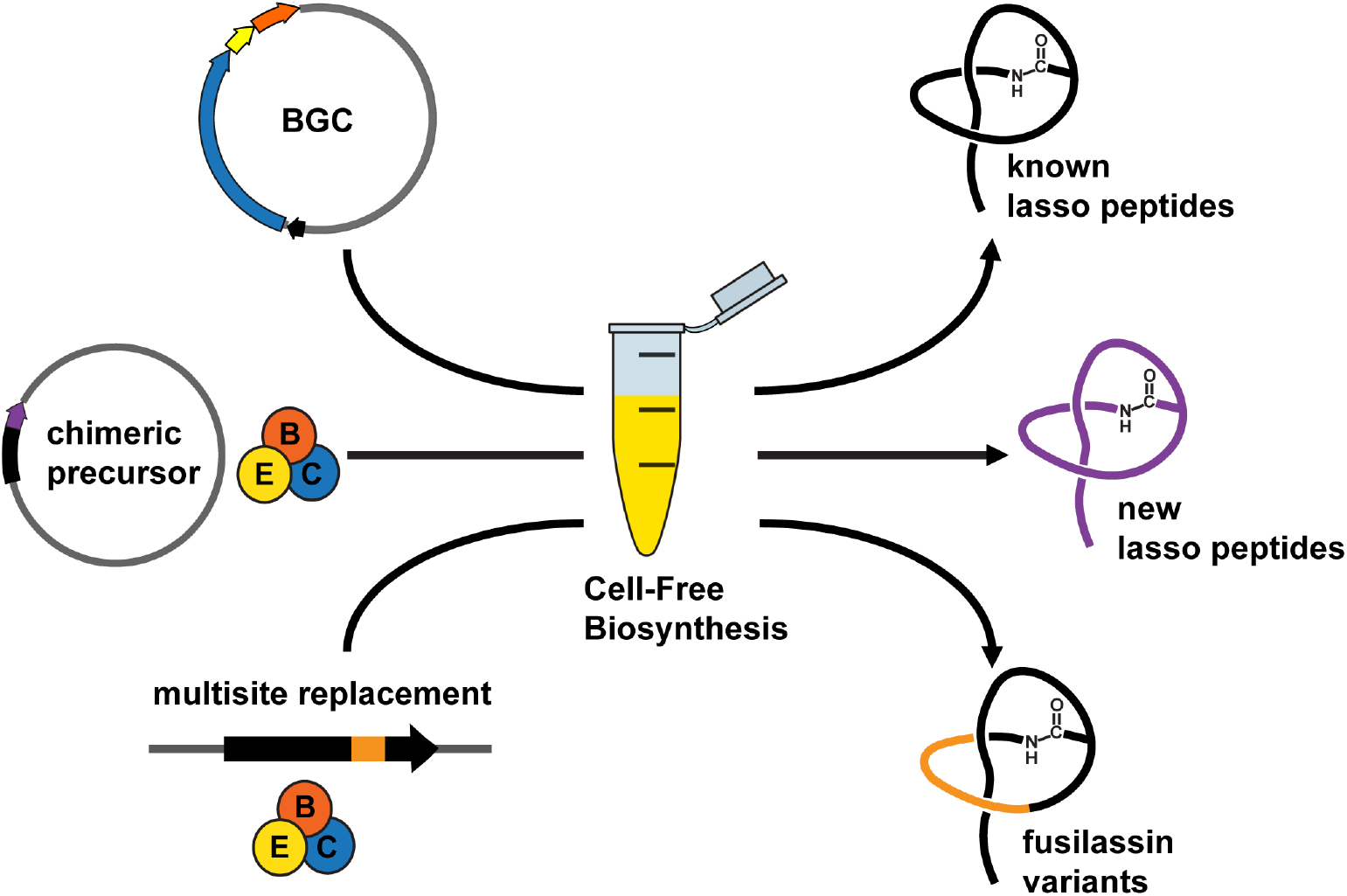

