## Supplementary material for "Cell-Free Biosynthesis to Evaluate Lasso Peptide Formation and Enzyme-Substrate Tolerance": SI document

### Materials and Methods

**General materials and methods.** The synthetic DNA encoding FusA chimeric precursor peptides were codon optimized for *E. coli* expression and were purchased from Twist Bioscience (**Table S2**). The synthetic DNA containing NNK degenerate codons (**Table S3**) and primers used in this study (**Table S8**) were purchased from Integrated DNA Technologies. Q5 DNA polymerase, restriction enzymes, T4 DNA ligase, and high efficiency *Escherichia coli* (*E. coli*) DH5α competent cells were obtained from New England Biolabs. Amicon Ultra-4 centrifugal filters were purchased from EMD Millipore. *E. coli* DH5α and *E. coli* BL21(DE3) cells were used for plasmid maintenance and protein production, respectively. Chemical reagents were from Sigma Aldrich and used without further purification unless otherwise noted. Sanger sequencing was performed by the Core DNA Sequencing Facility at the University of Illinois at Urbana-Champaign. Matrix-assisted laser desorption/ionization time-of-flight mass spectrometry (MALDI-TOF-MS) data were acquired in reflector positive mode on a Bruker UltrafleXtreme instrument (Bruker Daltonics) at the University of Illinois School of Chemical Sciences Mass Spectrometry Laboratory. MALDI-TOF-MS data analysis was carried out using the Bruker FlexAnalysis software.

**Molecular biology techniques.** The chimeric precursor peptides with different core sequences fused to the FusA leader region were cloned into a modified pET28 vector that provides an N-terminal maltose-binding protein (MBP) tag using 5' BamHI and 3' HindIII restriction sites. Predicted FusC substrates (core positions 2-6 varied with: MNMYM, TWTEM, NMQIY, MYTFQ, and FYNWK), predicted FusC non-substrates (IKEVT, QDWFM, FLRCL, IDRSY, and LKNFT), hybrid precursor peptides (i.e. $FusA_{LP}$-$MobA_{CP}$:LIFVFP and $FusA_{LP}$-$NbsA_{CP}$:LIFVFP) were prepared through PCR overlap extension.[1] MBP-tagged fusilassin leader peptidase (WP_011291590.1, FusB), lasso cyclase (WP_104612995.1, FusC), and RiPP leader peptide recognition element (RRE) (WP_011291591.1, FusE) were cloned previously into pET28-MBP.[2] The obtained constructs were used for heterologous expression, and pET28-MBP-FusA was used as the template for mutagenesis. Mutagenesis was performed using the QuikChange method with Q5 Polymerase. mCherry was amplified from pmCherry (Clontech) and cloned into pET28 (no MBP tag) using 5' BamHI and 3' HindIII restriction sites. The obtained construct was used to determine transcription inhibition by capistruin. All mutants were verified by Sanger sequencing.

***E. coli* cell lysate preparation.** A single BL21(DE3) colony was inoculated into 10 mL of YTPG medium (10 g/L yeast extract, 16 g/L tryptone, 3g/L $KH_2PO_4$, 7g/L $K_2HPO_4$, 5 g/L NaCl, 18 g/L glucose) and grown at 37 °C overnight with shaking. The culture was then used to inoculate 1 L of YPTG medium and grown at 37 °C with shaking until an optical density ($OD_{600}$) reaches ~0.5, at which point 1 mM IPTG was added. When an $OD_{600}$ of ~3 was observed, the cells were harvested by centrifugation at 4,000 × *g* for 20 min at 4 °C. The cells were washed twice with 2 × 400 mL of cold buffer S30A (10 mM Tris, pH = 8.2, 14 mM magnesium acetate, 60 mM potassium glutamate, 2 mM 1,4-dithiothreitol). After the second wash, the cell pellet was resuspended in ~40 mL of cold buffer S30A in a pre-weighed 50 mL conical tube, followed by centrifugation at 5,000 × *g* for 10 min at 4 °C. The cell pellet was then weighted and resuspended in buffer S30A at 1 mL/g cell. The cells were lysed with a homogenizer (single pass, 20,000-30,0000 psi), then subjected to centrifugation at 30,000 × *g* for 30 minutes at 4 °C.

A Pierce Bradford assay was used to determine the total protein concentration in the cell extract (~38 mg/mL). The cell extract was aliquoted and stored at -80 °C until needed.

**Energy buffer preparation.** The energy buffer was prepared following a previously described protocol.[3]

**DNA template preparation for cell-free biosynthesis (CFB) reactions.** Circular plasmids used in CFB were purified with maxi-prep. To be specific, *E. coli* DH5α chemically competent cells were transformed with reconstituted plasmid, and the cells were grown overnight on Luria-Bertani (LB) agar plates containing 50 µg/mL kanamycin at 37 °C. Single colonies were picked to inoculate 200 mL of LB medium containing 50 µg/mL kanamycin and grown at 37 °C for 16–18 h with shaking. Cells were harvested via centrifugation at 3,800 × *g* for 20 min at 4 °C. The plasmids were isolated and purified using a Qiagen plasmid maxi kit per the manufacturer's instructions. The concentration of purified plasmid was determined by measuring the absorbance at 260 nm using a Nanodrop One$^{C}$ instrument (Thermo Scientific).

Linear DNA templates used in CFB reactions were amplified through polymerase chain reaction (PCR) from the recombinant pET28 plasmids using primers 44 and 45 (**Table S8**). The resulting PCR products were purified with Qiagen PCR clean up kit per the manufacturer's instructions. For the linear DNA templates used in library screening, the PCR products were purified with ethanol precipitation (details are described below). The concentrations of purified plasmids were determined using the Nanodrop as above.

**Capistruin, burhizin, fusilassin, and cellulassin production by CFB.** Reactions were carried out at a total volume of 21 µL, including: 6.9 µL *E. coli* cell extract, 8.8 µL energy buffer, 0.21 µL isopropyl β-D-1-thiogalactopyranoside (IPTG, 50 mM stock concentration) and 0.63 µL 1,4-dithiothreitol (DTT, 100 mM stock concentration). Circular plasmids (20 nM) encoding the biosynthetic gene clusters (BGCs) of capistruin, burhizin, fusilassin (both pET28-MBP-*fusA* and pACYC-*fusBE*-*fusC* were added at 20 nM), and cellulassin were used to initiate the CFB reactions. The reactions proceeded at room temperature for 16 h, then were desalted with ZipTip $C_{18}$ (Sigma-Aldrich) and eluted with 60% acetonitrile (ACN). The eluents were analyzed by MALDI-TOF-MS using α-cyano-4-hydroxycinnamic acid (CHCA) as the matrix.

**High-resolution mass spectrometry and tandem mass spectrometry (HRMS/MS) of lasso peptides.** The CFB reactions producing desired lasso peptides were desalted using ZipTip $C_{18}$ and eluted into 60% aq. ACN. The eluents were directly infused into a ThermoFisher Scientific Orbitrap Fusion ESI-MS using an Advion TriVersa Nanomate 100. The MS was calibrated and tuned with Pierce LTQ Velos ESI Positive Ion Calibration Solution (ThermoFisher). The MS was operated using the following parameters: resolution, 100,000; isolation width (MS/MS), 1 *m/z*; normalized collision energy (MS/MS), 70; activation q value (MS/MS), 0.4; activation time (MS/MS), 30 ms. Fragmentation was performed using collision-induced dissociation (CID) at 35 to 70%. The resulting data were averaged and analyzed using the Qualbrowser application of Xcalibur software (Thermo-Fisher Scientific).

**Carboxypeptidase Y evaluation of threadedness.** The CFB reactions were treated with 60% ACN. Debris from the quenched reactions was removed by centrifugation at 8,000 × *g* for 2 min at room temperature. The supernatant was evaporated to dryness under vacuum and resuspended in 50 ng/µL carboxypeptidase Y in phosphate-buffer saline (PBS). This mixture was briefly mixed by vortex and allowed to react at room temperature for 18 h. Resulting peptidase mixtures were treated with 60% acetonitrile and analyzed by MALDI-TOF-MS with sinapinic acid (SA) as the matrix. For thermal unthreading assays, supernatant obtained after 60% ACN treatment were incubated at 95 °C for 2 h. This product was then evaporated to dryness under vacuum and resuspended in 50 ng/µL carboxypeptidase Y in PBS and reacted at room temperature for 18 h. The digested product was analyzed with MALDI-TOF-MS as above.

**Purification of the capistruin standard.** A methanol extract of spent media of *Burkholderia* sp. FERM BP-3421 cultures expressing the capABCD[4] was provided by Alessandra Eustáquio (Univ. of Illinois, Chicago). Capistruin was purified from the extract by high-performance liquid chromatography (HPLC, Agilent 1200 series). Extract was injected onto a C18 column (Thermo Scientific betasil 250 × 10 mm 5 µM particle size). Capistruin was separated using mobile phase A (water, 0.1% TFA) and mobile phase B (acetonitrile, 0.1% TFA) over a gradient of 4 minutes at 10% B, 2 min 10−32.5% B, 8 min 32.5−35% B, and 2 min 35-95%, 4 min 95%, at a flow rate of 5 mL/min. UV-vis absorbance from 190−450 nm was recorded. Fractions corresponding to UV absorbance peaks (220nm) were collected and tested for the presence of capistruin by MALDI-TOF-MS, using CHCA as the matrix. Capistruin eluted at 14 min; this fraction was concentrated to dryness on the lyophilizer (Labcono FreeZone 6L). The purification was repeated to obtain a >95% pure sample of capistruin. A sample of the purified capistruin was resuspended in water at a concentration of 5 mM, considered the "standard" sample for our quantification of capistruin by CFB.

**Purification and yield determination of capistruin from CFB.** Reactions were carried out at a total volume of 750 µL including: 262.5 µL *E. coli* cell extract, 337.5 µL energy buffer, 24 µL DTT (100 mM stock conc.), 9 µL IPTG (50 mM stock conc.). The PCR product from pET41-capABCD (1400 ng, primers 44 and 45 **Table S8**), including the T7 promoter, capABCD, and terminator, was added to initiate the reaction. The reaction was separated into 8 tubes for higher aeration, and incubated at 37 °C for 4 h, followed by 28 °C for 20 h with shaking at 1500 rpm. ACN was added to a final concentration of 50% to precipitate salts and proteins. The reaction was centrifuged at 10,000 × *g* for 10 minutes at room temperature to remove precipitate and the supernatant was concentrated on a lyophilizer.

A standard curve was determined by injections of 3, 7.5, 10, 15, 20 and 30 nmol (approx. 6, 7.5, 10, 15, 20, 30, and 60 µg) of the standard sample of capistruin onto the HPLC, under the same conditions. The area of the 220nm absorbance peak corresponding to capistruin was calculated for each concentration and plotted together to establish the standard curve ($\chi^2 = 0.99$).

Capistruin was purified from the CFB reaction by HPLC in the same manner as the standard sample. The lyophilized CFB reaction was resuspended in 500 µL 50% ACN and purified in a single injection. Fractions corresponding to UV absorbance peaks (220nm) were collected and tested for the presence of capistruin by MALDI-TOF-MS, using CHCA as the matrix. Capistruin eluted at 14 min, identical retention time to the standard capistruin sample. To determine the yield of capistruin from CFB, the area of the 220nm absorbance peak was compared to the standard curve. Purified CFB-produced capistruin was concentrated to dryness on the lyophilizer and stored at -20 °C until needed.

**Validation the identity of capistruin purified from CFB by liquid chromatography-mass spectrometry.** Capistruin from the CFB reaction was compared to the standard sample by liquid chromatography-mass spectrometry (LC-MS, Shimadzu LCMS 2020). A sample of the purified capistruin from the CFB reaction was injected onto a C18 column (Macherey Nagel EC Nucleodur 250 × 4.6 mm 5 μM particle size, 100 μL injections), using mobile phase A (water, 0.1% FA) and mobile phase B (ACN, 0.1% FA) over a gradient of 5 minutes at 10% B, 5 min 10−35% B, 15 min 35−65% B, and 5 min 65-95%, 4 min 95%, at a flow rate of 1 mL/min. UV-vis absorbance from 190−450 nm was recorded and mass at m/z 500-1500 were monitored. The extracted ion chromatograms of 1025.4 m/z $[M+2H]^{2+}$ were compared between capistruin purified from CFB, the standard sample of capistruin, and a co-injection.

**Capistruin-dependent inhibition of mCherry production.** Capistruin, purified from 750 μL of CFB reaction, was resuspended in water to 100 μM. A CFB reaction was set up with 100 ng plasmid encoding mCherry under T7 control with capistruin was added to a final concentration of 0 or 10 μM. The reactions were incubated at 30 °C for 16 h with 225 rpm shaking for aeration. The samples were centrifuged at 15,000 × *g* for 1 min at room temperature to remove precipitate, diluted 1:10 in water, and the fluorescence emission at 600 nm was quantified (Tecan Infinite M2000) using an excitation of 550 nm (n = 2).

**Expression and purification of MBP-tagged FusB, FusE, and FusC.** Previously described methods were employed for the expression and purification of MBP-tagged FusB, FusE, and FusC.[2] Protein concentrations were determined by Bradford colorimetric assay and absorbance at 280 nm.[5] Protein purity was visually inspected by SDS-PAGE using Coomassie staining (**Figure S10**).

**Expression and purification of GamS**. GamS was purified from *E. coli* BL21(DE3) bearing the pBAD-*gamS* plasmid as previously described with minor differences.[6] An overnight culture was used to inoculate 1 L of LB containing 50 μg/mL ampicillin and grown to the $OD_{600}$ of 0.45. Arabinose was added to a final concentration of 0.25% *w/v*, followed by an induction period at 37 °C for 4 h. Cells were harvested via centrifugation at 5,000 × *g* for 30 minutes at 4 °C and resuspended in 30 mL lysis buffer [50 mM Tris pH 8.0, 500 mM NaCl, 5 mM imidazole, 0.1% Triton X-100 (*v/v*), 3 mg/mL lysozyme, 2 μM leupeptin, 2 μM benzamidine HCl, 2 μM E64, and 30 mM phenylmethylsulfonyl fluoride]. Cells were lysed by sonication for ten cycles (30 s each) at 20% amplitude, with 30 s intervals of resting at 4 °C. Insoluble cell debris was removed by centrifugation at 18,000 × *g* for 60 min at 4 °C. The resultant supernatant was loaded onto pre-equilibrated nickel-nitrilotriacetic acid (Ni-NTA) resin (ThermoFisher; 5 mL of resin per L of cells). The column was washed with 30 mL of wash buffer (50 mM Tris pH 8.0, 500 mM NaCl, 25 mM imidazole). The 6xHis-tagged proteins were eluted using 15 mL elution buffer (50 mM Tris pH 8.0, 500 mM NaCl, 250 mM imidazole). The eluent was concentrated using a 30 kDa molecular weight cut-off Amicon Ultra centrifugal filter (EMD Millipore). A buffer exchange with 10 × volume of protein storage buffer [50 mM Tris-Cl pH 7.5, 100 mM NaCl, 1 mM

DTT, 1 mM ethylenediaminetetraacetic acid (EDTA), 2% dimethyl sulfoxide (DMSO)] was performed prior to final concentration and storage. Protein concentrations were determined by Bradford colorimetric assay and absorbance at 280 nm. Protein purity was visually inspected by SDS-PAGE using Coomassie staining (**Figure S10**).

**Chimeric substrates and FusA Ala variants analysis.** The plasmid or linear DNA templates for each mutant and chimeric substrate were prepared as described above and used in a CFB reaction, which was carried out with the total volume of 21 µL including 6.9 µL *E. coli* cell extract, 8.8 µL energy buffer, 0.21 µL IPTG (50 mM initial concentration), 0.63 µL DTT (100 mM stock conc.), 0.5 µL GamS (125 µM stock conc.), 0.5 µL FusB (500 µM stock conc.), 1 µL FusC (200 µM stock conc.), 0.5 µL FusE (500 µM stock conc.) and 2000 ng plasmid DNA (chimeric substrates) or 500 ng linear DNA (FusA ala variants). The reactions proceeded at room temperature for 16 h and were quenched by adding ACN to a final concentration of 60%. Debris from the quenched reactions was removed by centrifugation at 8,000 × *g* for 2 min at room temperature. The supernatant was analyzed by MALDI-TOF-MS using SA as the matrix.

**Linear core peptide detection using the PURExpress system.** For each precursor peptide, a 10 µL reaction was set up using NEB PURExpress kit per the manufacturer's instructions (NEB E6800L) with 2 µL of DNA template added (500 ng for plasmid and 200 ng for linear DNA). The reaction was incubated at 37 °C for 3 h. Each heterologously expressed and purified fusilassin protein (20 µM) was added in 10 µL of synthetase buffer (50 mM Tris-HCl pH 7.5, 125 mM NaCl, 20 mM $MgCl_2$, 10 mM DTT, 5 mM ATP), and then added into the PURExpress reactions and incubated for another 3 h at 37 °C. The reaction was quenched by the addition of ACN directly into the reaction (60% *v/v* final). The debris was removed by centrifugation at 8,000 × *g* for 2 min at room temperature prior to analyzing the supernatant by MALDI-TOF-MS with SA as the matrix.

**FusA degenerate library screening in 96-well format.** DNA containing degenerate segments were synthesized by IDT (**Table S3**) and amplified through PCR using primers 46 and 47 (**Table S8**). A linear section of pET28 vector was amplified through PCR using primers 48 and 49 (**Table S8**), followed by addition of 1 µL DpnI to the reaction and allowed to react at 37 °C for 1 h. The PCR amplicons were purified using a QIAprep PCR Cleanup Spin Kit. The isolated FusA degenerate library and the linear section of pET28 plasmid was then simultaneously digested with KpnI and SacI. Digested insert and vector were then purified with QIAprep PCR Cleanup Spin Kit (Qiagen) and ligated using T4 DNA ligase (NEB). The ligation reactions were directly transformed into high efficiency, chemically competent *E. coli* DH5α (NEB 5-alpha Competent *E. coli*). The cells were plated on 4 LB agar plates (10 cm) containing 50 µg/mL kanamycin. The plates were incubated at 37 °C for 18 h. The PCR mixture was prepared using primers 44 and 45 (**Table S8**) and aliquoted into the 96-well plate with 50 µL per well. Single colonies (n = 96) were picked and added into each well. The PCR products were purified by addition of 100 µL ethanol and 5 µL sodium acetate (3 M, pH 5.2) to each well followed by DNA precipitation at -20 °C for 1 h. After DNA precipitation, the plate was subjected to centrifugation at 4,500 × *g* for 30 min at 4 °C to harvest the DNA product. The supernatant was removed, and the DNA precipitate was resuspended in 70% ethanol for an additional washing step. Supernatant was removed by centrifugation at 4,500 × $g$ for 30 min at 4 °C. The DNA precipitate was air-dried and dissolved in 25 μL of water.

The CFB reaction mixture was prepared with 180 μL *E. coli* cell extract, 141 μL energy buffer, 12.6 μL DTT (100 mM stock conc.), 4.2 μL IPTG (50 mM stock conc.), 10 μL GamS (125 μM stock conc.), 10 μL MBP-tagged FusB (500 μM stock conc.), 10 μL MBP-tagged FusC (500 μM stock conc.) and 10 μL MBP-tagged FusE (500 μM stock conc.). The reaction mixture was then aliquoted into the 96-well plate with 3.6 μL in each well. Linear DNA template (1 μL, ~200 ng) was then added individually to initiate the CFB reactions. Reactions were quenched after incubating of 16 h at room temperature by addition of 5.4 μL ACN to each well (60% *v/v*). The plate was then subjected to centrifugation at 4,500 × $g$ for 15 min at 4 °C. The supernatant was analyzed by MALDI-TOF-MS with SA as the matrix.

For multiple FusA variants produced in a single reaction, each CFB reaction containing one FusA variant was initiated as described above. Afterward, 6 reactions were pooled, allowed to react at room temperature for 16 h, and then analyzed by MALDI-TOF-MS as above.

**Heatmap analysis for the amino acid occurrence at each position.** The observed occurrence ($O_{obs}$) of each residue at each position was calculated based on the sequences of FusC substrates or non-substrates. The predicted occurrence ($O_{pre}$) of each residue based on the degenerate NNK codon (N=A, T, C, G; K = T, G) was as follows: Glu: 3.1%; Gly: 6.3%; Asp: 3.1%; Met: 3.1%; Ala: 6.3%; Val: 6.3%; Asn: 3.1%; Lys: 3.1%; Thr: 6.3%; Ile: 3.1%; Gln: 3.1%; Arg: 9.4%; His: 3.1%; Phe: 3.1%; Ser: 9.4%; Leu: 9.4%; Tyr: 3.1%; Trp: 3.1%; Cys: 3.1%; Pro: 6.3%. The percent difference (D%) between the two was then calculated using:

$$\text{D\%} = (O_{obs} - O_{pre})/O_{pre} * 100\% \text{ if } O_{obs} \text{ is higher than } O_{pre}$$
$$\text{D\%} = (O_{obs} - O_{pre})/O_{obs} * 100\% \text{ if } O_{obs} \text{ is lower than } O_{pre}$$

**Calculation of percent surface area of lasso peptide ring region.** For each lasso peptide structure with a PDB file, the exposed surface area for each residue was calculated in UCSF Chimera.[7] The surface command for each strand calculated the solvent-excluded molecular surface. The solvent accessible surface area (SASA) was calculated by residue.[8] The SASA for the ring region was the sum of the residues in the macrocycle, while the SASA of the loop/tail was the sum of the remaining residues (**Table S4**).

**Compilation of all predicted lasso peptides found in NCBI.** A diverse set of known lasso leader peptidases (**Table S5**) were used as queries for a four-iteration Position-Specific Iterated BLAST (PSI-BLAST[9]) search (November 2020). Default settings were used except the maximum target sequence was increased to 20,000. The eleven searches were combined for a non-redundant list of 9,571 protein accession identifiers from Protein Family[10] PF13471 (transglut_core3) within the NCBI database. Each NCBI protein accession identifier was submitted to RODEO (Rapid ORF Description & Evaluation Online) using the lasso peptide scoring function.[11] Initial analysis yielded ~452,000 putative open reading frames co-occurring with the queried leader peptidase. Hypothetical peptides receiving a RODEO score <5 were disqualified from further analysis. Peptides receiving a score of 5 or greater (~38,500) are provided in Dataset 1. Peptides receiving a score of 10 or greater were considered lasso precursor peptides. Leader peptidase queries that did not co-occur with a putative lasso cyclase (PF00733, asparagine synthetase homologs) were also removed. Entries with the same leader peptidase query, where the ending nucleotide position was the same, but the starting positions were variable due to ambiguity over start codons, were considered identical. The longest of these entries with the highest RODEO lasso score was retained while the remaining were removed such that each unique core peptide sequence (same stop codon) occurred only once per leader peptidase query, resulting in ~8,000 precursor peptides. The ring size and acceptor residue were predicted and manually inspected. Entries with core sequences lacking a single Asp or Glu within position 7–16 were considered invalid and removed. In total, 7,701 total peptides were identified in the NBCI database that were encoded in a suspected lasso peptide BGC that appeared to be competent substrates.

**Bioinformatic analysis of naturally encoded lasso core peptides.** To reveal sequence trends in the core region of predicted lasso peptide precursors, any database entries (n = 7,701) with identical core sequences were removed, yielding 4,485 unique predicted core sequences. The acceptor site was determined as either the Asp or Glu that was in the 7$^{th}$, 8$^{th}$, or 9$^{th}$ core position. If multiple Asp or Glu were found at positions 7−9 for uncharacterized lasso peptide families, the acceptor site was presumed to yield the larger macrocycle, which was based on a recent statistical analysis of naturally encoded lasso precursor peptides (5.5%, 36.6%, and 57.9% for 7-mers, 8-mers, and 9-mers, respectively).[2] The only exception was if Glu was found at both positions 7 and 8, the acceptor site was presumed to be at position 7 (n = 170/7701, 2.2%), similar to the observed structure of the xanthomonins.[12] If there was neither Asp or Glu at position 7−9, the acceptor residue was assigned as the next-closest Asp or Glu to the N-terminal of the core peptide beyond position 9 (10-mers n = 505/7701, 6.6%, 11+-mers n = 318/7701, 4.1%).

The variability of the residues was calculated with non-redundant core sequences, either only sequences predicted to have a 9-residue macrolactam or 9-residue macrolactam with a Glu-acceptor (same macrocycle as fusilassin). Three different variability measurements were calculated: Shannon entropy,[13] consurf,[14] and relative entropy (**Table S6**).[15] The frequency of each amino acid at each position of the lasso peptides was then determined with the non-redundant core sequences. For analysis, only 9-mer predicted lasso peptides were used, the same macrocycle size as fusilassin. The frequency of each amino acid was compared to its expected frequency. The expected frequency was chosen to be the weighted average codon frequency for the most represented genus (**Table S7**)[16].

**Table S1. Genetically predicted lasso peptides used for FusA chimeric precursor generation.** The leader region of FusA was retained while the core sequences of 10 alternative precursor peptides were used (acceptor residue is red) to evaluate FusB/C/E compatibility. The FusA core sequence is aligned to each entry with bold and underlined representing identical and similar residues. Hyphens represent gaps in the alignment. Values in parenthesis indicate percent identity/similarity (lenient). Percent identity (%ID) of the lasso cyclase present in the alternative BGC is also compared with FusC (WP_104612995.1).

| Name | NCBI accession (lasso cyclase) | Bacterial strain | Core sequence (top)<br>FusA core sequence (bottom) | %ID to FusC |
|---|---|---|---|---|
| CelA | WP_068757176.1 | *Thermobifida cellulosilytica* TB100 | WIQGKWGLEIYLIFPRYL<br>WYTAEWGLELIFVFPRFI (44/100) | 77 |
| HalA | WP_068689552.1 | *Thermobifida halotolerans* | YKSG-RGLELWLFLPRMV<br>WYTAEWGLELIFVFPRFI (33/83) | 71 |
| MthA | WP_078762521.1 | *Marinactinospora thermotolerans* DSM 45154 | Y-NAINKLEIIFIWPRLFN<br>WYTAEWGLELIFVFPRFI (33/78) | 55 |
| NcaA | WP_052423756.1 | *Nonomuraea candida* | YV-GLRNRESLLGYPRNIW<br>WYTAEWGLELIFVFPRFI (28/56) | 49 |
| MobA | WP_051072671.1 | *Streptomyces mobaraensis* DSM 40847 | YI-GLEGSEPITHSFSKFW<br>WYTAEWGLELIFV-FPRFI (28/56) | 49 |
| RubA | WP_045701015.1 | *Streptomyces rubellomurinus* | AL-GLHGAEPFFPTLHTSWW<br>WYTAEWGLELIFVFPRFI (17/44) | 45 |
| NbsA | WP_089507158.1 | *Streptomyces* sp. NBS 14/10 | YF-GLTGYENVIHFYDRL<br>WYTAEWGLELIFVFP-RFI (28/61) | 44 |
| SruA | WP_107644840.1 | *Streptomyces* sp. Ru87 | RG-GEPIWEEVVVPWDYWV<br>WYTAEWGLELIFVFPRFI (17/56) | 44 |
| SleA | WP_048574008.1 | *Streptomyces leeuwenhoekii* | LYGVRN-DEEINWHFDYWT<br>WYTAEWGLELIFVFPRFI (11/50) | 42 |
| AreA | WP_048574008.1 | *Actinoplanes regularis* | TGMYGRRGYERTLQT-KA (D10E[1])<br>WYT-AEWGLELIFVFPRFI (22/44) | 41 |
| CitA[2] | WP_038518372.1 | *Streptomyces albulus* | LLGLA-GNERTLVLSKN (D8E[1])<br>WYTAEWGLELIFVFPRFI (17/39) | 38 |

[1]The acceptor residues of AreA and CitA were changed from Asp to Glu.

[2]CitA is the precursor peptide of citrulassin A, which was reported previously[2] and not included in this work.

**Table S2. Nucleotide sequences of chimeric precursor peptides.** The codons are optimized for *E. coli*. All sequences are provided 5′ to 3′. Restriction sites (BamHI, HindIII) are underlined. LP, leader peptide; CP, core peptide.

**$\textit{FusA}_{LP}$-$\textit{CelA}_{CP}$**

GGATCCATGGAAAAGAAGAAGTACACGGCCCCTCAGCTGGCTAAGGTAGGCGAATTTAAGGAAGCCACTGGATGGATTCAAGG
CAAATGGGGCCTGGAGATTTACCTGATCTTCCCGCGCTATTTGTAGAAGCTT

**$\textit{FusA}_{LP}$-$\textit{HalA}_{CP}$**

GGATCCATGGAAAAGAAGAAGTACACGGCCCCACAACTGGCCAAGGTTGGTGAGTTTAAGGAAGCGACTGGTTACAAGTCGGG
TCGTGGTTTAGAGCTTTGGCTGTTTCTCCCCCGTATGGTATAAAAGCTT

**$\textit{FusA}_{LP}$-$\textit{MthA}_{CP}$**

GGATCCATGGAAAAGAAGAAGTACACGGCCCCTCAACTGGCCAAGGTTGGCGAGTTTAAGGAAGCAACAGGTTACAATGCCAT
CAATAAGTTAGAGATCATCTTCATCTGGCCTCGTTTATTCAATTAAAAGCTT

**$\textit{FusA}_{LP}$-$\textit{NcaA}_{CP}$**

GGATCCATGGAAAAGAAGAAGTACACGGCCCCTCAATTGGCAAAGGTGGGTGAGTTTAAGGAAGCGACAGGTTACGTCGGATT
GCGTAATCGTGAGTCGCTTTTAGGGTACCCCCGTAATATCTGGTAAAAGCTT

**$\textit{FusA}_{LP}$-$\textit{MobA}_{CP}$**

GGATCCATGGAAAAGAAGAAGTACACGGCACCCCAACTCGCTAAGGTGGGCGAGTTTAAGGAAGCGACGGGATACATCGGTCT
CGAGGGAAGCGAGCCTATCACTCACTCGTTCTCCAAGTTCTGGTAAAAGCTT

**$\textit{FusA}_{LP}$-$\textit{RubA}_{CP}$**

GGATCCATGGAAAAGAAGAAGTACACGGCACCACAACTCGCTAAGGTAGGTGAGTTTAAGGAAGCGACGGGTGCCTTAGGACT
CCACGGTGCTGAGCCTTTCTTCCCTACGCTCCACACGTCGTGGTGGTAAAAGCTT

**$\textit{FusA}_{LP}$-$\textit{NbsA}_{CP}$**

GGATCCATGGAAAAGAAGAAGTACACTGCACCTCAATTGGCCAAGGTGGGCGAGTTTAAGGAAGCGACTGGGTACTTCGGTTT
GACTGGGTACGAGAATGTAATCCACTTCTACGACCGTCTCTAAAAGCTT

**$\textit{FusA}_{LP}$-$\textit{SruA}_{CP}$**

GGATCCATGGAAAAGAAGAAGTACACTGCCCCACAATTGGCAAAGGTGGGTGAGTTTAAGGAAGCAACGGGTCGTGGTGGTGA
GCCCATCTGGGAAGAGGTTGTCGTCCCATGGGACTACTGGGTTTAAAAGCTT

**$\textit{FusA}_{LP}$-$\textit{SleA}_{CP}$**

GGATCCATGGAAAAGAAGAAGTACACAGCACCTCAACTCGCCAAGGTAGGTGAGTTTAAGGAAGCGACTGGTTTATACGGTGT
CCGTAATGACGAGGAGATCAATTGGCACTTCGACTACTGGACTTAAAAGCTT

**$\textit{FusA}_{LP}$-$\textit{AreA}_{CP}$(D10E)**

GGATCCATGGAAAAGAAGAAGTACACGGCACCTCAACTCGCAAAGGTTGGCGAGTTTAAGGAAGCGACGGGTATGTACGGACG
TCGTGGTTACGAACGTACGCTCCAAACGAAGGCCTGAAAGCTT

**Table S3. Nucleotide sequences of FusA degenerate libraries.** All sequences are provided 5′ to 3′ direction. *Top*, "Ring" library with the NNK (orange) region located in the ring region of FusA. *Bottom*, "Loop" library with NNK codons appearing in the loop region of FusA. Restriction sites (KpnI and SacI) are underlined. Color-coding: blue, T7 promotor/terminator; purple, lac operator; red, ribosome-binding sequence. The FusA coding sequence is in bold.

**FusA ring degenerate library:**

GGTACCATCCCGCGAAATTAATACGACTCACTATAGGGGAATTGTGAGCGGATAACAATTCCCCTCTA
GAAATAATTTTGTTTAACTTTAAGAAGGAGATATACC**ATGGAAAAGAAGAAGTACACCGCTCCGCAGC**
**TCGCTAAGGTCGGCGAATTCAAGGAGGCCACCGGCTGGNNKNNKNNKNNKNNKGGCCTCGAGCTGATC**
**TTCGTCTTCCCGCGCTTCATCTGA**AGGAAGCTGAGTTGGCTGCTGCCACCGCTGAGCAATAACTAGCA
TAACCCCTTGGGGCCTCTAAACGGGTCTTGAGGGGTTTTTTGGAGCTC

**FusA loop degenerate library:**

GGTACCATCCCGCGAAATTAATACGACTCACTATAGGGGAATTGTGAGCGGATAACAATTCCCCTCTA
GAAATAATTTTGTTTAACTTTAAGAAGGAGATATACC**ATGGAAAAGAAGAAGTACACCGCTCCGCAGC**
**TCGCTAAGGTCGGCGAATTCAAGGAGGCCACCGGCTGGTACACCGCGGAATGGGGCCTCGAGNNKNNK**
**NNKNNKNNKCCGCGCTTCATCTGA**AGGAAGCTGAGTTGGCTGCTGCCACCGCTGAGCAATAACTAGCA
TAACCCCTTGGGGCCTCTAAACGGGTCTTGAGGGGTTTTTTGGAGCTC

**Table S4. Expected surface area for ring region among available lasso peptide structures in PDB.** The surface areas for 21 lasso peptides were analyzed and the estimated surface areas for both ring and loop + tail were calculated. Blue, ring region; red, acceptor residue. Underlined residues are not shown in the crystal structure.

| Name | Core sequences | PDB entry | Estimated surface area (ring) (Å$^2$) | Estimated surface area (loop+tail) (Å$^2$) | % estimated surface area (ring) |
|---|---|---|---|---|---|
| microcin J25 | GGAGHVPEYFVGIGTPISFYG | 1Q71 | 561 | 1170 | 32% |
| astexin-1 | GLSQGVEPDIGQTYFEESRINQD | 2LTI | 632 | 1231 | 34% |
| caulosegnin I | GAFVGQPEAVNPLGREIQG | 2LX6 | 647 | 1132 | 36% |
| astexin-3 | GPTPMVGLDSVSGQYWDQHAPLAD | 2M8F | 576 | 1461 | 28% |
| lassomycin | GLRRLFADQLVGRRNI | 2MAI | 771 | 1063 | 42% |
| xanthomonin II | GGPLAGEEMGGITT | 2MFV | 519 | 752 | 41% |
| caulonodin V | SIGDSGLRESMSSQTYWP | 2MLJ | 800 | 922 | 46% |
| streptomonomicin | SLGSSPYNDILGYPALIVIYP | 2MW3 | 736 | 1320 | 36% |
| citrocin | GGVGKIIEYFIGGGVGRYG | 6MW6 | 633 | 980 | 39% |
| chaxapeptin | GFGSKPLDSFGLNFF | 2N5C | 703 | 818 | 46% |
| astexin-2 | GLTQIQALDSVSGQFRDQLG | 2N6U | 828 | 1153 | 42% |
| xanthomonin I | GGPLAGEEIGGFNVPGISEE | 4NAG | 335 | 774 | 30% |
| caulosegnin II | GTLTPGLPEDFLPGHYMPG | 5D9E | 714 | 1390 | 34% |
| sphaericin | GLPIGWWIERPSGWYFPI | 5GVO | 857 | 1027 | 45% |
| sphingopyxin I | GIEPLGPVDEDQGEHYLFAGG | 5JQF | 543 | 839 | 39% |
| rubrivinodin | GAPSLINSEDNPAFPQRV | 5OQZ | 698 | 909 | 43% |
| benenodin-1 | GVGFGRPDSILTQEQAKPMGLDRD | 5TJ1 | 608 | 1051 | 37% |
| acinetodin | GGKGPIFETWVTEGNYYG | 5UI6 | 620 | 962 | 39% |
| klebsidin | GSDGPIIEFFNPNGVMHYG | 5UI7 | 592 | 1120 | 35% |
| subterisin | GPPGDRIEFGVLAQLPGLDRD | 5XM4 | 722 | 908 | 44% |
| ubonodin | GGDGSIAEYFNRPMHIHDWQIMDSGYYG | 6POR | 516 | 2184 | 19% |

**average=38%**

**Table S5. Protein accession identifiers used to catalog all lasso peptide BGCs in NCBI.** Each protein is lasso leader peptidase (PF13471) from a known lasso peptide biosynthetic gene cluster (BGC) .

| **Lasso peptide** | **Genus species** | **Phylum** | **Class** | **Linkage** | **NCBI accession** |
|---|---|---|---|---|---|
| citrulassin A | *Streptomyces albus* | Actinobacteria | II | Leu-Asp 8mer | WP_030547855.1 |
| specialicin | *Streptomyces specialis* | Actinobacteria | I | Cys-Asp 9mer | WP_059005882.1 |
| xanthomonin III | *Xanthomonas citri* | Proteobacteria | II | Gly-Glu 7mer | WP_050545040.1 |
| pseudomycoidin | *Bacillus pseudomycoides* | Firmicutes | II | Ala-Asp 9mer | EEM14410.1 |
| caulonodin VI | *Caulobacter* sp. | Proteobacteria | II | Ala-Glu 9mer | WP_049771798.1 |
| LP2006 | *Nocardiopsis alba* | Actinobacteria | IV | Gly-Glu 8mer | WP_014912067.1 |
| pandonodin | *Pandoraea norimbergensis* | Proteobacteria | IV | Gly-Glu 8mer | WP_058377878.1 |
| 9401-LP1 | *Streptomyces* sp. | Actinobacteria | III | Ala-Asp 9mer | ARA91542.1 |
| paeninodin | *Paenibacillus dendritiformis* | Firmicutes | II | Ala-Asp 9mer | EHQ60563.1 |
| capistruin | *Burkholderia thailandensis* | Proteobacteria | II | Gly-Asp 9mer | ABC38564.1 |
| lariatin B | *Rhodococcus jostii* | Actinobacteria | II | Gly-Glu 8mer | BAL72549.1 |

**Table S6. Natural variability in the set of genomically predicted lasso peptides.** The set has been reduced to only include one entry for each unique core sequence ($n$ = 4,485). The position with the greatest variability per metric is highlighted in red while the next three most variable positions are in bold. Acceptor residues were predicted based on the position of the Asp and/or Glu residues as described in the Methods. White background indicates ring region while gray background indicates loop/tail region.

| Core position | 9-residue ring, Asp- and Glu-acceptor | | | 9-residue ring, Glu only-acceptor | | |
|---|---|---|---|---|---|---|
| | Shannon entropy | Consurf (1−9) | Relative entropy | Shannon entropy | Consurf (1−9) | Relative entropy |
| 1 | 2.91 | 6.5 | 0.155 | 3.05 | 7 | 0.156 |
| 2 | 3.74 | 5.5 | 0.074 | 4.02 | 5.5 | 0.054 |
| 3 | 3.91 | **4.5** | 0.044 | 4.03 | 5.5 | **0.020** |
| 4 | 3.87 | **4.5** | 0.044 | 4.00 | **4** | **0.026** |
| 5 | 3.85 | **4.5** | 0.046 | 3.84 | 5 | 0.036 |
| 6 | 3.94 | **4.5** | 0.037 | 3.96 | **5.5** | 0.037 |
| 7 | **4.21** | **3.5** | **0.025** | **4.14** | **3** | 0.036 |
| 8 | 3.99 | 5.5 | 0.204 | 3.97 | 5.5 | 0.269 |
| 9 | 0.89 | 9 | 0.386 | 0 | 9 | 0.489 |
| 10 | 4.10 | 5.5 | 0.201 | **4.12** | 5.5 | 0.248 |
| 11 | 4.01 | **4.5** | **0.027** | 4.06 | 5 | **0.023** |
| 12 | 3.97 | 5.5 | 0.038 | 4.04 | 5.5 | 0.029 |
| 13 | 4.07 | **4.5** | 0.035 | **4.17** | 5.5 | **0.028** |
| 14 | **4.12** | 5.5 | **0.030** | 4.10 | 5.5 | 0.029 |
| 15 | **4.15** | 5.5 | **0.031** | **4.11** | 5.5 | 0.040 |
| 16 | **4.20** | **4.5** | 0.035 | **4.11** | **4.5** | 0.057 |

**Table S7. Determination of the "expected frequency" as a weighted average of the most frequent genus to have a predicted lasso peptide gene cluster.** The genus, number of CDSs included in the analysis, the residue frequency and the weight for the average, is shown. The determined frequency is Glu: 6.0%; Gly: 8.2%; Asp: 5.7%; Met: 2.2%; Ala: 11.7%; Val: 7.6%; Asn: 2.9%; Lys: 3.9%; Thr: 5.6%; Ile: 5.1%; Gln: 3.3%; Arg: 6.8%; His: 2.1%; Phe: 3.6%; Ser: 5.5%; Leu: 9.8%; Tyr: 2.6%; Trp: 1.4%; Cys: 0.8%; Pro: 5.2%.

| Genus (Number of CDS) | *Streptomyces* (11721966) | *Bacillus* (24372190) | *Paenibacillus* (3179769) | *Mesorhizobiu m* (3258891) | *Sphingobium* (473562) | *Xanthomonas* (6020839) | *Shingomonas* (1262391) | *Sphingopyxis* (236936) | *Novosphingob ium* (388272) | *Caulobacter* (230357) | *Streptococcus* (3145281) | *Burkholderia* (22235578) | *Brevundimo nas* (191591) | *Stenotrophom onas* (1970883) | *Amycolatopsi s* (790150) | *Clostridium* (3257970) | *Actinomyces* (363766) | *Acidobacteria* (230631) | *Erythrobacter* (149133) |
|---|---|---|---|---|---|---|---|---|---|---|---|---|---|---|---|---|---|---|---|
| E | 5.7% | 7.4% | 6.7% | 5.5% | 5.2% | 4.8% | 5.7% | 5.3% | 5.3% | 5.3% | 7.1% | 4.7% | 5.6% | 5.1% | 5.7% | 7.6% | 5.8% | 5.5% | 6.5% |
| G | 9.6% | 6.8% | 7.5% | 8.6% | 8.8% | 8.2% | 3.6% | 9.0% | 8.9% | 9.0% | 6.6% | 8.3% | 9.0% | 8.6% | 9.3% | 4.7% | 9.0% | 8.1% | 8.8% |
| D | 5.9% | 4.8% | 5.1% | 5.7% | 6.1% | 5.7% | 6.7% | 6.2% | 5.8% | 5.9% | 5.6% | 5.7% | 6.0% | 5.7% | 5.9% | 5.6% | 5.9% | 5.1% | 6.2% |
| M | 1.6% | 2.7% | 2.8% | 2.4% | 2.5% | 2.1% | 2.5% | 2.4% | 2.4% | 2.1% | 2.5% | 2.1% | 2.3% | 2.1% | 1.5% | 2.7% | 2.0% | 2.2% | 2.4% |
| A | 13.7% | 6.9% | 8.2% | 12.4% | 13.1% | 13% | 15.4% | 13.6% | 13.3% | 13.9% | 7.5% | 14.2% | 13.8% | 12.9% | 13.3% | 5.9% | 13.2% | 11% | 12.6% |
| V | 8.4% | 7.1% | 7.0% | 7.4% | 6.9% | 7.3% | 8.2% | 7.0% | 7.2% | 7.6% | 7.0% | 7.5% | 7.6% | 7.4% | 8.9% | 6.5% | 8.7% | 7.3% | 7.0% |
| N | 1.7% | 4.5% | 3.8% | 2.7% | 2.6% | 2.6% | 2.8% | 2.6% | 2.6% | 2.4% | 4.5% | 2.6% | 2.4% | 2.6% | 1.9% | 6.3% | 2.1% | 3.3% | 2.6% |
| K | 2.1% | 7.3% | 5.3% | 3.7% | 3.0% | 2.7% | 3.1% | 3.1% | 2.9% | 3.3% | 7.0% | 2.7% | 2.8% | 2.7% | 2.3% | 9.2% | 2.3% | 3.6% | 3.0% |
| T | 6.2% | 5.5% | 5.5% | 5.3% | 5.1% | 5.2% | 6.2% | 5.2% | 5.3% | 5.4% | 5.7% | 5.2% | 5.3% | 4.9% | 6.0% | 5.0% | 6.3% | 6.0% | 5.2% |
| I | 3.0% | 7.9% | 6.6% | 5.4% | 5.2% | 4.1% | 5.5% | 5.1% | 4.8% | 4.3% | 7.4% | 4.3% | 4.4% | 4.0% | 3.5% | 9.8% | 4.2% | 4.9% | 5.0% |
| Q | 2.7% | 3.8% | 3.9% | 3.0% | 3.3% | 4.5% | 3.4% | 3.0% | 3.2% | 3.1% | 4.2% | 3.2% | 3.2% | 4.4% | 2.7% | 2.5% | 3.2% | 3.8% | 3.2% |
| R | 8.3% | 3.9% | 5.1% | 6.9% | 7.5% | 7.6% | 8.6% | 7.3% | 7.2% | 7.2% | 4.2% | 8.0% | 7.6% | 7.5% | 7.8% | 3.6% | 7.5% | 6.6% | 7.1% |
| H | 2.3% | 2.2% | 2.1% | 2.0% | 2.1% | 2.3% | 2.2% | 2.0% | 2.1% | 1.8% | 2.0% | 2.4% | 1.8% | 2.2% | 2.2% | 1.4% | 0.4% | 2.2% | 1.9% |
| F | 2.7% | 4.6% | 4.1% | 3.9% | 3.5% | 3.2% | 3.9% | 3.6% | 3.5% | 3.5% | 4.7% | 3.6% | 3.4% | 3.3% | 3.0% | 4.4% | 2.7% | 3.9% | 3.7% |
| S | 5.1% | 6.0% | 6.4% | 5.7% | 5.4% | 5.6% | 5.7% | 5.2% | 5.4% | 5.2% | 5.0% | 5.4% | 5.1% | 5.4% | 5.2% | 6.4% | 6.4% | 6.3% | 5.4% |
| L | 10.3% | 9.6% | 10.1% | 9.9% | 9.9% | 10.7% | 5.3% | 9.7% | 9.9% | 10% | 10.4% | 9.9% | 9.9% | 11.1% | 10.5% | 9.3% | 10.1% | 10% | 9.7% |
| Y | 2.0% | 3.7% | 3.5% | 2.2% | 2.3% | 2.4% | 2.5% | 2.2% | 2.2% | 2.2% | 3.9% | 2.4% | 2.0% | 2.3% | 2.0% | 4.2% | 2.1% | 2.7% | 2.2% |
| W | 1.5% | 1.0% | 1.3% | 1.4% | 1.5% | 1.6% | 1.6% | 1.5% | 1.5% | 1.5% | 0.9% | 1.4% | 1.5% | 1.6% | 1.5% | 0.8% | 1.5% | 1.4% | 1.4% |
| C | 0.8% | 0.8% | 0.8% | 0.9% | 0.8% | 1.0% | 0.8% | 0.8% | 0.9% | 0.7% | 0.6% | 1.0% | 0.7% | 0.8% | 0.8% | 1.3% | 0.8% | 0.9% | 0.8% |
| P | 6.3% | 3.5% | 4.1% | 5.1% | 5.3% | 5.3% | 6.1% | 5.3% | 5.4% | 5.6% | 3.3% | 5.4% | 5.5% | 5.3% | 6.0% | 2.9% | 5.8% | 5.4% | 5.1% |
| Weight of average | 16.7% | 10% | 5.5% | 3.6% | 3.5% | 3.0% | 2.8% | 2.4% | 2.4% | 2.0% | 1.5% | 1.3% | 1.2% | 1.1% | 1.1% | 1.0% | 0.9% | 0.6% | 0.6% |

**Table S8. Oligonucleotide primers used in this study.** All sequences are provided in the 5′ to 3′ direction. "f" and "r" indicate a forward or reverse primer, respectively. LP, leader peptide; CP, core peptide; capital letters, mutagenized codon; capitalized and italics, restriction enzyme recognition site.

| Number | Primer name | Oligonucleotide sequence |
|---|---|---|
| 1 | Cellulassin BGC_f | aaa*GGATCC*atgcaggaaaagaagaccc |
| 2 | Cellulassin BGC_r | aaa*GCGGCCGC*tcagtcgacggtgatcagg |
| 3 | FusA$_{LP}$-chimeric_f | cgc*GGATCC*atggaaaagaagaagtacac |
| 4 | FusA$_{LP}$-CelA$_{CP}$_r | ccc*AAGCTT*ctacaaatagcgcgggaag |
| 5 | FusA$_{LP}$-HalA$_{CP}$_r | ccc*AAGCTT*ttataccatacgggggag |
| 6 | FusA$_{LP}$-MthA$_{CP}$_r | ccc*AAGCTT*ttaattgaataaacgaggccag |
| 7 | FusA$_{LP}$-NcaA$_{CP}$_r | ccc*AAGCTT*ttaccagatattacggggg |
| 8 | FusA$_{LP}$-MobA$_{CP}$_r | ccc*AAGCTT*ttaccagaacttggagaacg |
| 9 | FusA$_{LP}$-NbsA$_{CP}$_r | ccc*AAGCTT*ttacccaccgaaagtaccg |
| 10 | FusA$_{LP}$-SleA$_{CP}$_r | ccc*AAGCTT*ttaagtccagtagtcgaagtg |
| 11 | FusA$_{LP}$-RubA$_{CP}$_r | ccc*AAGCTT*ttaccaccacgacgtgtg |
| 12 | FusA$_{LP}$-SruA$_{CP}$_r | ccc*AAGCTT*ttaaacccagtagtcccatg |
| 13 | FusA$_{LP}$-AreA$_{CP}$_r | ccc*AAGCTT*tcaggccttcgtttgg |
| 14 | FusA_Y2A_f | ccaccggctggGCGaccgcggaatggggcctcgagc |
| 15 | FusA_Y2A_r | cattccgcggtCGCccagccggtggcctccttgaat |
| 16 | FusA_T3A_f | ccggctggtacGCCgcggaatggggcctcgagctga |
| 17 | FusA_T3A_r | ccccattccgcCCGgtaccagccggtggcctccttg |
| 18 | FusA_E5A_f | ggtacaccgcgGCCtggggcctcgagctgatcttcg |
| 19 | FusA_E5A_r | tcgaggccccaGGCcgcggtgtaccagccggtggcc |
| 20 | FusA_W6A_f | acaccgcggaaGCGggcctcgagctgatcttcgtct |
| 21 | FusA_W6A_r | agctcgaggccCGCttccgcggtgtaccagccggtg |
| 22 | FusA_G7A_f | ccgcggaatggGCGctcgagctgatcttcgtcttcc |
| 23 | FusA_G7A_r | atcagctcgagCGCccattccgcggtgtaccagccg |
| 24 | FusA_L8A_f | cggaatggggcGCGgagctgatcttcgtcttcccgc |
| 25 | FusA_L8A_r | aagatcagctcCGCgccccattccgcggtgtaccag |
| 26 | FusA_L10A_f | ggggcctcgagGCGatcttcgtcttcccgcgcttca |
| 27 | FusA_L10A_r | aagacgaagatCGCctcgaggccccattccgcggtg |
| 28 | FusA_I11A_f | gcctcgagctgGCGttcgtcttcccgcgcttcatct |
| 29 | FusA_I11A_r | gggaagacgaaCGCcagctcgaggccccattccgcg |
| 30 | FusA_F12A_f | tcgagctgatcGCGgtcttcccgcgcttcatctgag |
| 31 | FusA_F12A_r | cgcgggaagacCGCgatcagctcgaggccccattcc |
| 32 | FusA_V13A_f | agctgatcttcGCGttcccgcgcttcatctgagcgg |
| 33 | FusA_V13A_r | aagcgcgggaaCGCgaagatcagctcgaggccccat |

| | | |
|---|---|---|
| 34 | FusA_F14A_f | tgatcttcgtcGCGccgcgcttcatctgagcggccg |
| 35 | FusA_F14A_r | atgaagcgcggCGCgacgaagatcagctcgaggccc |
| 36 | FusA_P15A_f | tcttcgtcttcGCGcgcttcatctgagcggccgcac |
| 37 | FusA_P15A_r | cagatgaagcgCGCgaagacgaagatcagctcgag |
| 38 | FusA_R16A_f | tcgtcttcccgGCGttcatctgagcggccgcactc |
| 39 | FusA_R16A_r | gctcagatgaaCGCcgggaagacgaagatcagctc |
| 40 | FusA_F17A_f | tcttcccgcgcGCGatctgagcggccgcactcgag |
| 41 | FusA_F17A_r | gccgctcagatCGCgcgcgggaagacgaagatcag |
| 42 | FusA_I18A_f | cttcccgcgcttcGCGtgagcggccgcactcgagc |
| 43 | FusA_I18A_r | cggccgctcaCGCgaagcgcgggaagacgaagatc |
| 44 | pET28_CFB_linear DNA_f | gctatcatgccataccgcgaaaggttttgcgccattcg |
| 45 | pET28_CFB_linear DNA_r | aaccgtctatcagggcgatggcccactacgtgaaccatc |
| 46 | FusA DEGlib_KpnI_f | cgc*GGTACC*atcccgcgaaattaatacgactcactatagg |
| 47 | FusA DEGlib_SacI_r | cgc*GAGCTC*caaaaaacccctcaagacccgtttagagg |
| 48 | pET28_DEGlib_SacI_f | cgc*GAGCTC*ctgaaaggaggaactatatcc |
| 49 | pET28_DEGlib_KpnI_r | cgg*GGTACC*cgagatctcgatcctctac |
| 50 | MthA_ring to FusA_f | TACACCGCGGAATGGGGCttagagatcatcttcatctgg |
| 51 | NcaA_ring to FusA_f | TACACCGCGGAATGGGGCcgtgagtcgcttttagggtac |
| 52 | MobA_ring to FusA_f | TACACCGCGGAATGGGGCagcgagcctatcactcactcg |
| 53 | NbsA_ring to FusA_f | TACACCGCGGAATGGGGCtacgagaatgtaatccacttc |
| 54 | MthA_CanA_MobA_NbsA_ring to FusA_r | GCCCCATTCCGCGGTGTAgtaacctgttgcttccttaaac |
| 55 | MthA_loop to FusA_f | CTGATCTTCGTCTTCCCGcgtttattcaattaaaagcttgcg |
| 56 | NcaA_loop to FusA_f | CTGATCTTCGTCTTCCCGcgtaatatctggtaaaagcttgcg |
| 57 | MobA_loop to FusA_f | CTGATCTTCGTCTTCCCGaagttctggtaaaagcttgcg |
| 58 | NbsA_loop to FusA_f | CTGATCTTCGTCTTCCCGcgtctctaaaagcttgcg |
| 59 | MthA_loop to FusA_r | CGGGAAGACGAAGATCAGctctaacttattgatggcattg |
| 60 | NcaA_loop to FusA_r | CGGGAAGACGAAGATCAGctcacgattacgcaatccgac |
| 61 | MobA_loop to FusA_r | CGGGAAGACGAAGATCAGctcgcttccctcgagacc |
| 62 | NbsA_loop to FusA_r | CGGGAAGACGAAGATCAGctcgtacccagtcaaacc |
| 63 | FusA ring_MNMYM_f | ATGAACATGTATATGggcctcgagctgatcttcgtcttcccg |
| 64 | FusA ring_MNMYM_r | CATATACATGTTCATccagccggtggcctccttgaattcg |
| 65 | FusA ring_FYNWK_f | TTTTATAACTGGAAAggcctcgagctgatcttcgtcttcccg |
| 66 | FusA ring_FYNWK_r | TTTCCAGTTATAAAAccagccggtggcctccttgaattcg |
| 67 | FusA ring_NMQIY_f | AACATGCAGATTTATggcctcgagctgatcttcgtcttcccg |
| 68 | FusA ring_NMQIY_r | ATAAATCTGCATGTTccagccggtggcctccttgaattcg |
| 69 | FusA ring_MYTFQ_f | ATGTATACCTTTCAGggcctcgagctgatcttcgtcttcccg |
| 70 | FusA ring_MYTFQ_r | CTGAAAGGTATACATccagccggtggcctccttgaattcg |

| | | |
|---|---|---|
| 71 | FusA ring_TWTEM_f | ACCTGGACCGAAATGggcctcgagctgatcttcgtcttcccg |
| 72 | FusA ring_TWTEM_r | CATTTCGGTCCAGGTccagccggtggcctccttgaattcg |
| 73 | FusA ring_IKEVT_f | ATTAAAGAAGTGACCggcctcgagctgatcttcgtcttcccg |
| 74 | FusA ring_IKEVT_r | GGTCACTTCTTTAATccagccggtggcctccttgaattcg |
| 75 | FusA ring_QDWFM_f | CAGGATTGGTTTATGggcctcgagctgatcttcgtcttcccg |
| 76 | FusA ring_QDWFM_r | CATAAACCAATCCTGccagccggtggcctccttgaattcg |
| 77 | FusA ring_FLRCL_f | TTTCTGCGCTGCCTGggcctcgagctgatcttcgtcttcccg |
| 78 | FusA ring_FLRCL_r | CAGGCAGCGCAGAAAccagccggtggcctccttgaattcg |
| 79 | FusA ring_IDRSY_f | ATTGATCGCAGCTATggcctcgagctgatcttcgtcttcccg |
| 80 | FusA ring_IDRSY_r | ATAGCTGCGATCAATccagccggtggcctccttgaattcg |
| 81 | FusA ring_LKNFT_f | CTGAAAAACTTTACCggcctcgagctgatcttcgtcttcccg |
| 82 | FusA ring_LKNFT_r | GGTAAAGTTTTTCAGccagccggtggcctccttgaattcg |

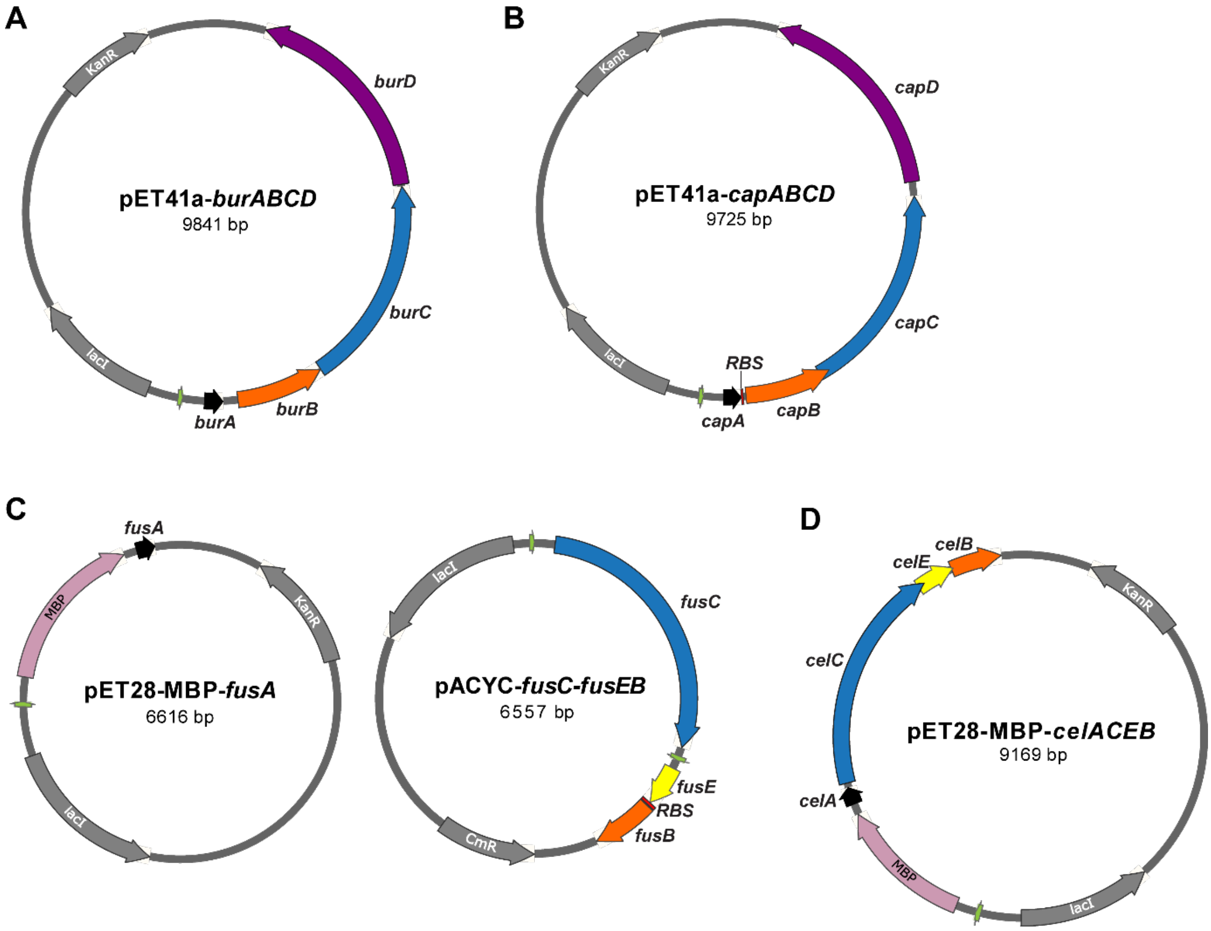

**Figure S1. Plasmid maps for constructs used in CFB reactions. (A)** Plasmid map of pET41a-*burABCD*. The BGC of burhizin from *Burkholderia rhizoxinica* was cloned into pET41a using the native gene organization. **(B)** Plasmid map of pET41a-*capABCD*. The BGC of capistruin from *Burkholderia thailandensis* was cloned into pET41a using the native organization with the intergenic region between *capA* and *capB* replaced with an *E. coli* ribosome-binding site (RBS). **(C)** Plasmid maps of pET28-MBP-*fusA* and pACYC-*fusC-fusEB*. The *fusA* (fusilassin precursor peptide) gene (or chimeric precursor) was cloned into pET28-MBP. The lasso cyclase gene (*fusC*) was cloned into the first multiple cloning site of pACYC. The RiPP recognition element (*fusE*) and leader peptidase (*fusB*) were Gibson assembled into the second multiple cloning site of pACYC with an *E. coli* RBS inserted between *fusE* and *fusB*.[2] **(D)** The plasmid map of pET28-MBP-*celACEB*. The BGC of cellulassin from *Thermobifida cellulosilytica* was cloned into pET28-MBP using the native organization. Green, T7 promotor; Red, inserted RBS. The NCBI accession identifiers for the cellulassin biosynthetic proteins are: CelA, WP_157080207.1; CelC, WP_068757176.1; CelE, WP_068757174.1; CelB, WP_068757172.1.

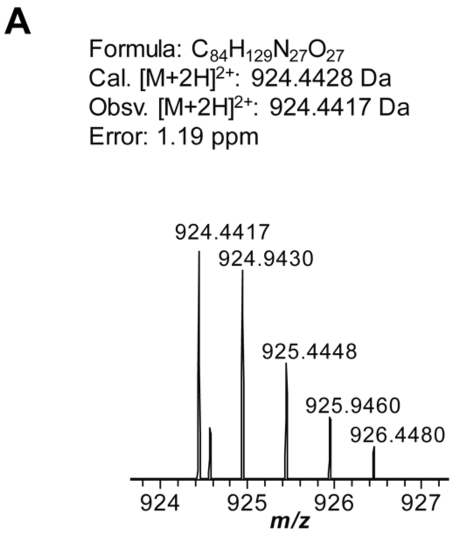

B

| Ion | Calculated mass (Da) | Observed mass (Da) | Error (ppm) |
|---|---|---|---|
| b8 | 773.3582 | 773.3552 | 3.88 |
| b9 | 872.4266 | 872.4242 | 2.75 |
| b10 | 1001.4692 | 1001.4668 | 2.40 |
| b11 | 1072.5063 | 1072.5035 | 2.61 |
| b12 | 1129.5278 | 1129.5265 | 1.15 |
| $[b13+2H]^{2+}$ | 643.3178 | 643.3170 | 1.24 |
| $[b14+2H]^{2+}$ | 736.3575 | 736.3556 | 2.58 |
| $[b15+2H]^{2+}$ | 779.8734 | 779.8697 | 4.74 |
| $[b16+2H]^{2+}$ | 837.3868 | 837.3849 | 2.27 |
| y3 | 377.1785 | 377.1767 | 4.77 |
| y4 | 563.2578 | 563.2563 | 2.66 |
| $[y5+2H]^{2+}$ | 360.1828 | 360.1816 | 3.33 |
| $[y6+2H]^{2+}$ | 388.6925 | 388.6924 | 0.26 |
| $[y7+2H]^{2+}$ | 424.2121 | 424.2110 | 2.59 |
| $[y8+2H]^{2+}$ | 488.7334 | 488.7325 | 1.84 |
| $[y9+2H]^{2+}$ | 538.2676 | 538.2668 | 1.49 |

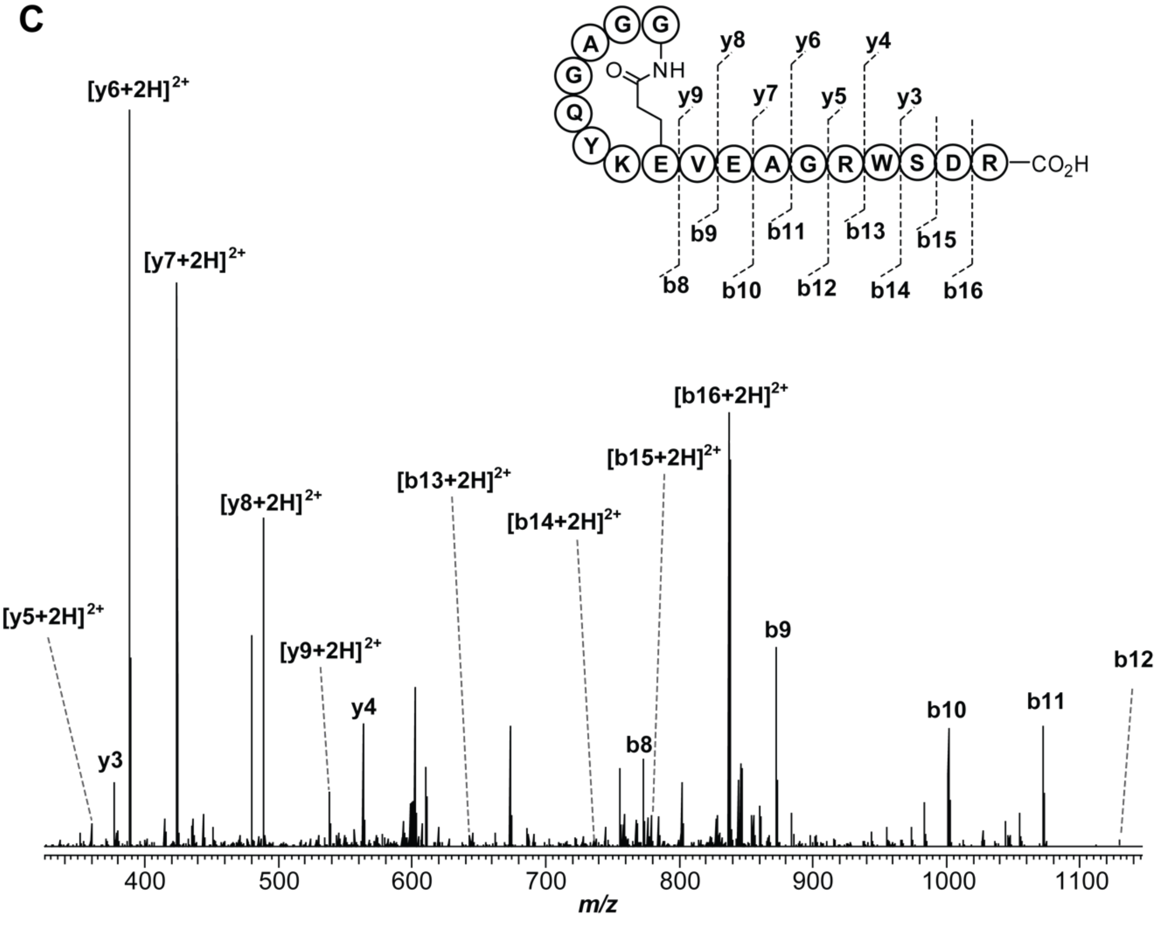

**Figure S2. High-resolution and tandem MS of burhizin produced from CFB. (A)** High-resolution mass spectrum of burhizin, produced from CFB with pET41a-*burABCD* as the DNA template. **(B)** Mass assignment for the $b^+$ and $y^+$ ions generated from the CID of burhizin. **(C)** Tandem MS spectrum of burhizin, consistent with a Gly1-Glu8 macrolactam.

A

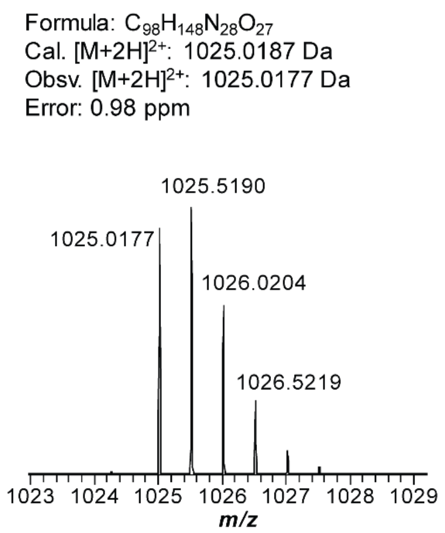

B

| Ion | Calculated mass (Da) | Observed mass (Da) | Error (ppm) |
|---|---|---|---|
| b11 | 1110.5332 | 1110.5311 | 1.89 |
| b12 | 1209.6016 | 1209.5990 | 2.15 |
| b13 | 1322.6857 | 1322.6831 | 1.97 |
| b14 | 1409.7177 | 1409.7146 | 2.20 |
| y4-$NH_3$ | 467.1931 | 467.1922 | 1.93 |
| y4 | 484.2196 | 484.2187 | 1.86 |
| y5-$NH_3$ | 623.2942 | 623.2927 | 2.41 |
| y5 | 640.3207 | 640.3195 | 1.87 |
| y6-$NH_3$ | 710.3262 | 710.3249 | 1.83 |
| y6 | 727.3527 | 727.3513 | 1.92 |
| y7-$NH_3$ | 823.4103 | 823.4088 | 1.82 |
| y7 | 840.4368 | 840.4353 | 1.78 |
| y8-$NH_3$ | 922.4787 | 922.4769 | 1.95 |
| y8 | 939.5052 | 939.5036 | 1.70 |

C

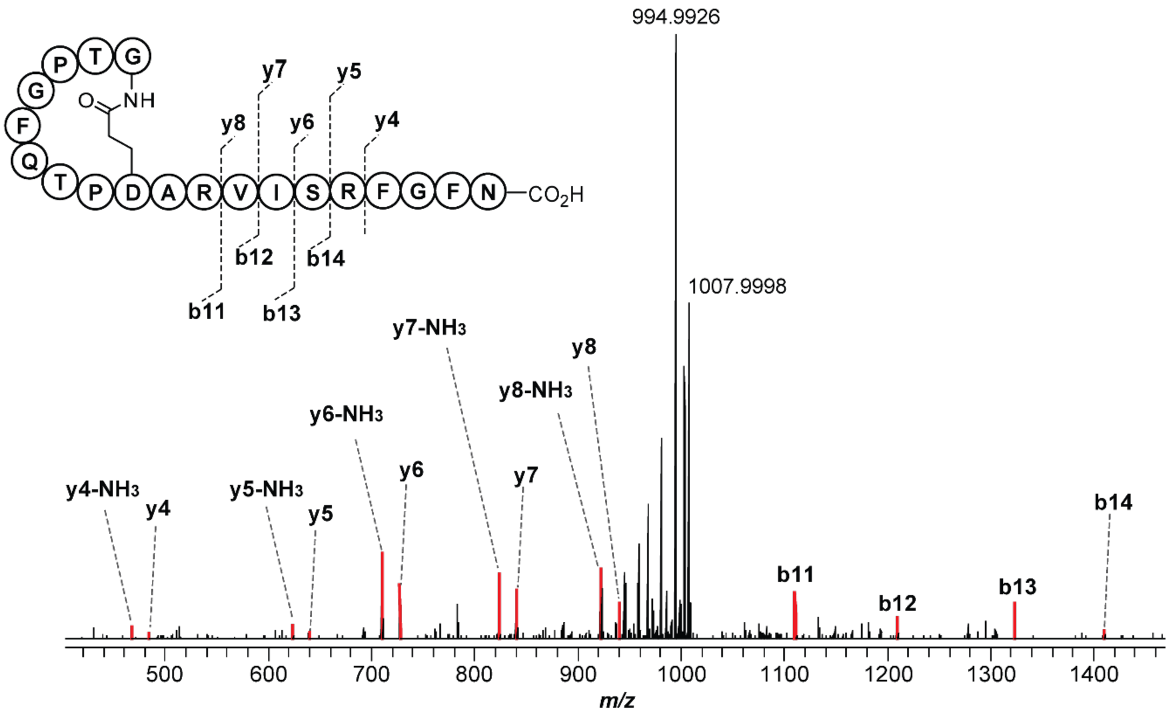

**Figure S3. High-resolution and tandem MS of capistruin produced from CFB. (A)** High-resolution mass spectrum of capistruin produced from CFB with pET41a-*capABCD* as the DNA template. **(B)** Mass assignment for the $b^+$ and $y^+$ ions generated from the CID of capistruin. **(C)** Tandem MS spectrum of capistruin. The most intense signals (*m/z* 994.9926 and 1007.9998) correspond to the neutral loss of multiple molecules of water and/or ammonia, consistent with previous reports on capistruin.[4,17] Assigned b and y ions are red.

A

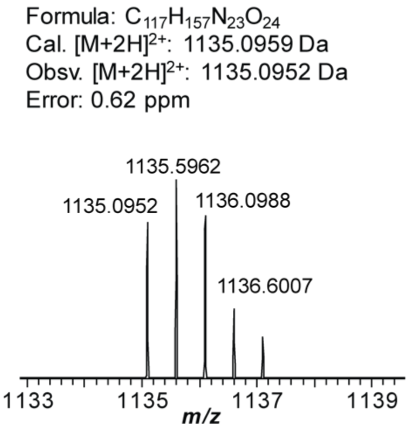

B

| Ion | Calculated mass (Da) | Observed mass (Da) | Error (ppm) |
|---|---|---|---|
| b9 | 1118.4947 | 1118.4922 | 2.24 |
| b10 | 1231.5788 | 1231.5760 | 2.27 |
| b11 | 1344.6628 | 1344.6600 | 2.08 |
| b12 | 1491.7313 | 1491.7286 | 1.81 |
| b13 | 1590.7997 | 1590.7969 | 1.76 |
| b14 | 1737.8681 | 1737.8643 | 2.19 |
| y4 | 532.3247 | 532.3234 | 2.44 |
| y5 | 679.3932 | 679.3917 | 2.21 |
| y6 | 778.4616 | 778.4600 | 2.06 |
| y7 | 925.5300 | 925.5283 | 1.84 |
| y8 | 1038.6140 | 1038.6118 | 2.12 |
| y9 | 1151.6981 | 1151.6952 | 2.52 |

C

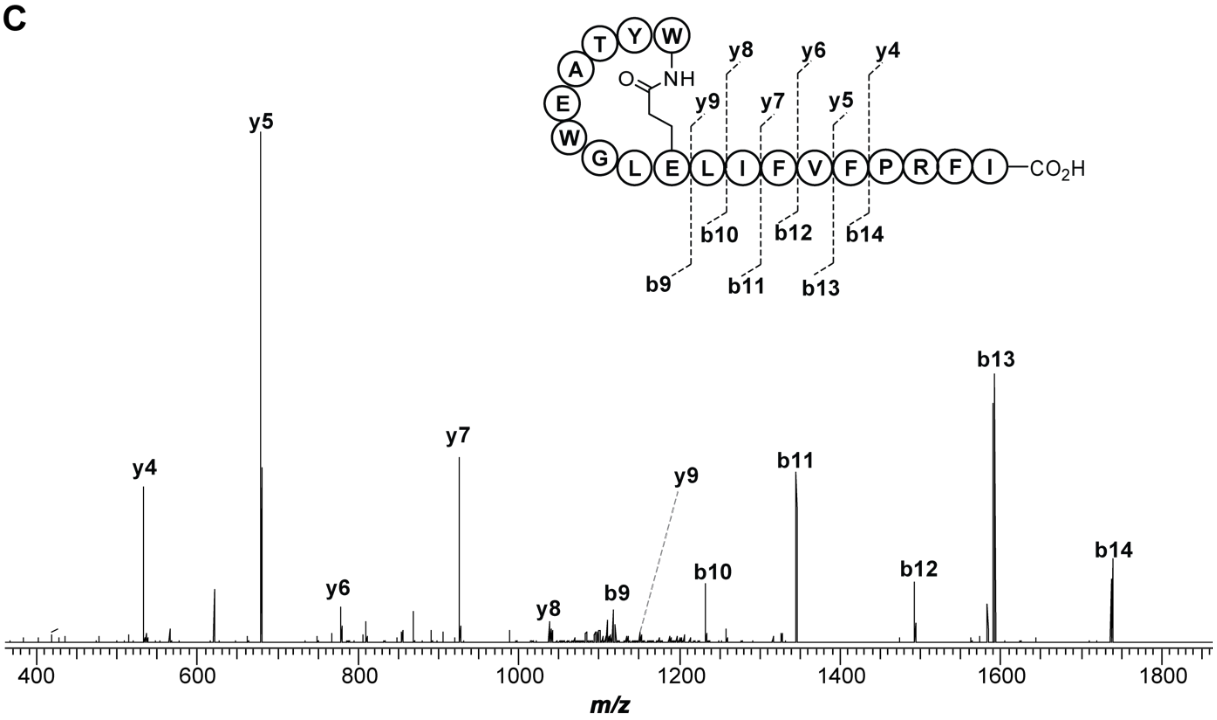

**Figure S4. High-resolution and tandem MS of fusilassin produced from CFB. (A)** High-resolution mass spectrum of fusilassin, produced from CFB with pET28-MBP-*fusA* and pACYC-*fusBE*-*fusC* as the DNA templates. **(B)** Mass assignment for the $b^+$ and $y^+$ ions generated from the CID of fusilassin **(C)** Tandem MS spectrum of fusilassin, consistent with a Trp1Glu9 macrolactam.

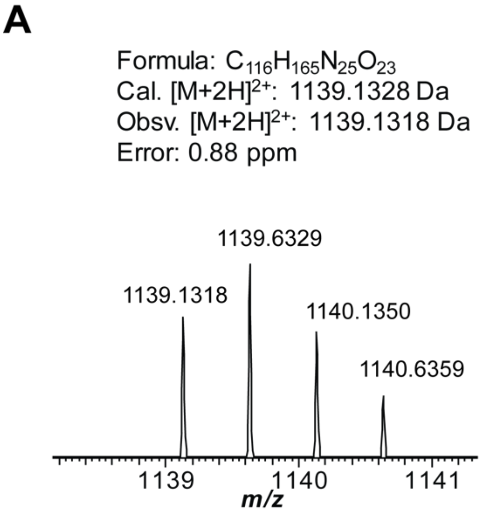

**B**

| Ion | Calculated mass (Da) | Observed mass (Da) | Error (ppm) |
|---|---|---|---|
| b9 | 1080.5631 | 1080.5613 | 1.67 |
| b10 | 1193.6471 | 1193.6454 | 1.42 |
| b11 | 1356.7105 | 1356.7063 | 3.10 |
| b12 | 1469.7945 | 1469.7930 | 1.02 |
| b13 | 1582.8786 | 1582.8760 | 1.64 |
| b14 | 1730.9470 | 1730.9463 | 0.40 |
| y3 | 451.2669 | 451.2660 | 1.99 |
| y4 | 548.3197 | 548.3182 | 2.74 |
| y5 | 695.3881 | 695.3865 | 2.30 |
| y6 | 808.4721 | 808.4708 | 1.61 |
| y7 | 921.5562 | 921.5546 | 1.74 |
| y8 | 1084.6195 | 1084.6152 | 3.96 |
| y9 | 1197.7036 | 1197.7007 | 2.42 |

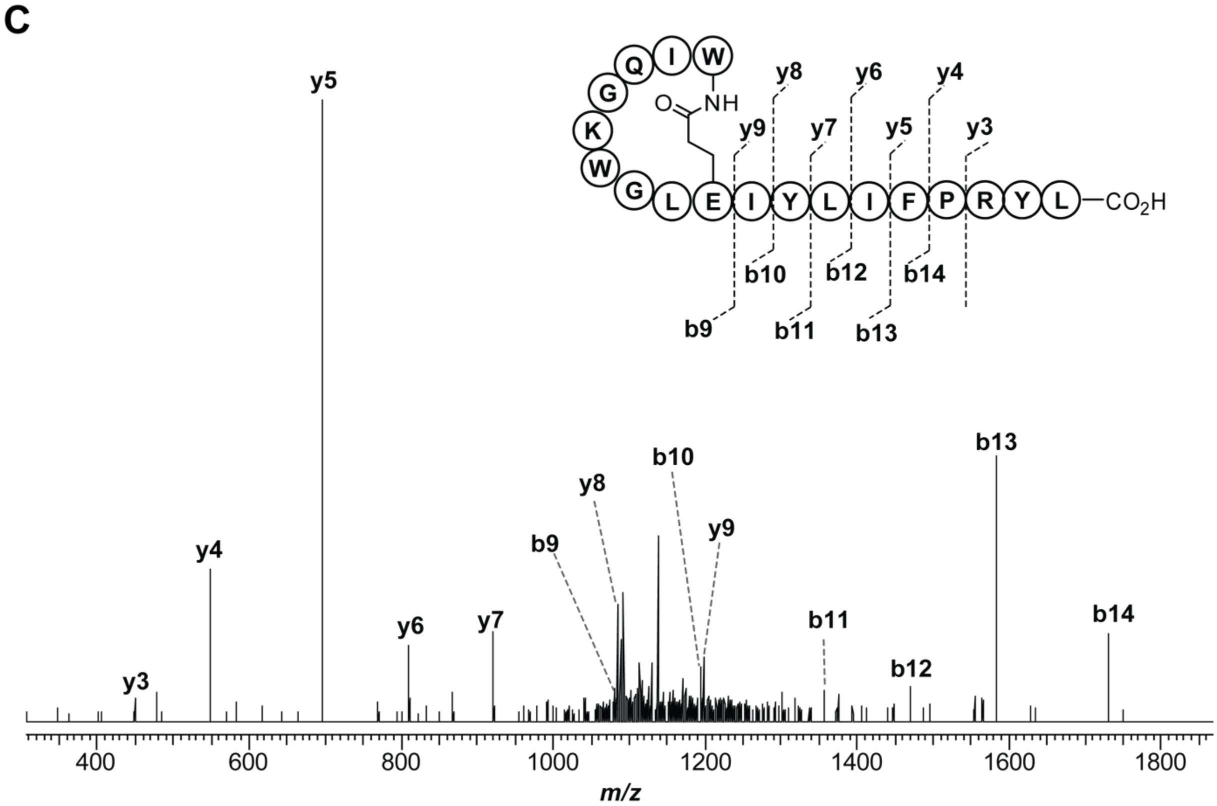

**Figure S5. High-resolution and tandem MS of cellulassin produced from CFB. (A)** High-resolution broadband spectrum of cellulassin, produced from CFB with pET28-MBP-*celACEB* as the DNA template. **(B)** Mass assignment for the $b^+$ and $y^+$ ions generated from the CID of cellulassin. **(C)** Tandem MS spectrum of cellulassin, consistent with a Trp1Glu9 macrolactam.

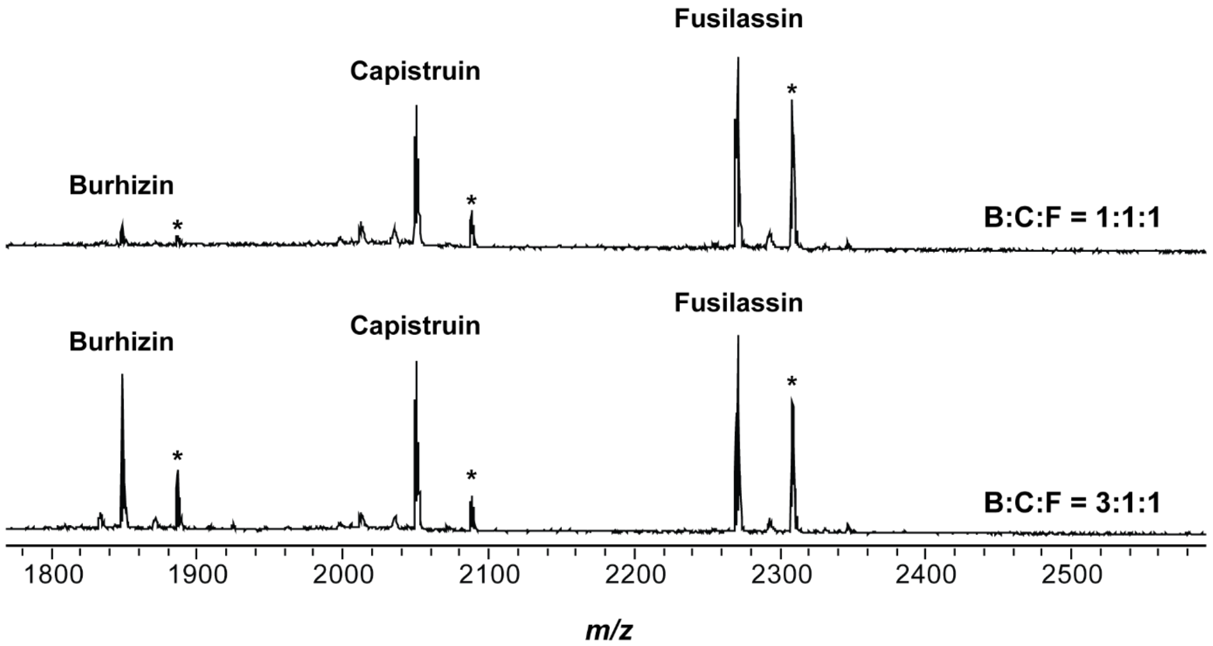

**Figure S6. Simultaneous production of multiple lasso peptides using CFB.** MALDI-TOF mass spectrum of burhizin, capistruin, and fusilassin produced in a single CFB reaction**.** Different ratios of DNA templates encoding the BGC of burhizin, capistruin, and fusilassin were supplied in a single CFB reaction. *Top*, the ratio of DNA templates is 1:1:1 for burhizin, capistruin, and fusilassin respectively; *bottom*, the ratio of DNA templates 3:1:1. B, burhizin; C, capistruin; F, fusilassin. * indicates $[M+K]^+$ ion of the lasso peptide.

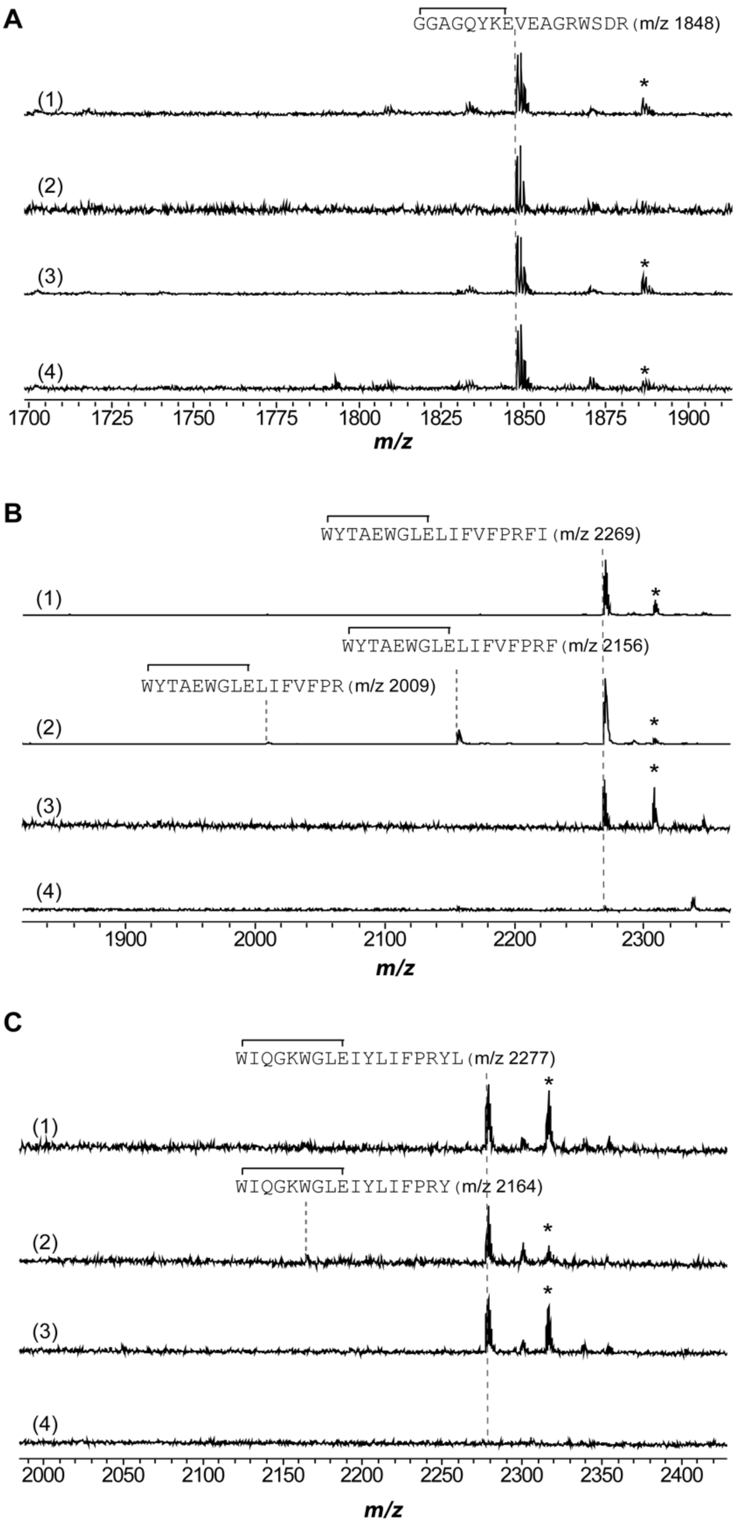

**Figure S7. Carboxypeptidase Y resistance of CFB-produced lasso peptides.** Burhizin **(A)**, fusilassin **(B)**, and cellulassin **(C)** were produced by CFB and shown in the top spectrum (1). Each lasso peptide was subjected to carboxypeptidase Y digestion for 18 h at room temperature (2); heat treatment at 95 °C for 2 h (3); heat treatment at 95 °C for 2 h followed by carboxypeptidase Y digestion for 18 h at room temperature (4). * indicates $[M+K]^+$ ion of the lasso peptide.

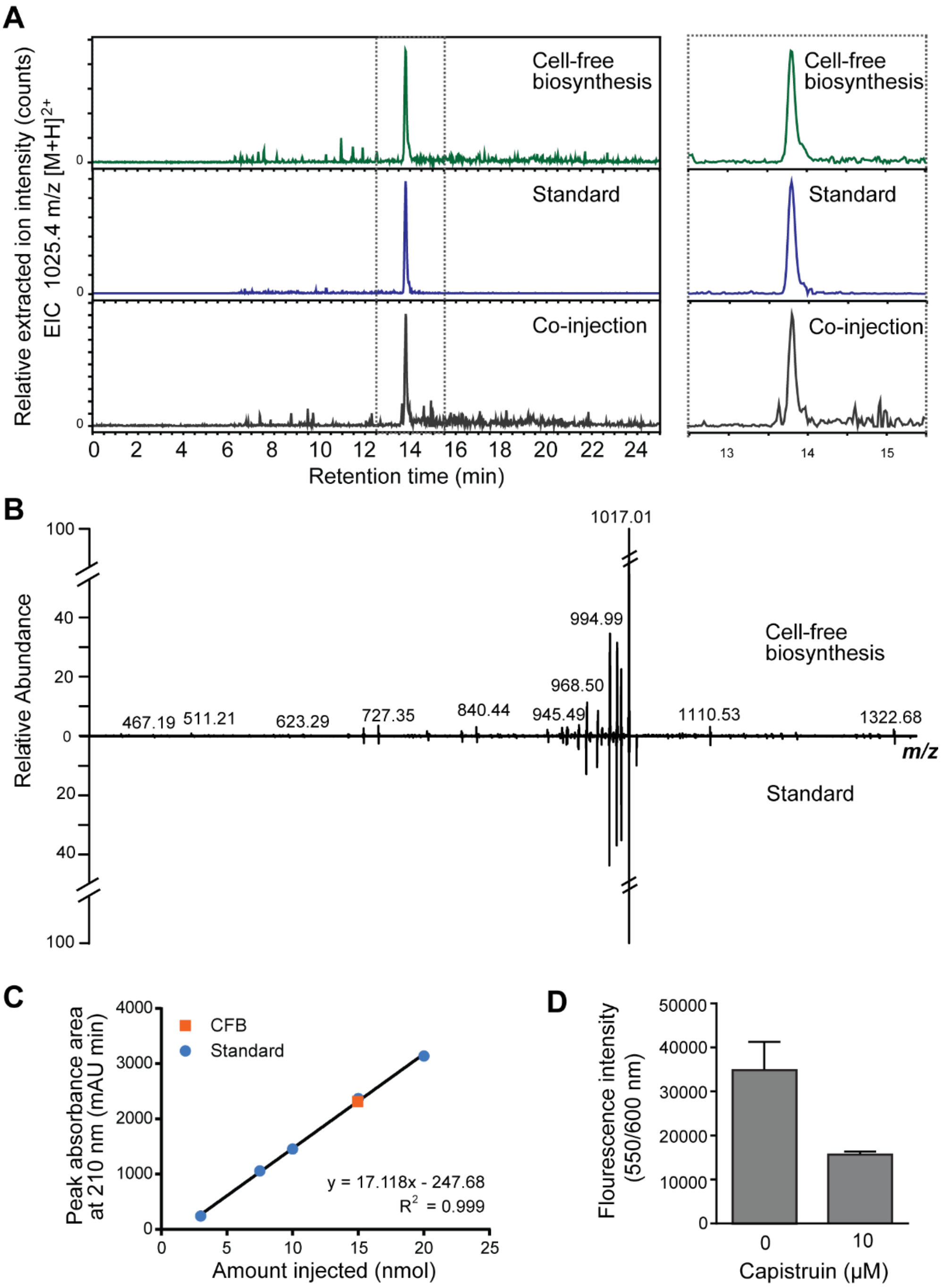

**Figure S8. Purification and yield determination of capistruin produced from CFB.** (A) LC-MS confirms CFB-produced capistruin is identical to an authentic sample.[4] Extracted ion chromatographs for *m/z* 1025.4 ($[M+2H]^{2+}$ ion of capistruin) are shown for samples of capistruin isolated from CFB (green), a standard isolated from heterologous expression of capistruin (blue), and a co-injection. (B) High-resolution tandem mass spectrum of capistruin from CFB (top) compared to the capistruin standard (bottom). (C) Quantification of capistruin from a 750 µL CFB reaction. The area of the 220 nm absorbance peak corresponding to capistruin was compared to a standard curve to determine yield. Orange and square, CFB produced capistruin; blue and circle, standard capistruin (D) Capistruin isolated from CFB inhibited the CFB-based production of mCherry (presumably by inhibition of RNA polymerase).

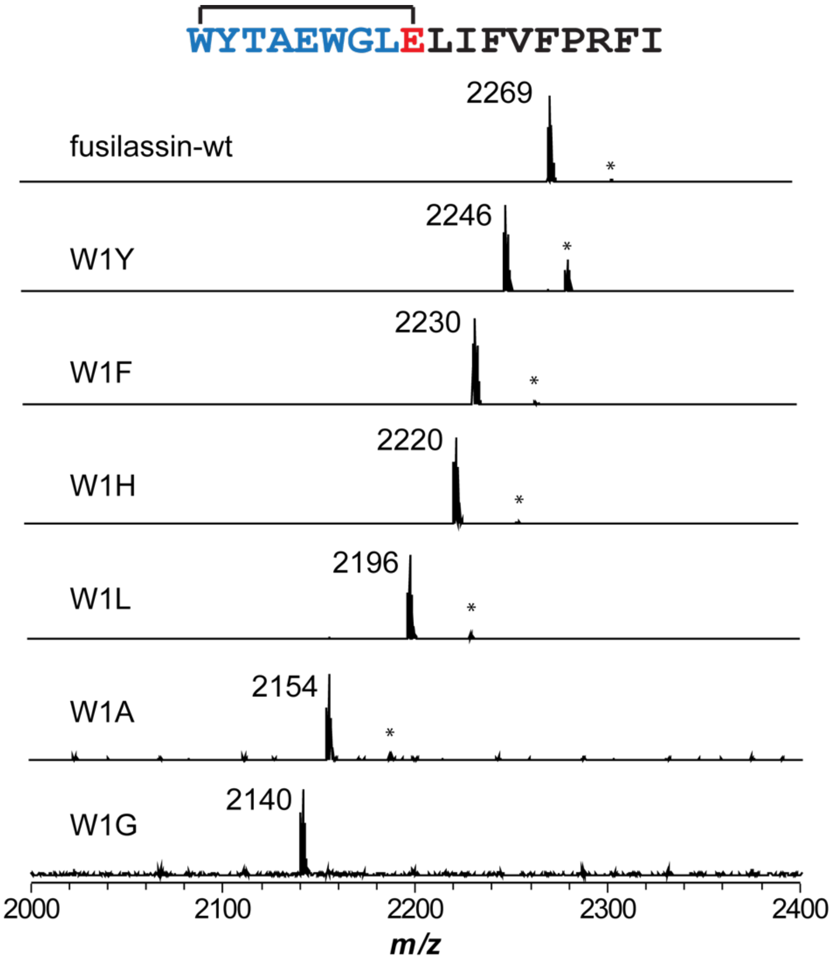

**Figure S9. MALDI-TOF-MS of Fusilassin variants with Trp1 substitutions.** Endpoint MALDI-TOF-MS assay of fusilassin variants produced from CFB with different substitutions on the first position. The mass label corresponds to the $[M+H]^+$ ion of the lasso peptide. * indicates $[M+K]^+$ ion of lasso peptide.

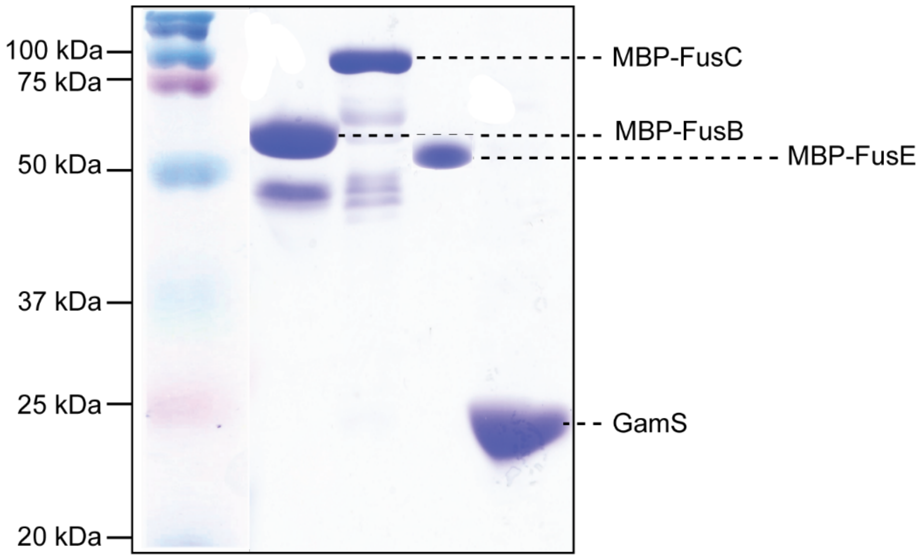

**Figure S10. SDS-PAGE gel of protein used in this study.** From left to right: maltose-binding protein (MBP)-tagged FusB; MBP-FusC; MBP-FusE; 6xHis-GamS. All four proteins were heterologously expressed in *E. coli* and purified to homogeneity by amylose affinity chromatography (MBP-FusB, MBP-FusC, MBP-FusE) or Ni-NTA affinity chromatography (6xHis-GamS).

**A**

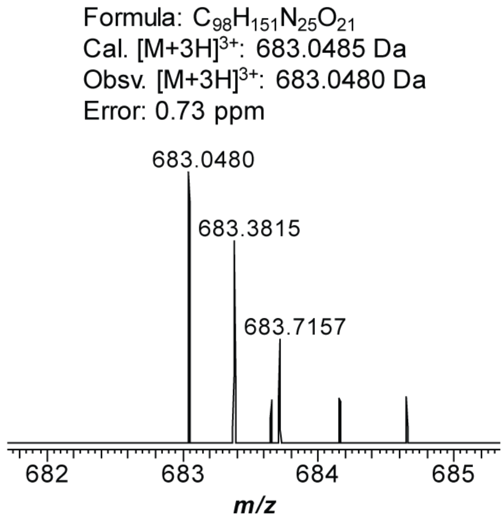

**B**

| Ion | Calculated mass (Da) | Observed mass (Da) | Error (ppm) |
|---|---|---|---|
| b8 | 873.4583 | 873.4557 | 2.98 |
| b9 | 986.5423 | 986.5410 | 1.32 |
| b10 | 1172.6217 | 1172.6191 | 2.22 |
| b11 | 1285.7057 | 1285.7007 | 3.89 |
| $[b12+2H]^{2+}$ | 716.8904 | 716.8895 | 1.26 |
| $[b13+2H]^{2+}$ | 773.4325 | 773.4316 | 1,16 |
| y4 | 502.2812 | 502.2800 | 2.39 |
| y5 | 615.3652 | 615.3638 | 2.28 |
| y6 | 762.4336 | 762.4321 | 1.84 |
| y7 | 875.5177 | 875.5158 | 2.40 |
| y8 | 1061.5970 | 1061.5938 | 3.01 |
| y9 | 1174.6811 | 1174.6787 | 2.04 |

**C**

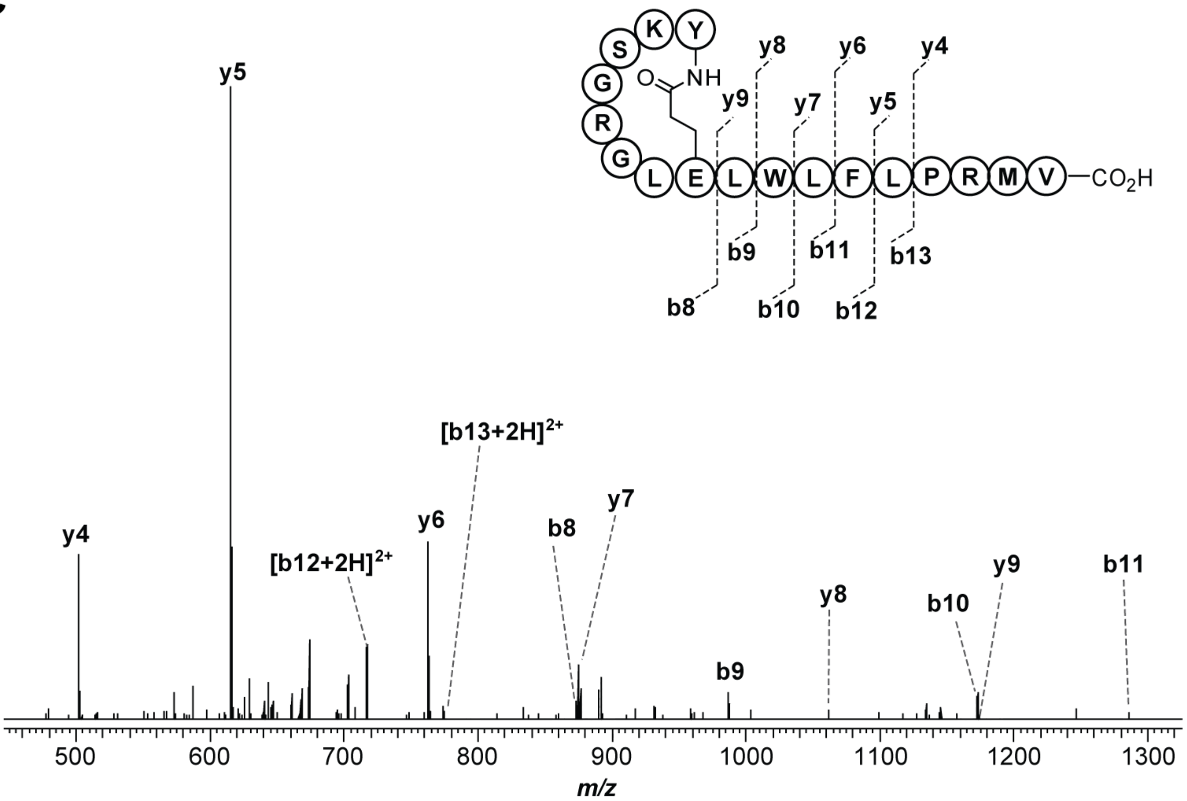

**Figure S11. High-resolution and tandem MS of halolassin produced from CFB using chimeric substrate FusA$_{LP}$-HalA$_{CP}$ as the DNA template. (A)** High-resolution broadband spectrum of halolassin, produced from CFB with FusA$_{LP}$-HalA$_{CP}$ as the DNA template and reacted with heterologously expressed and purified FusB, C, and E. **(B)** Mass assignment for the $b^+$ and $y^+$ ions generated from the CID of halolassin. **(C)** Tandem MS spectrum of halolassin, consistent with a Tyr1-Glu8 macrolactam.

A

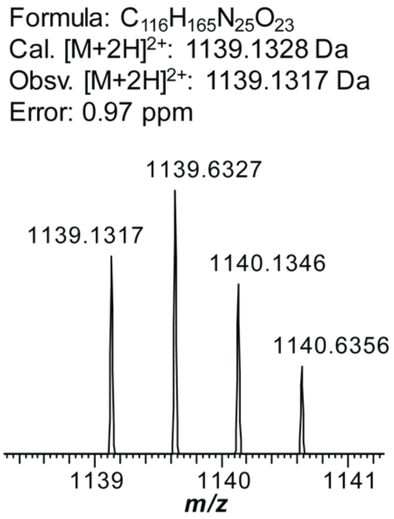

B

| Ion | Calculated mass (Da) | Observed mass (Da) | Error (ppm) |
|---|---|---|---|
| b9 | 1080.5631 | 1080.5602 | 2.68 |
| b10 | 1193.6471 | 1193.6444 | 2.18 |
| b11 | 1356.7105 | 1356.7063 | 3.10 |
| b12 | 1469.7945 | 1469.7915 | 2.04 |
| b13 | 1582.8786 | 1582.8748 | 2.40 |
| b14 | 1730.9470 | 1730.9447 | 1.33 |
| y4 | 548.3197 | 548.3183 | 2.55 |
| y5 | 695.3881 | 695.3863 | 2.59 |
| y6 | 808.4721 | 808.4703 | 2.23 |
| y7 | 921.5562 | 921.5541 | 2.28 |
| y8 | 1084.6195 | 1084.6167 | 2.58 |
| y9 | 1197.7036 | 1197.7007 | 2.42 |

C

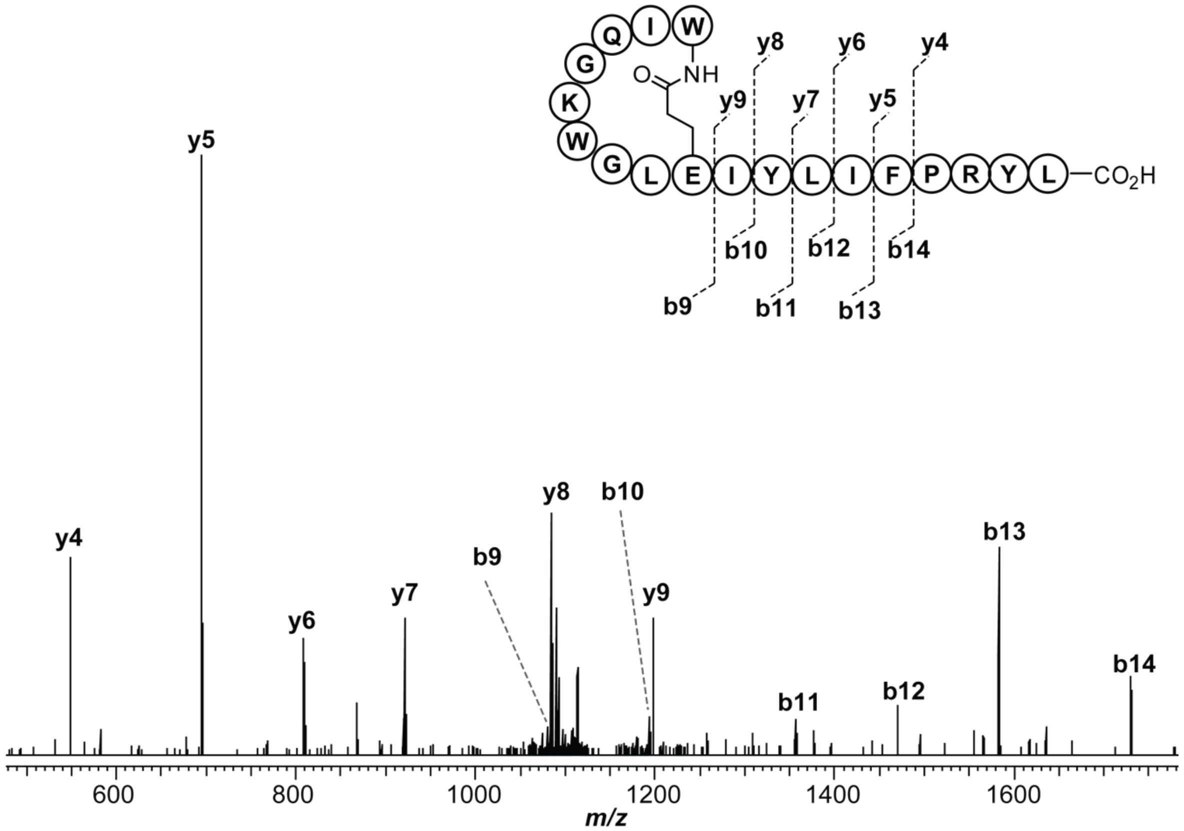

**Figure S12. High-resolution and tandem MS of cellulassin produced from CFB using chimeric substrate FusA$_{LP}$-CelA$_{CP}$ as the DNA template.** **(A)** High-resolution broadband spectrum of cellulassin, produced from CFB with FusA$_{LP}$-HalA$_{CP}$ as the DNA template and reacted with heterologously expressed and purified FusB, C, and E. **(B)** Mass assignment for the $b^+$ and $y^+$ ions generated from the CID of cellulassin. **(C)** Tandem MS spectrum of cellulassin, consistent with a Trp1-Glu9 macrolactam.

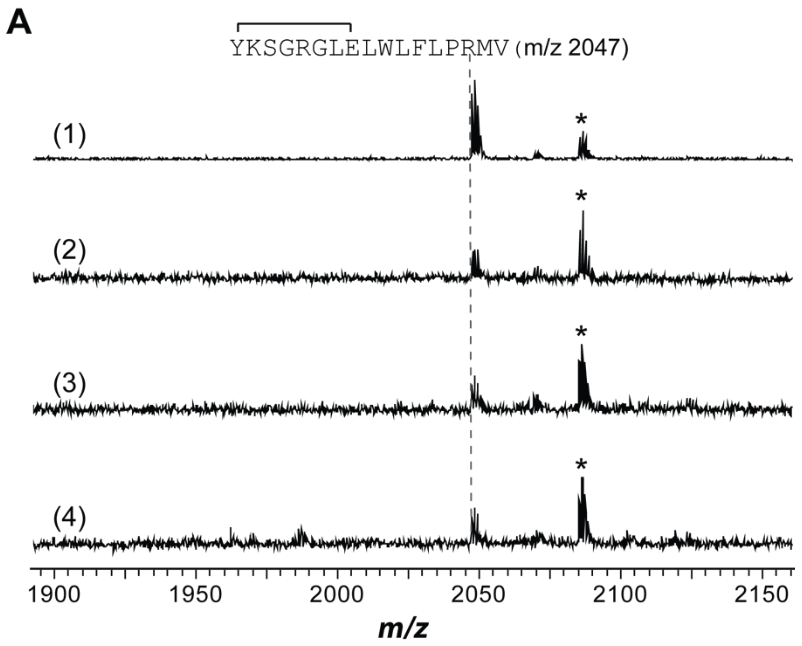

**Figure S13. Carboxypeptidase Y treatment of halolassin and cellulassin produced with chimeric substrates from CFB.** Halolassin and cellulassin were produced through CFB using chimeric precursors as the DNA template (1). The produced lasso peptides were treated under 3 different conditions: carboxypeptidase Y digestion for 18 h (2); heat treatment at 95 °C for 2 h (3); heat treatment at 95 °C for 2 h followed by carboxypeptidase Y digestion for 18 h at room temperature (4). The presence/absence of intact lasso peptide upon treatment under condition (4) and the presence of intact lasso peptide under conditions (2) and (3) indicate the threaded topology of a lasso peptide. ^ indicates the uncyclized linear core peptide. * indicates the $[M+K]^+$ ion of the lasso peptide.

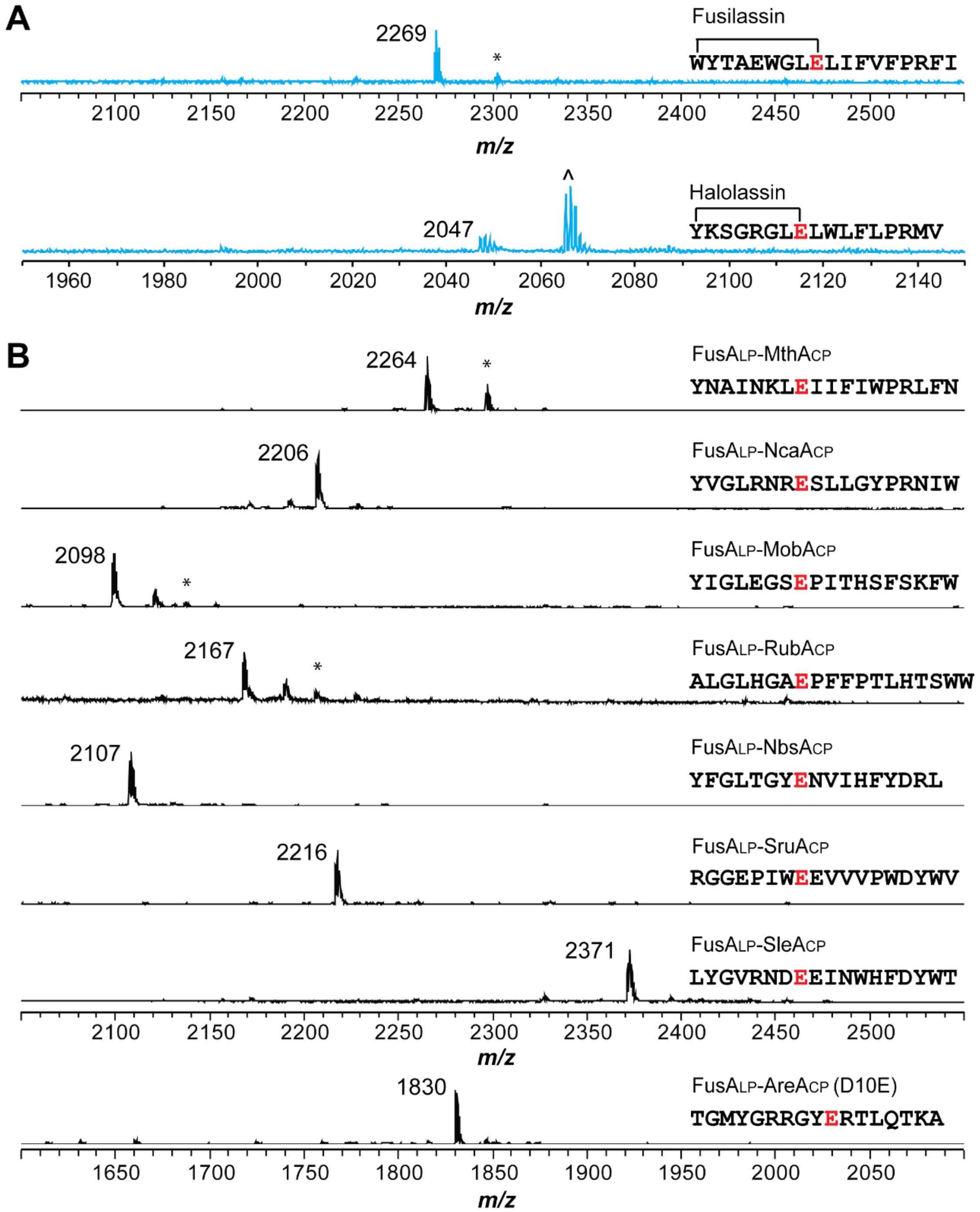

**Figure S14. Chimeric substrate processing in PURExpress reactions. (A)** Wildtype FusA and FusA$_{LP}$-HalA$_{CP}$ were synthesized using PURExpress and treated with purified FusB, FusC, and FusE. Analysis by MALDI-TOF-MS demonstrated the production of mature lasso peptides (blue) fusilassin (*m/z* 2269), and to a lesser extent, halolassin (*m/z* 2047). Mass labels corresponds to the $[M+H]^+$ ion of the lasso peptide. ^ indicates the uncyclized linear core. * indicates the $[M+K]^+$ of the lasso peptide. **(B)** Eight chimeric substrates were produced by PURExpress and reacted with purified FusB, FusC, and FusE. MALDI-TOF-MS detected masses corresponding to the $[M+H]^+$ ions for the linear core peptides, indicating all substrates were processed by FusB/E but none were processed by FusC. * indicates $[M+K]^+$ ion for the linear core peptide.

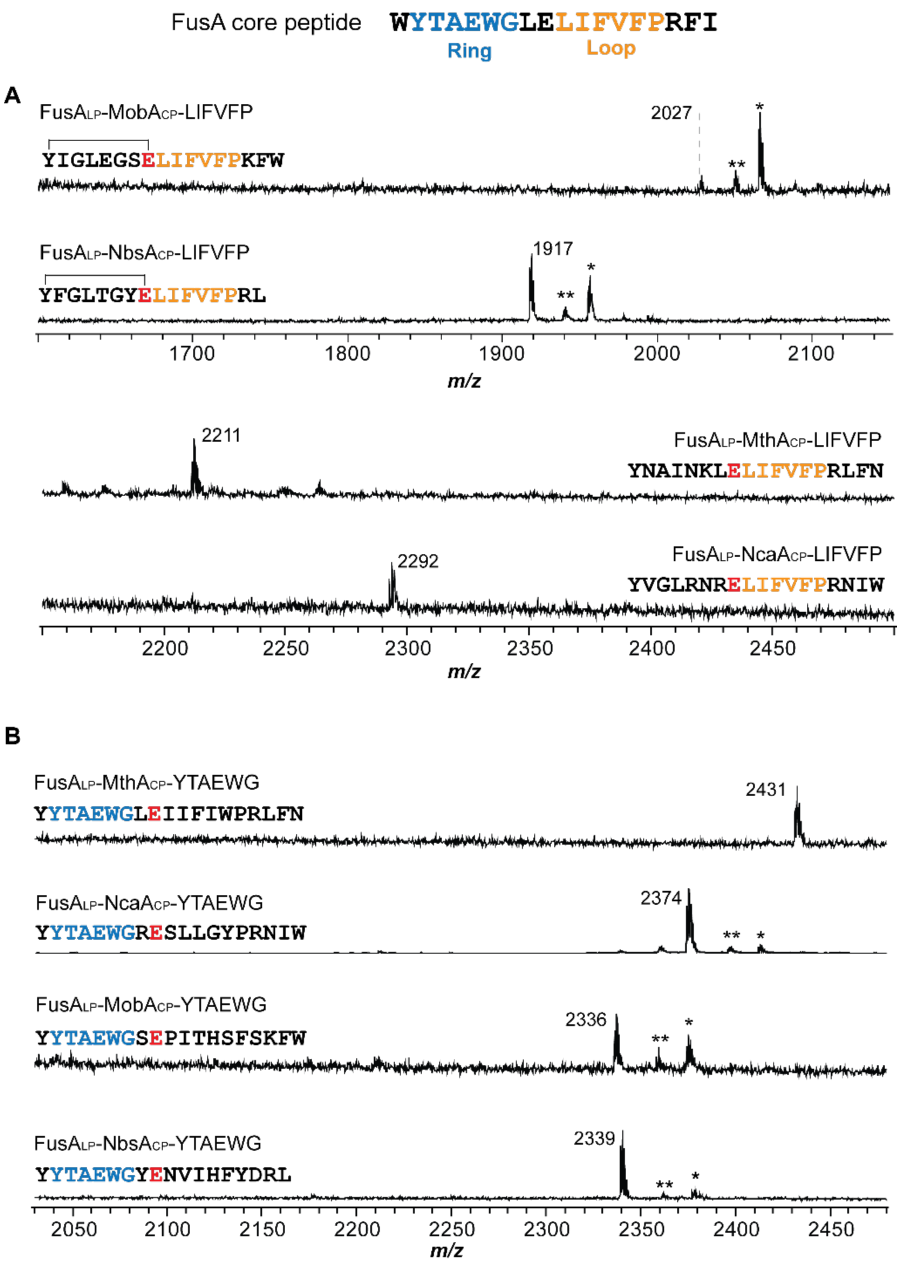

**Figure S15. Replacement of the ring/loop sequences of chimeric substrates with FusA ring/loop sequence. (A)** Four new chimeric substrates, in which the loop region of the core sequence was replaced with FusA loop (orange), were produced by CFB and reacted with purified FusB, FusC, and FusE. Cyclized FusA$_{LP}$-MobA$_{CP}$-LIFVFP (*m/z* 2027) and FusA$_{LP}$-NbsA$_{CP}$-LIFVFP (*m/z* 1917) were detected, while the other two failed to cyclize. FusA$_{LP}$-MthA$_{CP}$-LIFVFP and FusA$_{LP}$-NcaA$_{CP}$-LIFVFP were produced through PURExpress and reacted with FusB, FusC, and FusE. Their corresponding linear core peptides were detected by MALDI (*m/z* of 2211 and 2292, respectively). **(B)** Four additional chimeric substrates, in which a portion of the ring region was replaced with FusA ring sequence (blue), were also prepared. All four new chimeric substrates failed to cyclize in CFB and their linear core peptides were produced in PURExpress and detected by MALDI (*m/z* of 2431, 2347, 2336, and 2339, respectively). * indicates the $[M+K]^+$ ion of cyclized lasso peptide/uncyclized core peptide, ** indicates the $[M+Na]^+$ ion of cyclized lasso peptide/uncyclized core peptide.

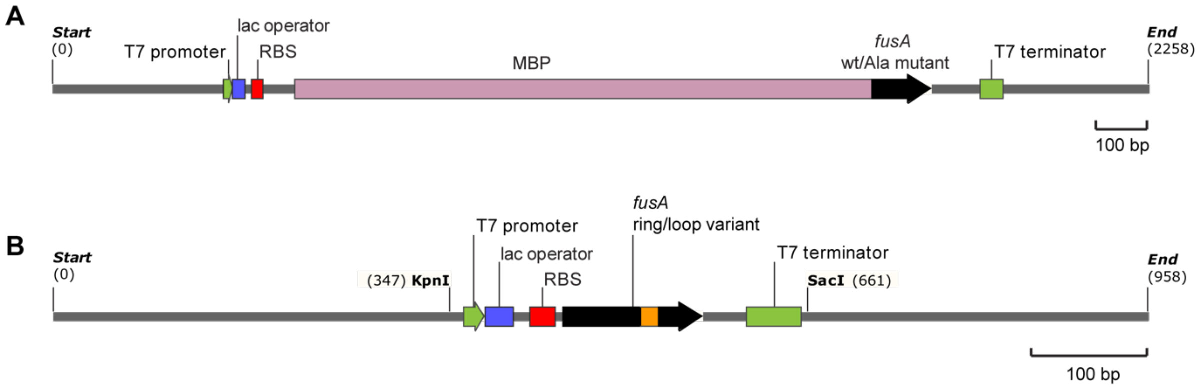

**Figure S16. Linear DNA template used in CFB reactions. (A)** The linear DNA template of *fusA* wild type and Ala-substituted variants used in CFB reactions. **(B)** The linear DNA template of *fusA* ring/loop variants used for CFB reactions. T7 promotor and T7 terminator is shown in green; lac operators are shown in blue; RBSs are shown in red; *fusA* genes are shown in black; MBP, maltose-binding protein, is shown in pink.

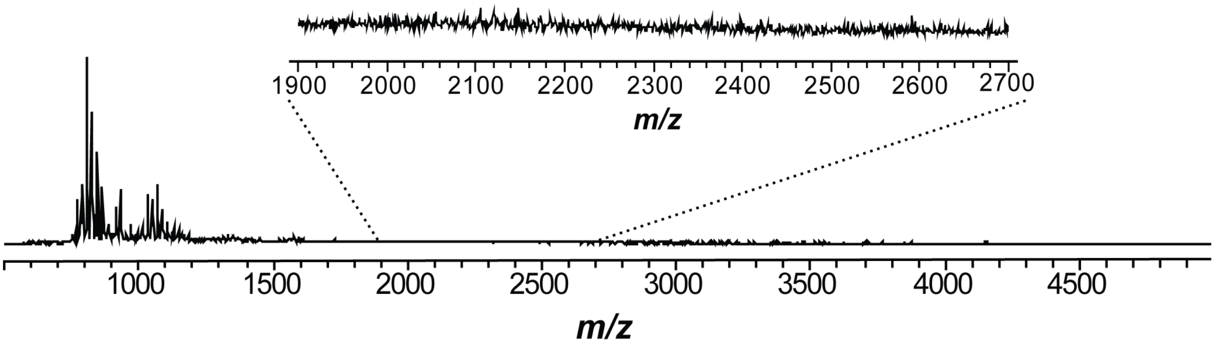

**Figure S17. CFB reaction without DNA template.** The control reaction was conducted the same way as other CFB reactions including fusilassin biosynthetic proteins, but with no DNA template. No signal was present in the mass range of 1900−2700 Da.

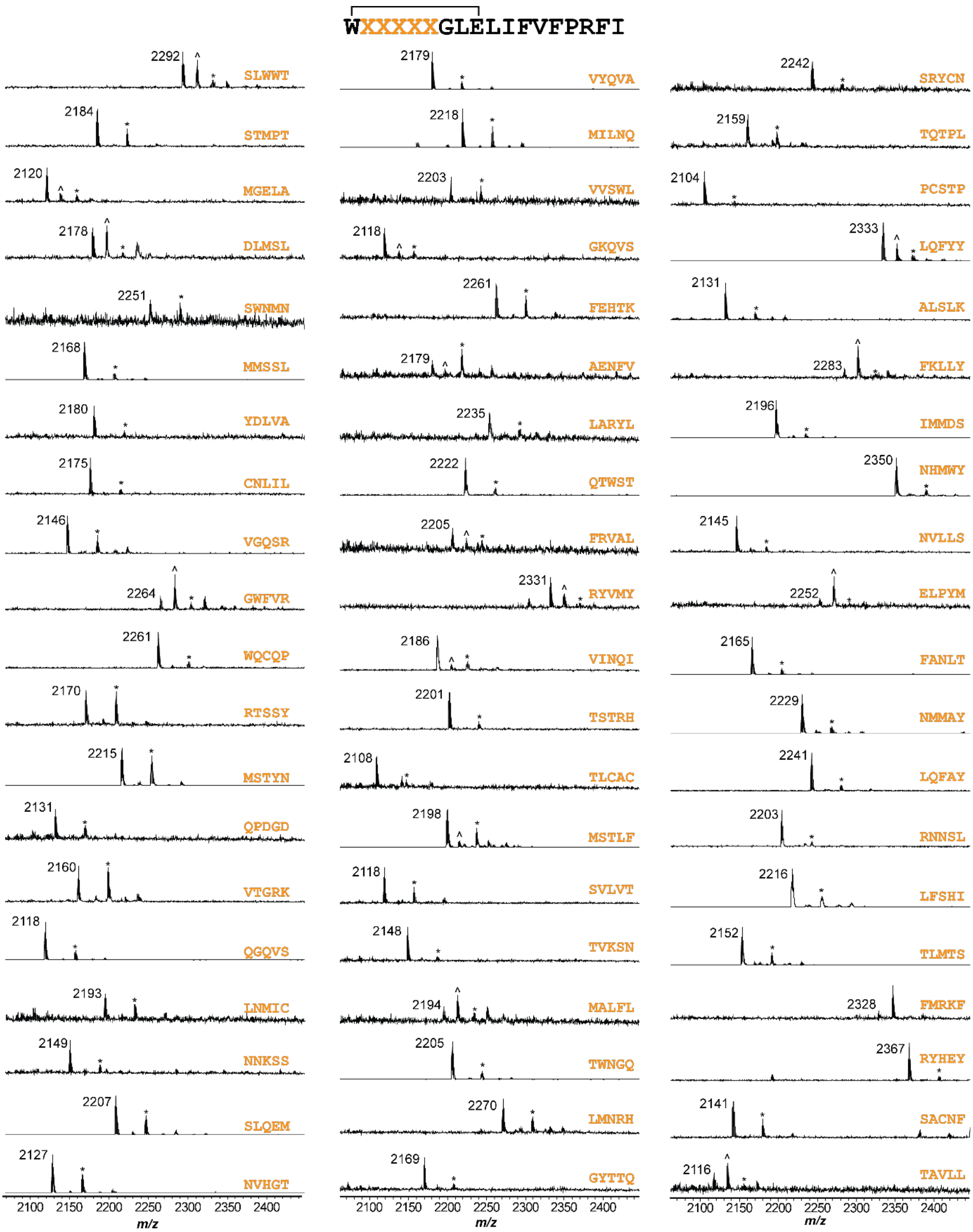

**Figure S18. MALDI-TOF mass spectra of the fusilassin ring variants produced from CFB.** Fusilassin variants (270 out of 513; an additional 10 are shown in main text **Figure 4**) detected by MALDI-TOF-MS are shown. The DNA encoding each ring variant product was subjected to Sanger sequencing. Orange depicts the varied sequence of positions 2–6 of the FusA core region. The mass label corresponds to the $[M+H]^+$ ion of the lasso peptide. * indicates $[M+K]^+$ ion for the lasso peptide. ^ indicates the uncyclized linear core peptide. Figure continues onto the next 4 pages.

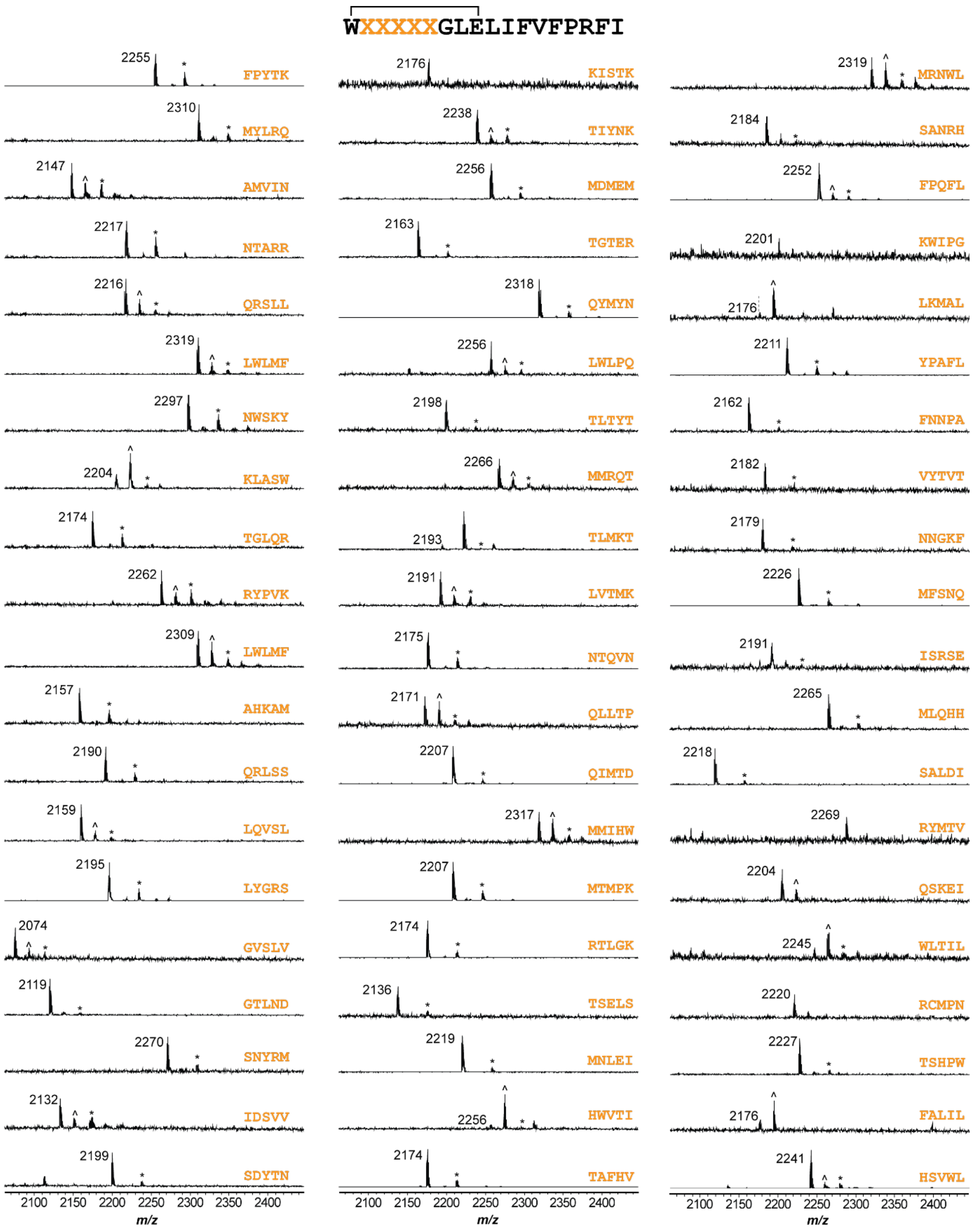

**Figure S18.** *Continued from previous page (2 of 5)*

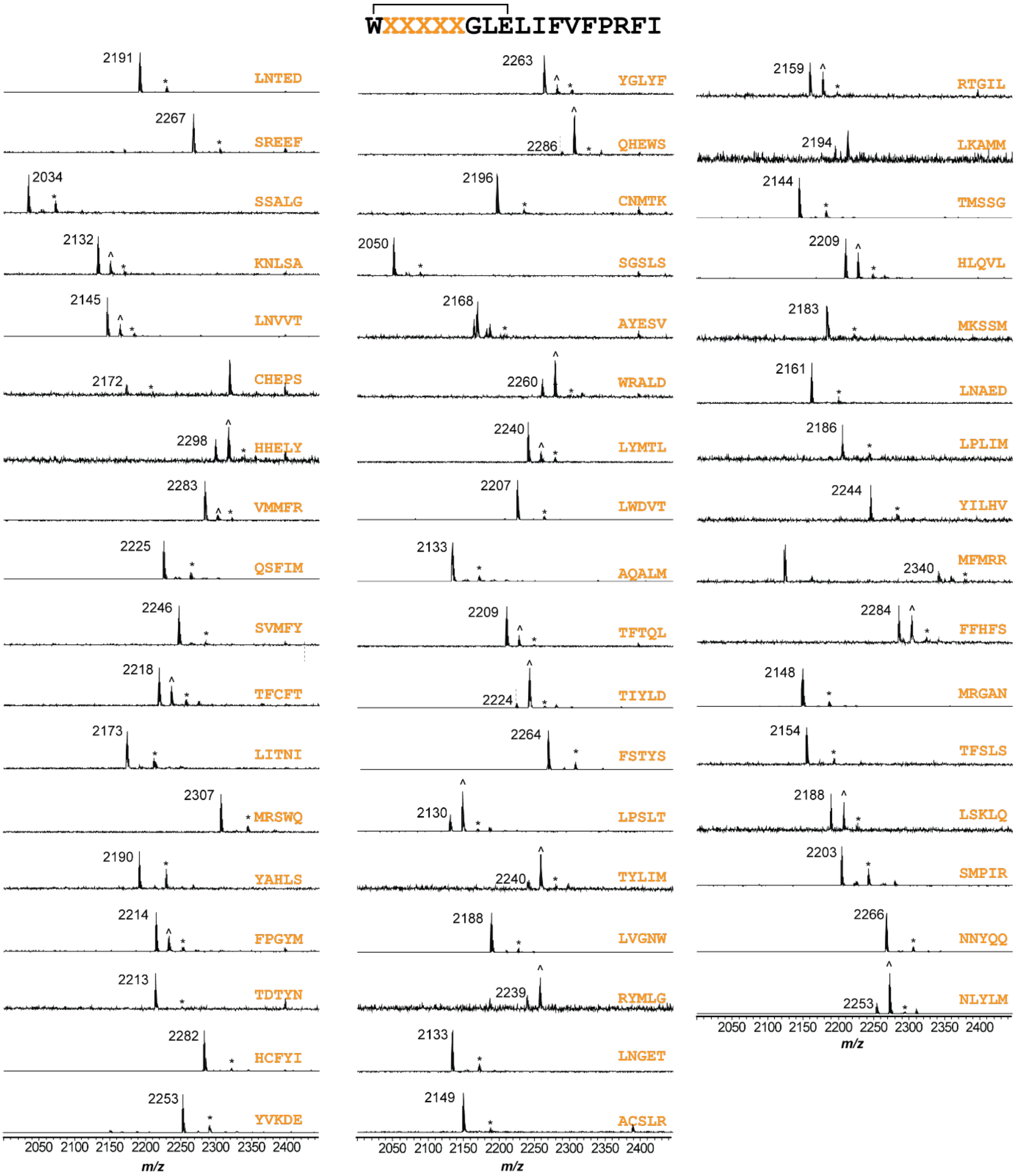

**Figure S18.** *Continued from previous page (3 of 5)*

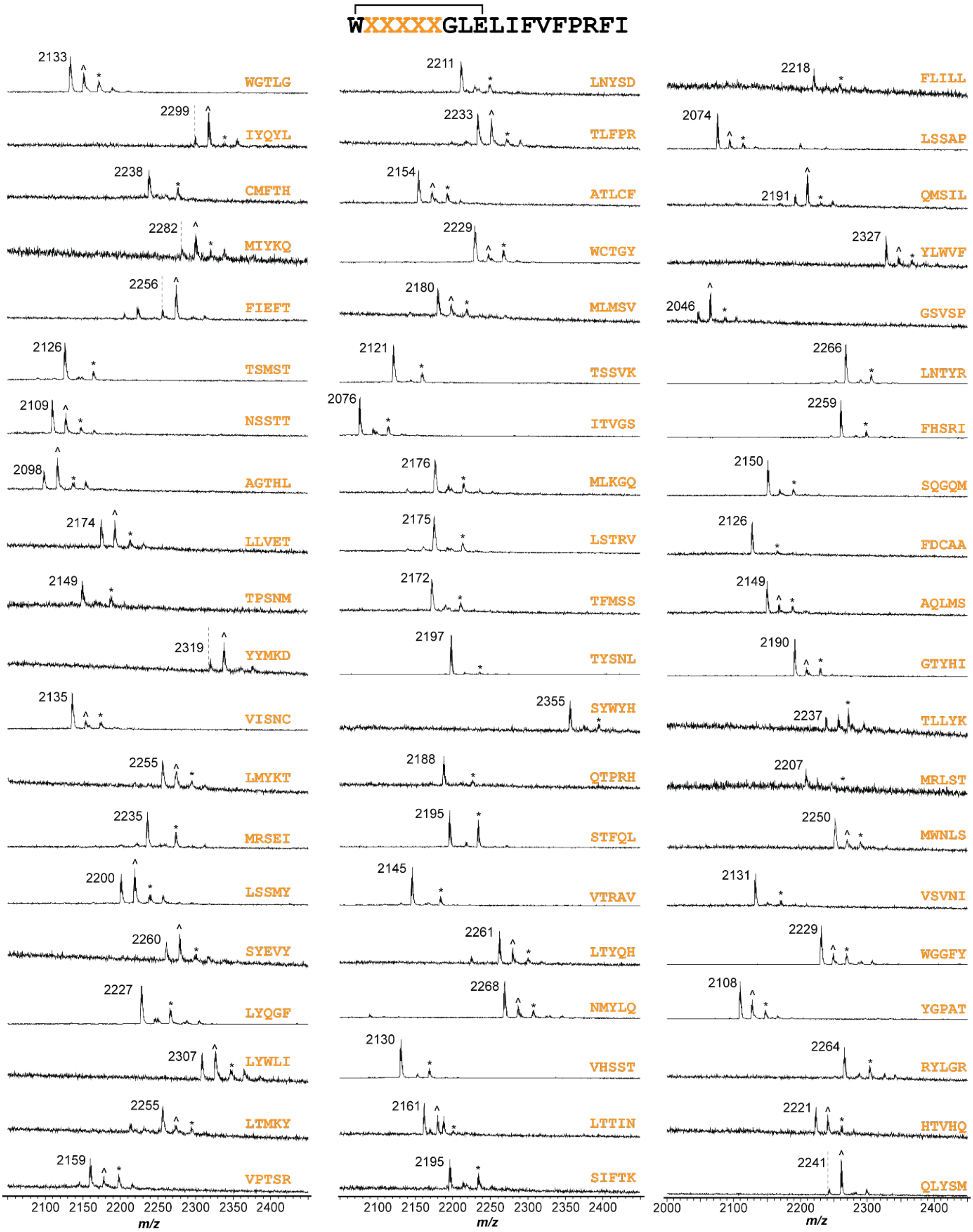

**Figure S18.** *Continued from previous page (4 of 5)*

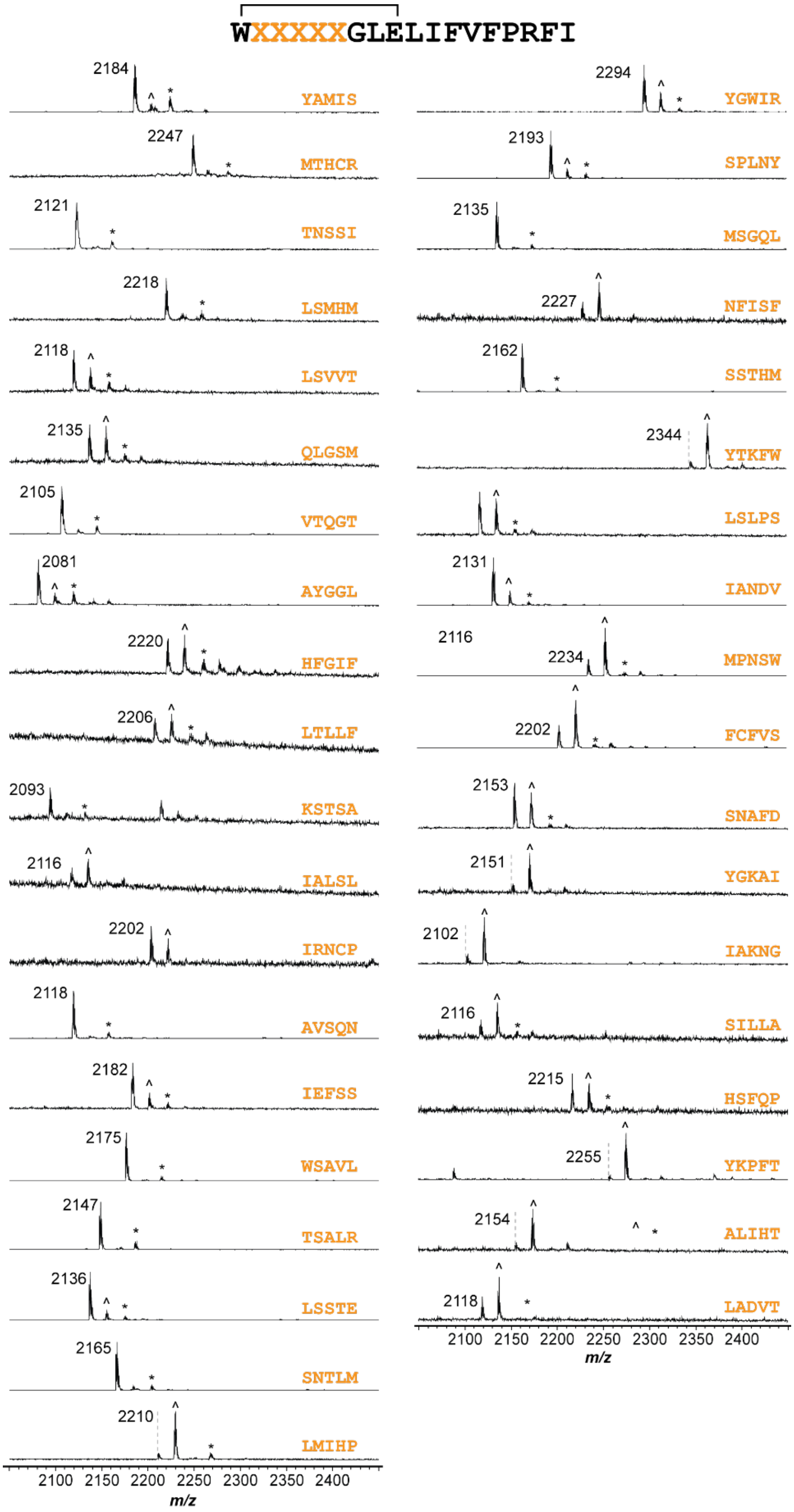

**Figure S18.** *Continued from previous page (5 of 5)*

**A**

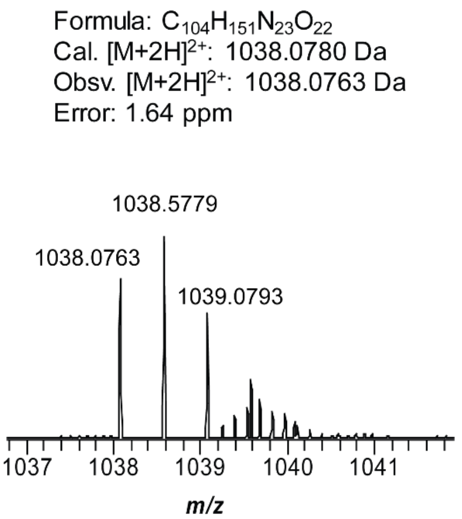

**B**

| Ion | Calculated mass (Da) | Observed mass (Da) | Error (ppm) |
|---|---|---|---|
| b9 | 924.4579 | 924.4562 | 1.84 |
| b10 | 1037.5420 | 1037.5397 | 2.22 |
| b11 | 1150.6261 | 1150.6235 | 2.26 |
| b12 | 1297.6945 | 1294.6917 | 2.16 |
| b13 | 1396.7629 | 1396.7603 | 1.86 |
| b14 | 1543.8313 | 1543.8287 | 1.68 |
| b15 | 1640.8841 | 1640.8810 | 1.89 |
| b16 | 1796.9852 | 1796.9816 | 2.00 |
| y4 | 532.3247 | 532.3235 | 2.25 |
| y5 | 679.3932 | 679.3917 | 2.21 |
| y6 | 778.4616 | 778.4600 | 2.06 |
| y7 | 925.5300 | 925.5282 | 1.94 |
| y8 | 1038.6140 | 1038.6118 | 2.12 |
| y9 | 1151.6981 | 1151.6957 | 2.08 |

**C**

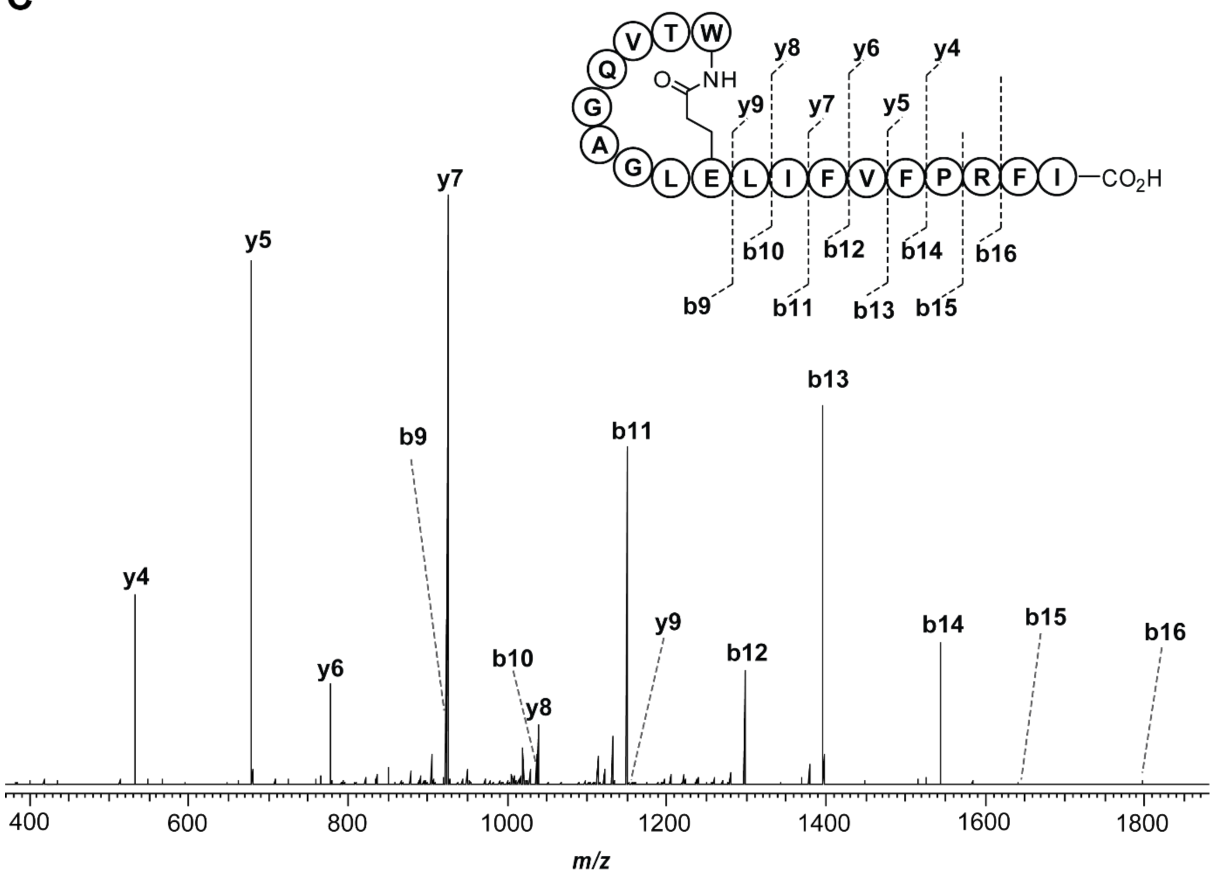

**Figure S19. High-resolution and tandem MS of TVQGA variant produced from CFB. (A)** High-resolution broadband spectrum of fusilassin TVQGA variant, produced from CFB. **(B)** Mass assignment for the $b^+$ and $y^+$ ions generated from the CID of fusilassin TVQGA variant. **(C)** Tandem MS spectrum of fusilassin TVQGA variant, consistent with a Trp1Glu9 macrolactam.

A

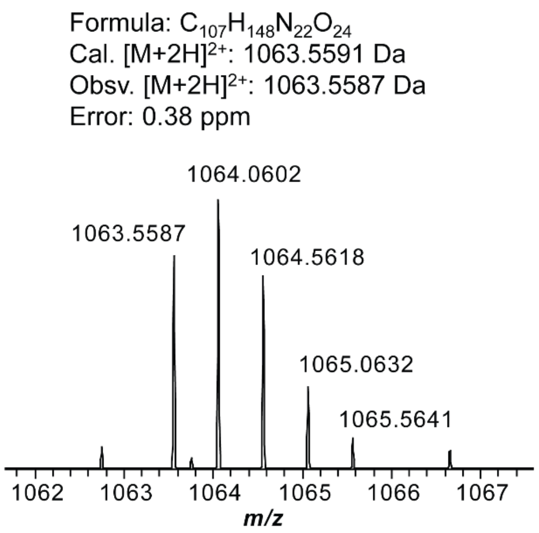

B

| Ion | Calculated mass (Da) | Observed mass (Da) | Error (ppm) |
|---|---|---|---|
| b9 | 975.4212 | 975.4202 | 1.03 |
| b10 | 1088.5042 | 1088.5039 | 0.28 |
| b11 | 1201.5893 | 1201.5879 | 1.17 |
| b12 | 1348.6578 | 1348.6564 | 1.04 |
| b13 | 1447.7262 | 1447.7253 | 0.62 |
| b14 | 1594.7946 | 1594.7936 | 0.63 |
| y4 | 532.3247 | 532.3240 | 2.44 |
| y5 | 679.3932 | 679.3923 | 2.21 |
| y6 | 778.4616 | 778.4606 | 2.06 |
| y7 | 925.5300 | 925.5289 | 1.84 |
| y8 | 1038.6140 | 1038.6128 | 2.12 |
| y9 | 1151.6981 | 1151.6968 | 2.52 |

C

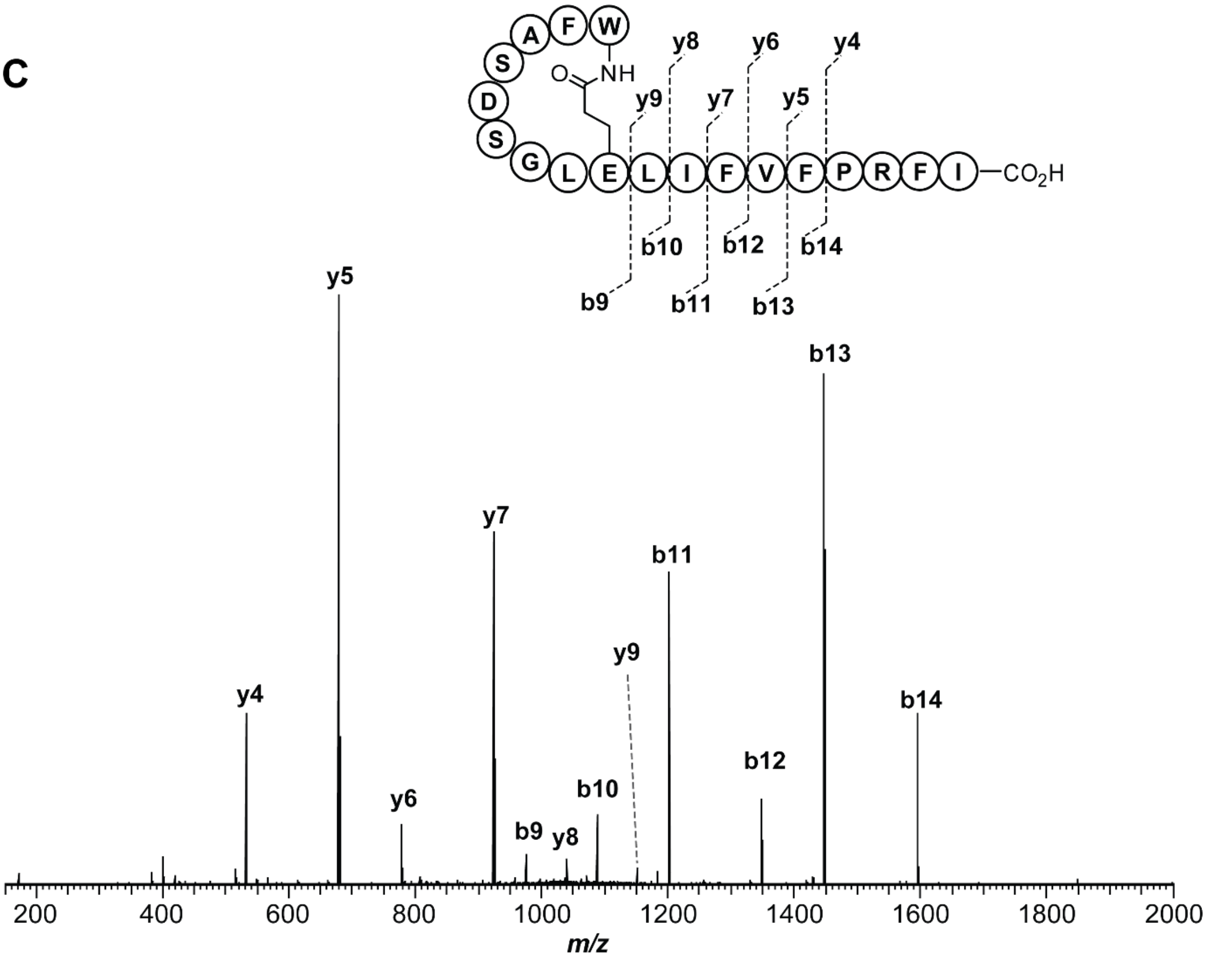

**Figure S20. High-resolution and tandem MS of FASDS variant produced from CFB. (A)** High-resolution broadband spectrum of fusilassin FASDS variant, produced from CFB. **(B)** Mass assignment for the $b^+$ and $y^+$ ions generated from the CID of fusilassin FASDS variant. **(C)** Tandem MS spectrum of fusilassin FASDS variant, consistent with a Trp1Glu9 macrolactam.

A

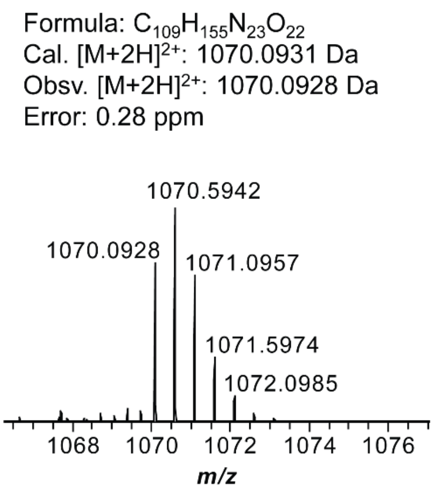

B

| Ion | Calculated mass (Da) | Observed mass (Da) | Error (ppm) |
|---|---|---|---|
| b9 | 988.4892 | 988.4881 | 1.11 |
| b10 | 1101.5733 | 1101.5729 | 0.36 |
| b11 | 1214.6574 | 1214.6561 | 1.07 |
| b12 | 1361.7258 | 1361.7242 | 1.22 |
| b13 | 1460.7942 | 1460.7936 | 0.41 |
| b14 | 1607.8626 | 1607.8611 | 0.93 |
| y4 | 532.3247 | 532.3240 | 1.31 |
| y5 | 679.3932 | 679.3923 | 1.32 |
| y6 | 778.4616 | 778.4606 | 1.28 |
| y7 | 925.5300 | 925.5289 | 1.19 |
| y8 | 1038.6140 | 1038.6127 | 1.25 |
| y9 | 1151.6981 | 1151.6968 | 1.13 |

C

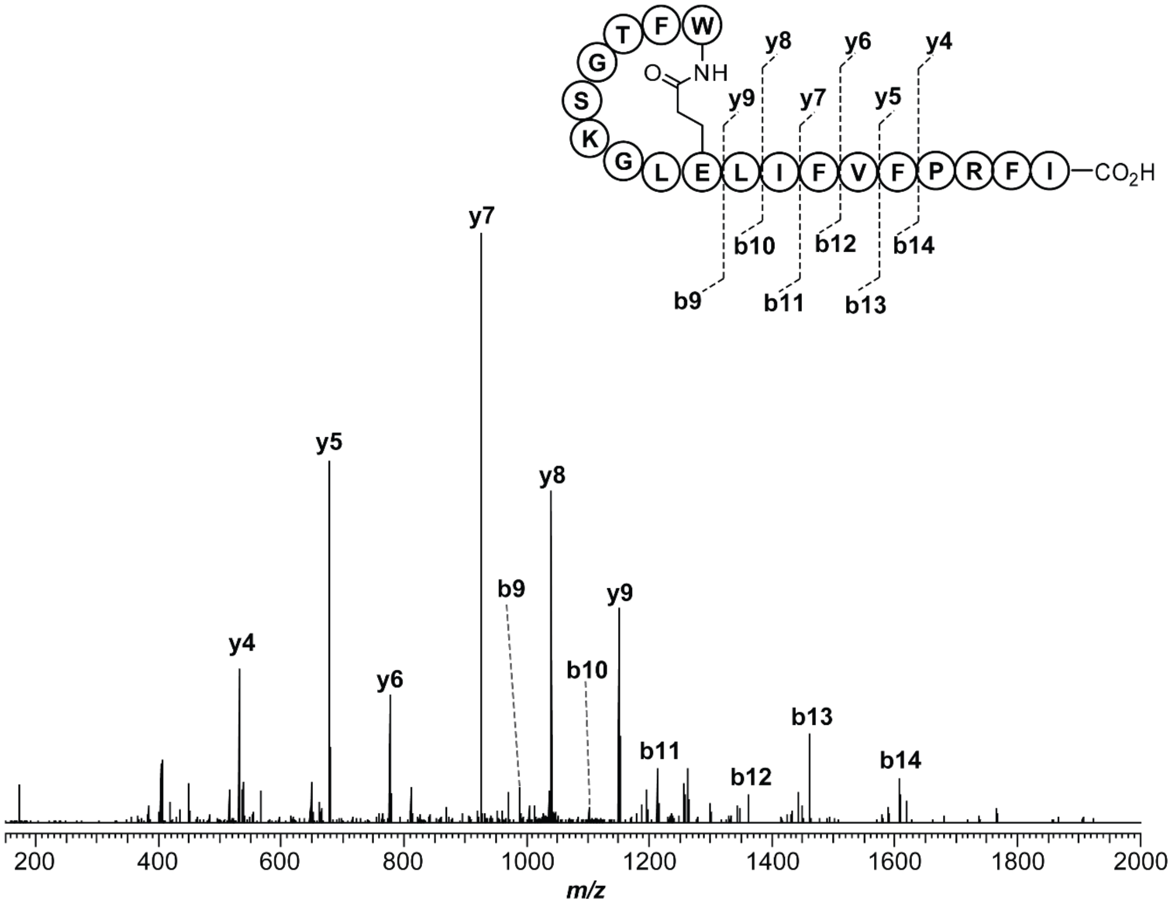

**Figure S21. High-resolution and tandem MS of FTGSK variant produced from CFB. (A)** High-resolution broadband spectrum of fusilassin FTGSK variant, produced from CFB. **(B)** Mass assignment for the $b^+$ and $y^+$ ions generated from the CID of fusilassin FTGSK variant. **(C)** Tandem MS spectrum of fusilassin FTGSK variant, consistent with a Trp1Glu9 macrolactam.

**A**

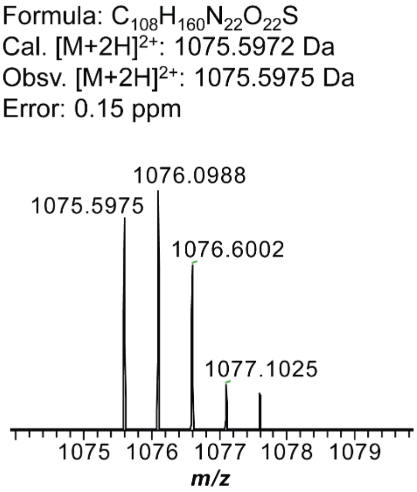

**B**

| Ion | Calculated mass (Da) | Observed mass (Da) | Error (ppm) |
|---|---|---|---|
| b9 | 999.4974 | 999.4968 | 0.60 |
| b10 | 1112.5814 | 1112.5808 | 0.54 |
| b11 | 1225.6655 | 1225.6644 | 0.90 |
| b12 | 1372.7339 | 1372.7332 | 0.51 |
| b13 | 1471.8023 | 1471.8020 | 0.20 |
| b14 | 1618.8707 | 1618.8706 | 0.06 |
| y4 | 532.3247 | 532.3242 | 0.94 |
| y5 | 679.3932 | 679.3926 | 0.88 |
| y6 | 778.4616 | 778.4611 | 0.64 |
| y7 | 925.5300 | 925.5293 | 0.76 |
| y8 | 1038.6140 | 1038.6136 | 0.39 |
| y9 | 1151.6981 | 1151.6979 | 0.17 |

**C**

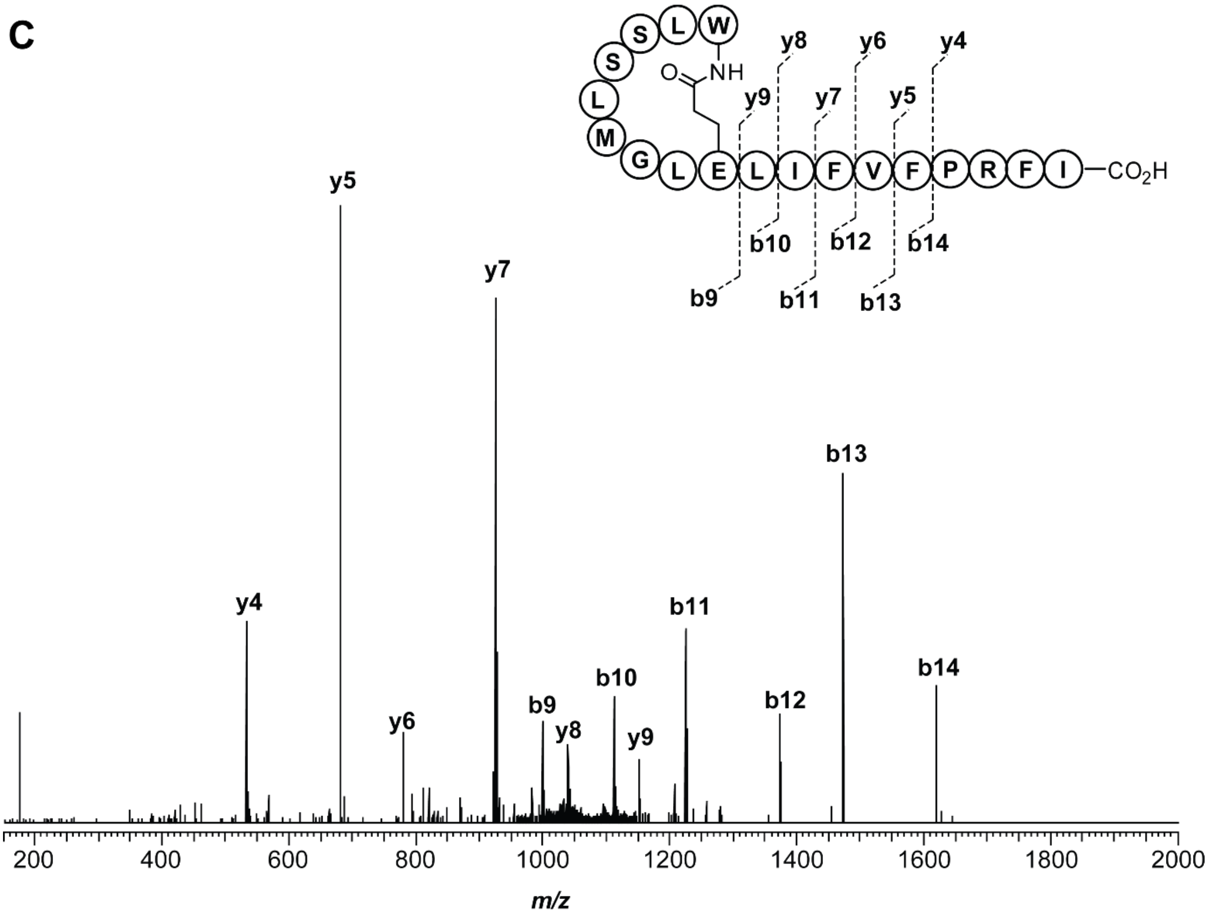

**Figure S22. High-resolution and tandem MS of LSSLM variant produced from CFB. (A)** High-resolution broadband spectrum of fusilassin LSSLM variant, produced from CFB. **(B)** Mass assignment for the $b^+$ and $y^+$ ions generated from the CID of fusilassin LSSLM variant. **(C)** Tandem MS spectrum of fusilassin LSSLM variant, consistent with a Trp1Glu9 macrolactam.

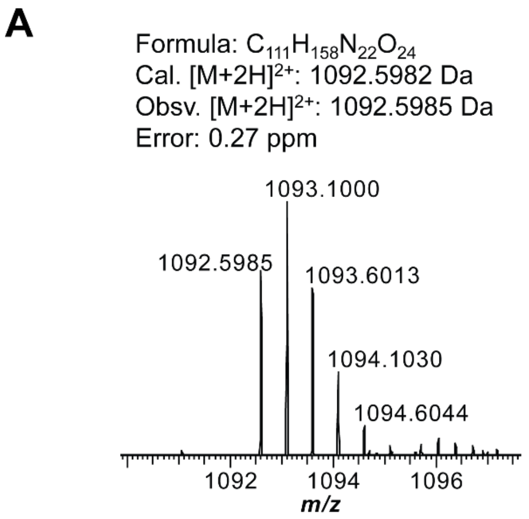

**B**

| Ion | Calculated mass (Da) | Observed mass (Da) | Error (ppm) |
|---|---|---|---|
| b9 | 1033.4995 | 1033.4989 | 0.58 |
| b10 | 1146.5835 | 1146.5828 | 0.61 |
| b11 | 1259.6676 | 1259.6667 | 0.71 |
| b12 | 1406.7360 | 1406.7357 | 0.21 |
| b13 | 1505.8044 | 1505.8044 | 0 |
| b14 | 1652.8728 | 1652.8719 | 0.54 |
| y4 | 532.3247 | 532.3243 | 0.75 |
| y5 | 679.3932 | 679.3926 | 0.88 |
| y6 | 778.4616 | 778.4611 | 0.64 |
| y7 | 925.5300 | 925.5294 | 0.65 |
| y8 | 1038.6140 | 1038.6133 | 0.67 |
| y9 | 1151.6981 | 1151.6974 | 0.61 |

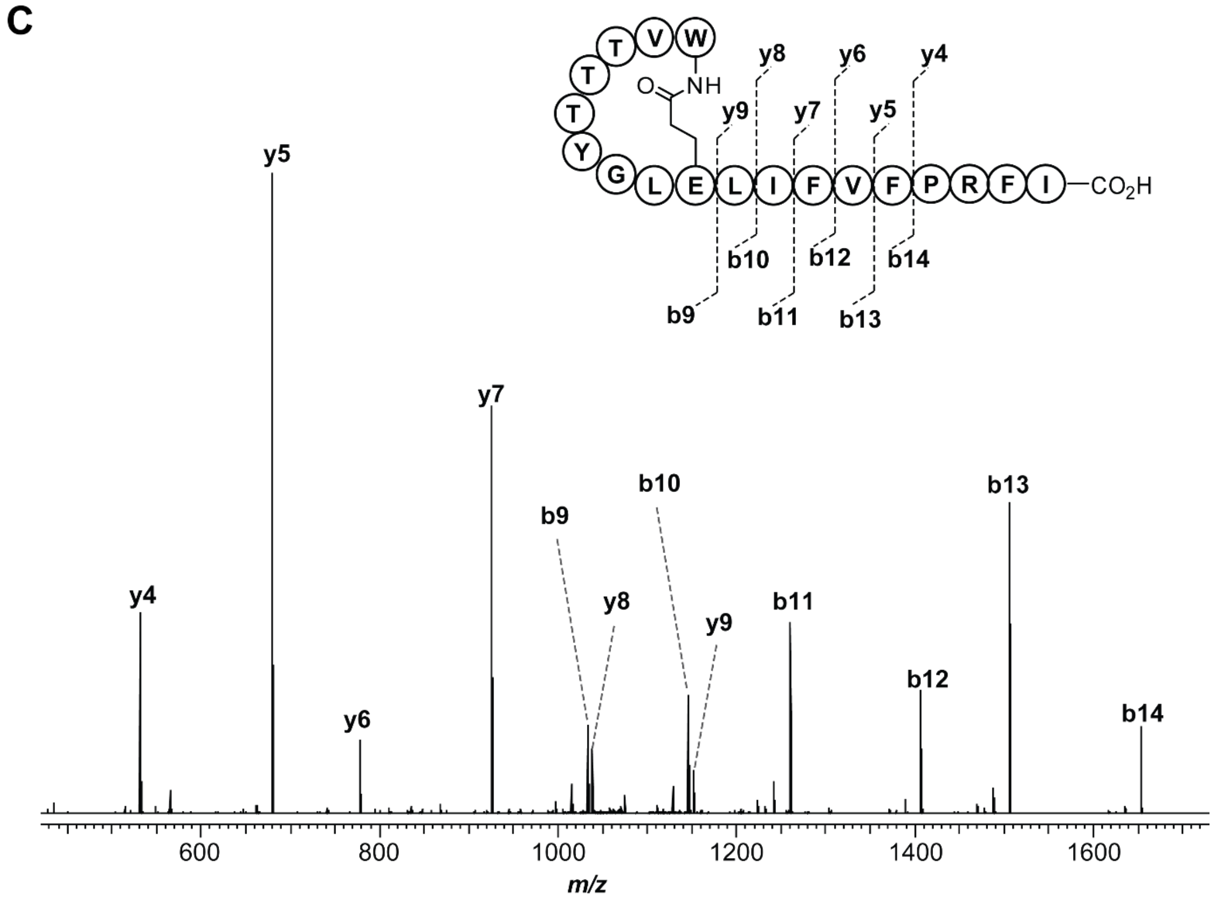

**Figure S23. High-resolution and tandem MS of VTTTY variant produced from CFB. (A)** High-resolution broadband spectrum of fusilassin VTTTY variant, produced from CFB. **(B)** Mass assignment for the $b^+$ and $y^+$ ions generated from the CID of fusilassin VTTTY variant. **(C)** Tandem MS spectrum of fusilassin VTTTY variant, consistent with a Trp1Glu9 macrolactam.

**A**

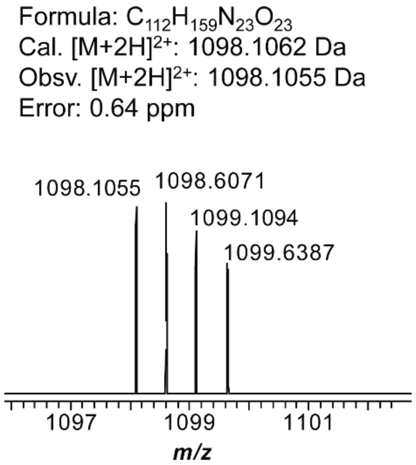

**B**

| Ion | Calculated mass (Da) | Observed mass (Da) | Error (ppm) |
|---|---|---|---|
| b9 | 1044.5155 | 1044.5135 | 1.91 |
| b10 | 1157.5995 | 1157.5972 | 1.99 |
| b11 | 1270.6836 | 1270.6809 | 1.97 |
| b12 | 1417.7520 | 1417.7499 | 1.48 |
| b13 | 1516.8204 | 1516.8185 | 1.25 |
| b14 | 1663.8888 | 1663.8861 | 1.62 |
| y4 | 532.3247 | 532.3237 | 1.88 |
| y5 | 679.3932 | 679.3918 | 2.06 |
| y6 | 778.4616 | 778.4602 | 1.80 |
| y7 | 925.5300 | 925.5281 | 2.05 |
| y8 | 1038.6140 | 1038.6123 | 1.64 |
| y9 | 1151.6981 | 1151.6963 | 1.56 |

**C**

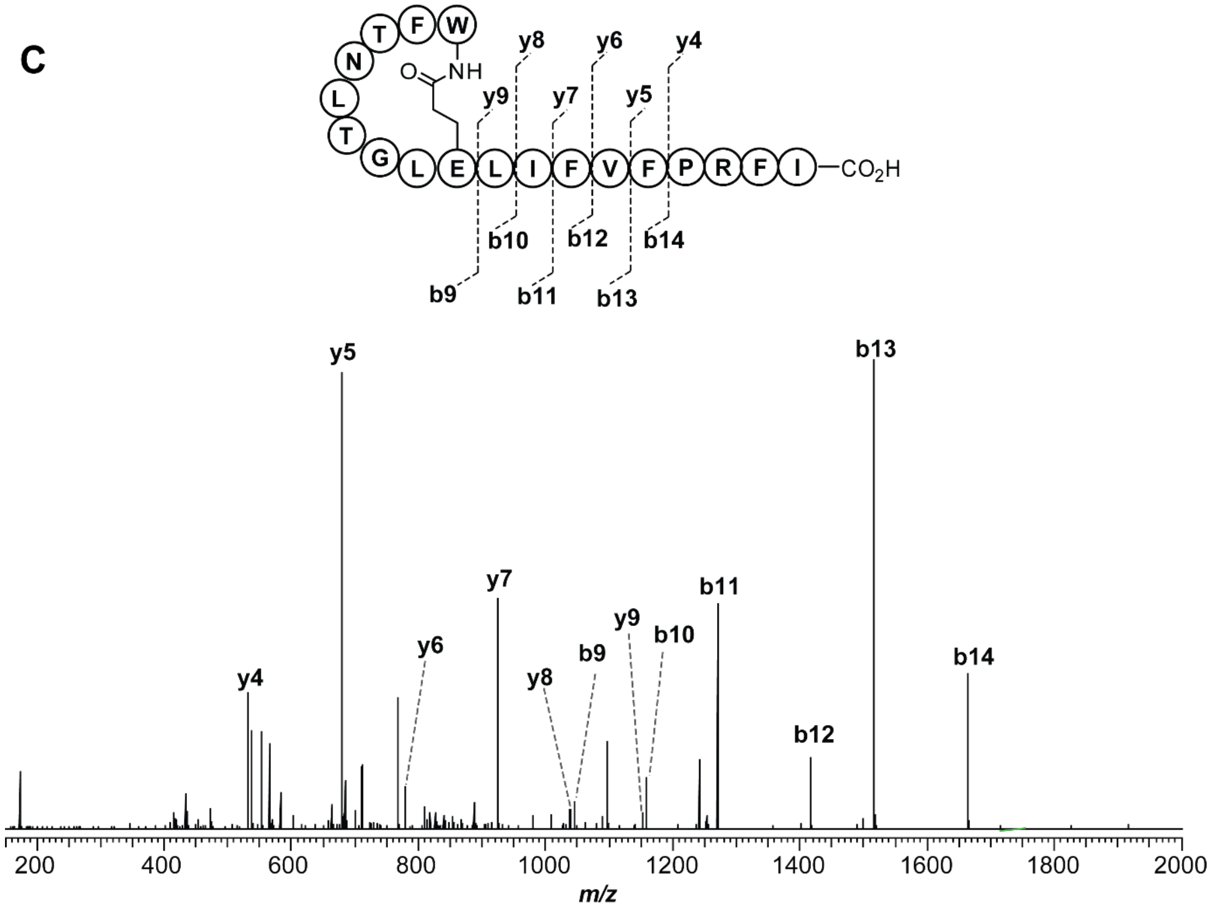

**Figure S24. High-resolution and tandem MS of FTNLT variant produced from CFB. (A)** High-resolution broadband spectrum of fusilassin FTNLT variant, produced from CFB. **(B)** Mass assignment for the $b^+$ and $y^+$ ions generated from the CID of fusilassin FTNLT variant. **(C)** Tandem MS spectrum of fusilassin FTNLT variant, consistent with a Trp1Glu9 macrolactam.

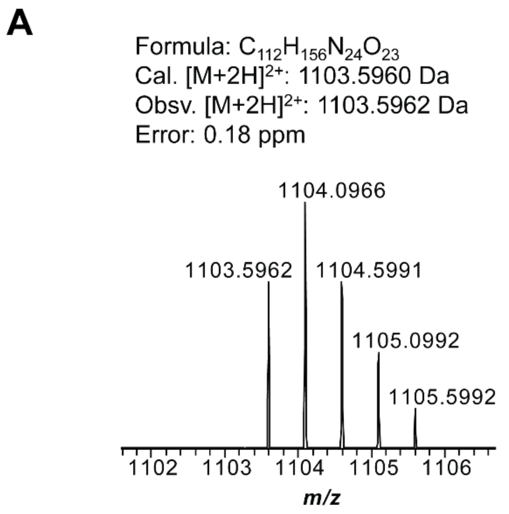

**B**

| Ion | Calculated mass (Da) | Observed mass (Da) | Error (ppm) |
|---|---|---|---|
| b9 | 1055.4951 | 1055.4941 | 0.95 |
| b10 | 1168.5791 | 1168.5781 | 0.86 |
| b11 | 1281.6632 | 1281.6621 | 0.86 |
| b12 | 1428.7316 | 1428.7310 | 0.42 |
| b13 | 1527.8000 | 1527.7998 | 0.13 |
| b14 | 1674.8684 | 1674.8674 | 0.60 |
| y4 | 532.3247 | 532.3242 | 0.94 |
| y5 | 679.3932 | 679.3925 | 1.03 |
| y6 | 778.4616 | 778.4609 | 0.90 |
| y7 | 925.5300 | 925.5292 | 0.86 |
| y8 | 1038.6140 | 1038.6128 | 1.16 |
| y9 | 1151.6981 | 1151.6979 | 0.17 |

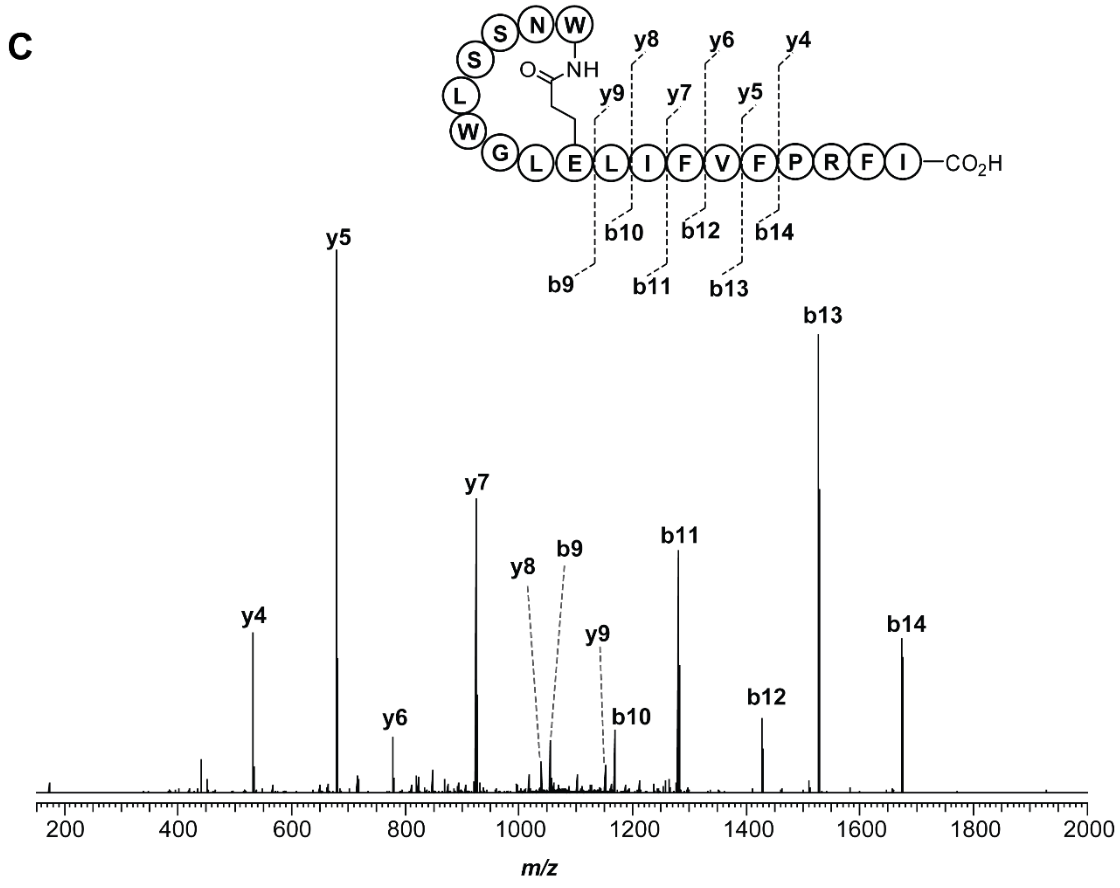

**Figure S25. High-resolution and tandem MS of NSSLW variant produced from CFB. (A)** High-resolution broadband spectrum of fusilassin NSSLW variant, produced from CFB. **(B)** Mass assignment for the $b^+$ and $y^+$ ions generated from the CID of fusilassin NSSLW variant. **(C)** Tandem MS spectrum of fusilassin NSSLW variant, consistent with a Trp1Glu9 macrolactam.

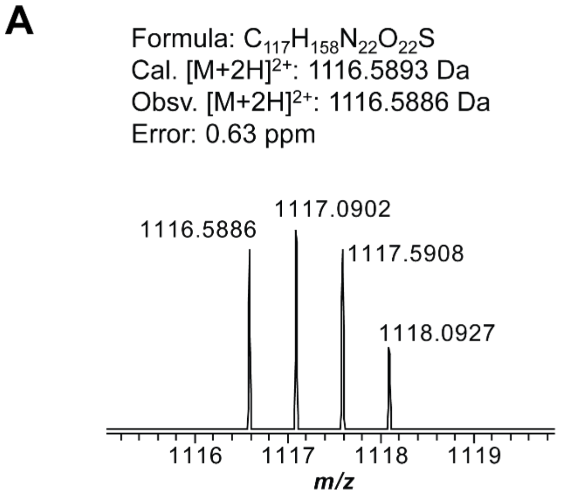

**B**

| Ion | Calculated mass (Da) | Observed mass (Da) | Error (ppm) |
|---|---|---|---|
| b9 | 1081.4817 | 1081.4791 | 2.40 |
| b10 | 1194.5658 | 1194.5634 | 2.01 |
| b11 | 1307.6498 | 1307.6473 | 1.91 |
| b12 | 1454.7183 | 1454.7156 | 1.86 |
| b13 | 1553.7867 | 1553.7845 | 1.42 |
| b14 | 1700.8551 | 1700.8512 | 2.29 |
| y4 | 532.3247 | 532.3237 | 1.88 |
| y5 | 679.3932 | 679.3918 | 2.06 |
| y6 | 778.4616 | 778.4606 | 1.28 |
| y7 | 925.5300 | 925.5281 | 2.05 |

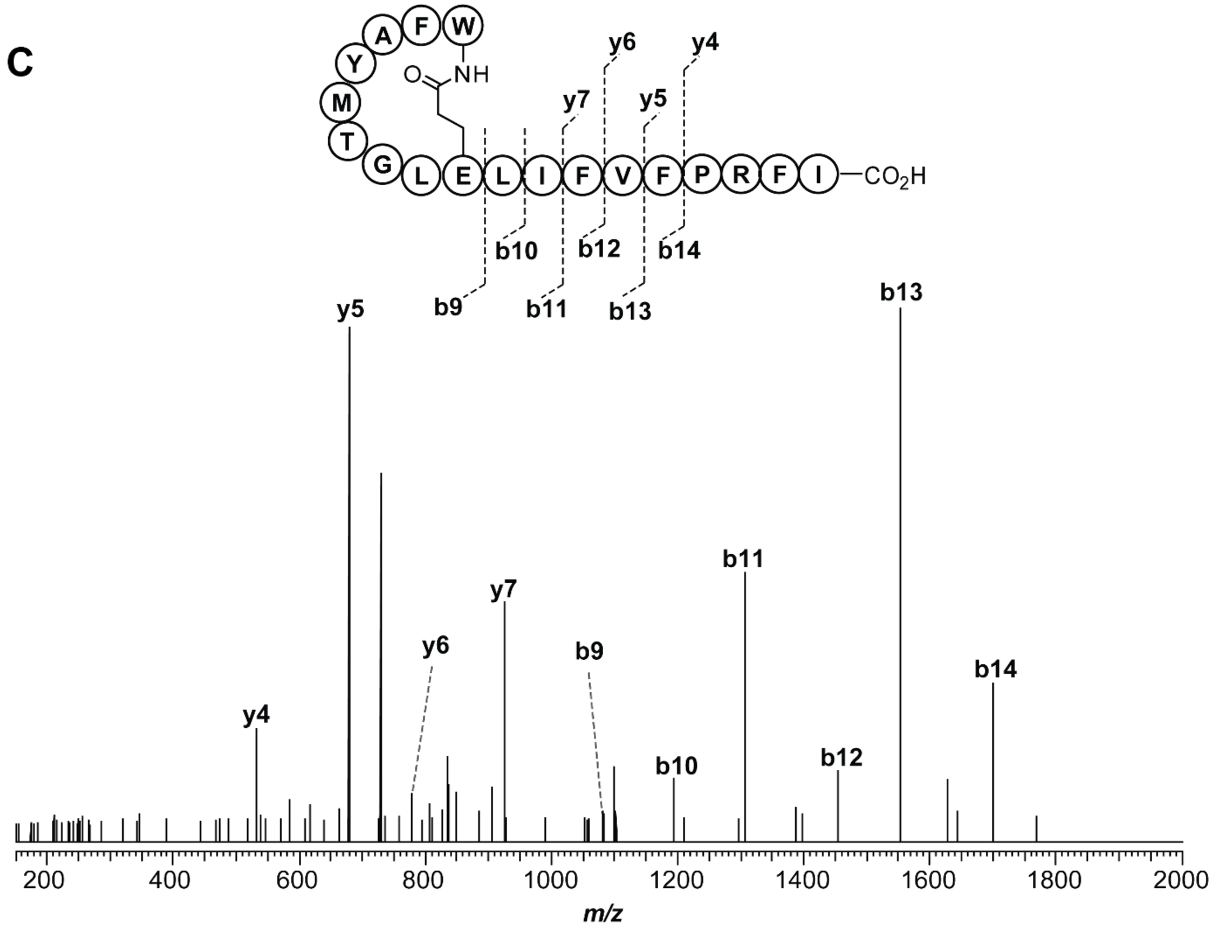

**Figure S26. High-resolution and tandem MS of FAYMT variant produced from CFB. (A)** High-resolution broadband spectrum of fusilassin FAYMT variant, produced from CFB. **(B)** Mass assignment for the $b^+$ and $y^+$ ions generated from the CID of fusilassin FAYMT variant. **(C)** Tandem MS spectrum of fusilassin FAYMT variant, consistent with a Trp1Glu9 macrolactam.

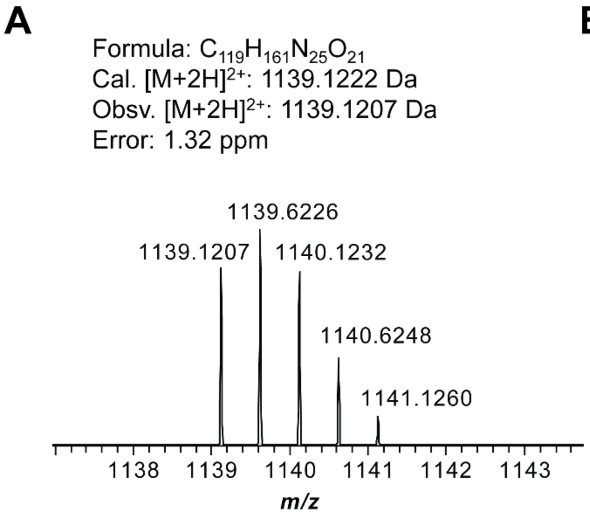

**B**

| Ion | Calculated mass (Da) | Observed mass (Da) | Error (ppm) |
|---|---|---|---|
| b9 | 1126.5474 | 1126.5454 | 1.78 |
| b10 | 1239.6315 | 1239.6294 | 1.69 |
| b11 | 1352.7156 | 1352.7134 | 1.63 |
| b12 | 1499.7840 | 1499.7825 | 1.00 |
| b13 | 1598.8524 | 1598.8503 | 1.31 |
| b14 | 1745.9208 | 1745.9175 | 1.89 |
| y4 | 532.3247 | 532.3237 | 1.88 |
| y5 | 679.3932 | 679.3920 | 1.77 |
| y6 | 778.4616 | 778.4602 | 1.80 |
| y7 | 925.5300 | 925.5283 | 1.84 |
| y8 | 1038.6140 | 1038.6121 | 1.83 |
| y9 | 1151.6981 | 1151.6963 | 1.56 |

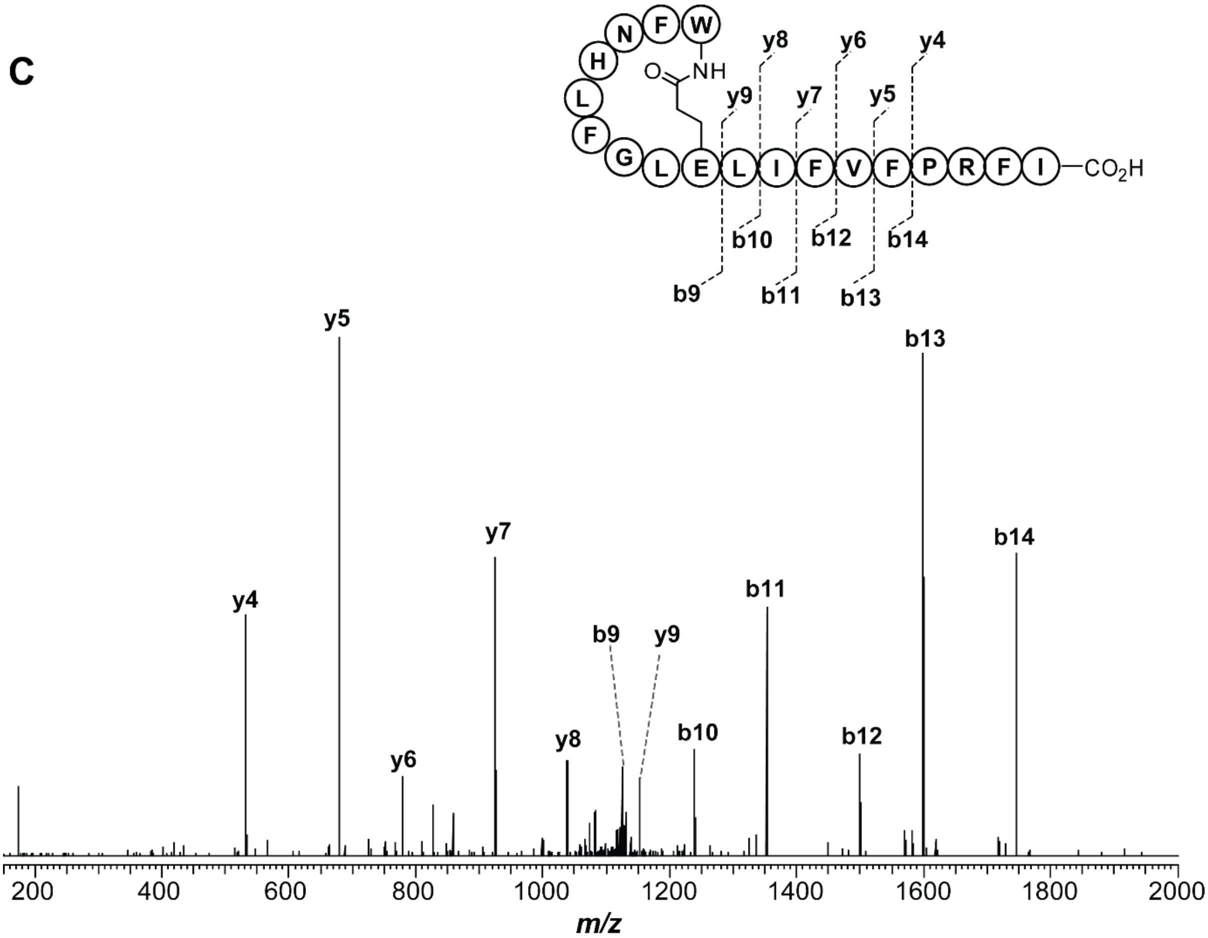

**Figure S27. High-resolution and tandem MS of FNHLF variant produced from CFB. (A)** High-resolution broadband spectrum of fusilassin FNHLF variant, produced from CFB. **(B)** Mass assignment for the $b^+$ and $y^+$ ions generated from the CID of fusilassin FNHLF variant. **(C)** Tandem MS spectrum of fusilassin FNHLF variant, consistent with a Trp1Glu9 macrolactam.

**A**

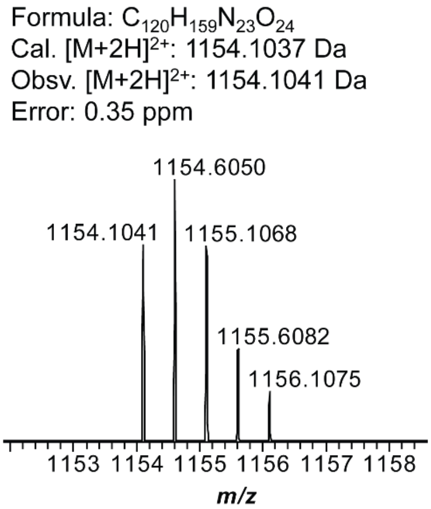

**B**

| Ion | Calculated mass (Da) | Observed mass (Da) | Error (ppm) |
|---|---|---|---|
| b9 | 1156.5104 | 1156.5093 | 0.78 |
| b10 | 1269.5944 | 1269.5936 | 0.63 |
| b11 | 1382.6785 | 1382.6776 | 0.65 |
| b12 | 1529.7469 | 1529.7467 | 0.13 |
| b13 | 1628.8153 | 1628.8147 | 0.37 |
| b14 | 1775.8837 | 1775.8818 | 1.07 |
| y4 | 532.3247 | 532.3242 | 0.94 |
| y5 | 679.3932 | 679.3926 | 0.88 |
| y6 | 778.4616 | 778.4609 | 0.90 |
| y7 | 925.5300 | 925.5294 | 0.65 |
| y8 | 1038.6140 | 1038.6133 | 0.67 |
| y9 | 1151.6981 | 1151.6976 | 0.43 |

**C**

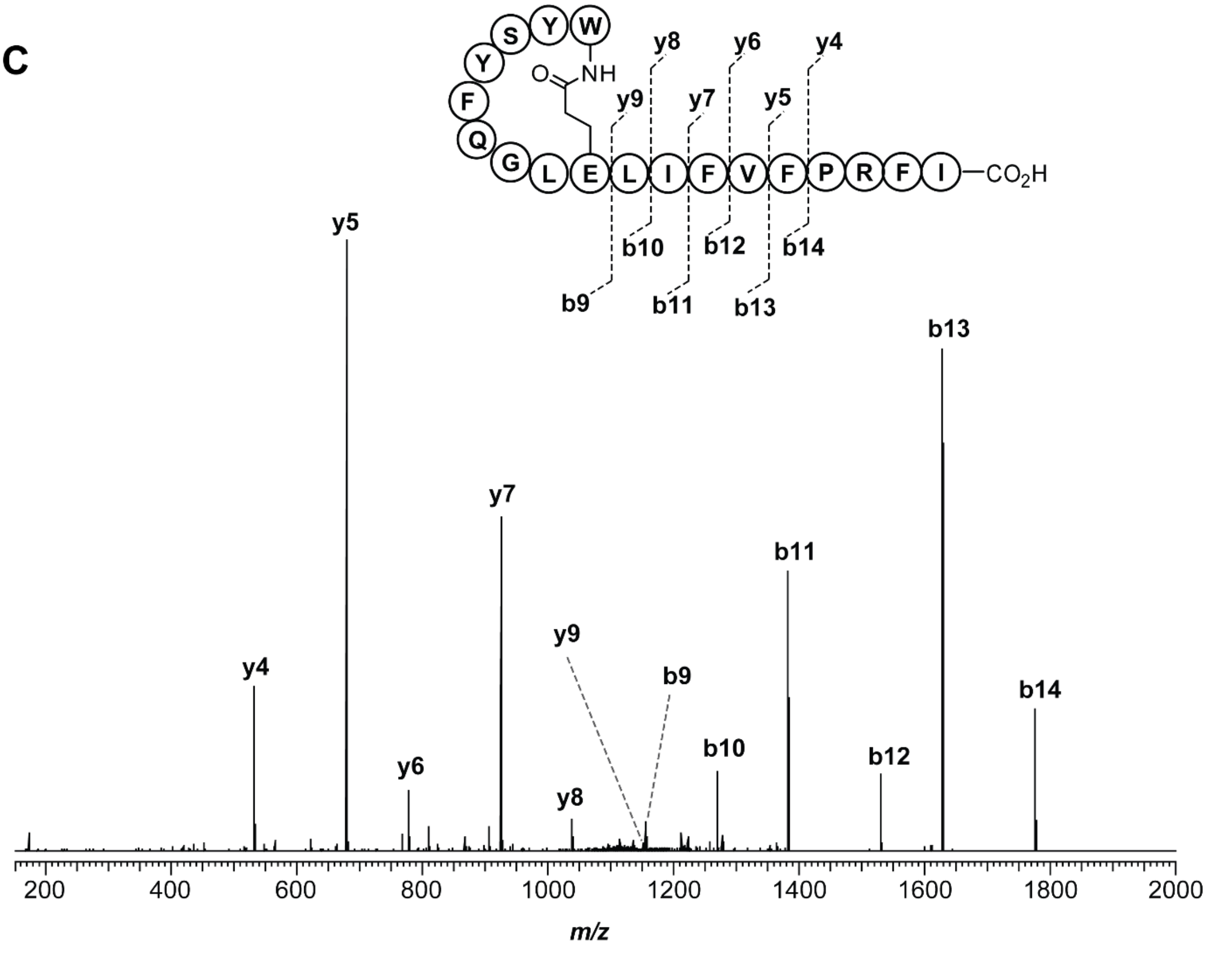

**Figure S28. High-resolution and tandem MS of YSYFQ variant produced from CFB. (A)** High-resolution broadband spectrum of fusilassin YSYFQ variant, produced from CFB. **(B)** Mass assignment for the b$^+$ and y$^+$ ions generated from the CID of fusilassin YSYFQ variant. **(C)** Tandem MS spectrum of fusilassin YSYFQ variant, consistent with a Trp1Glu9 macrolactam.

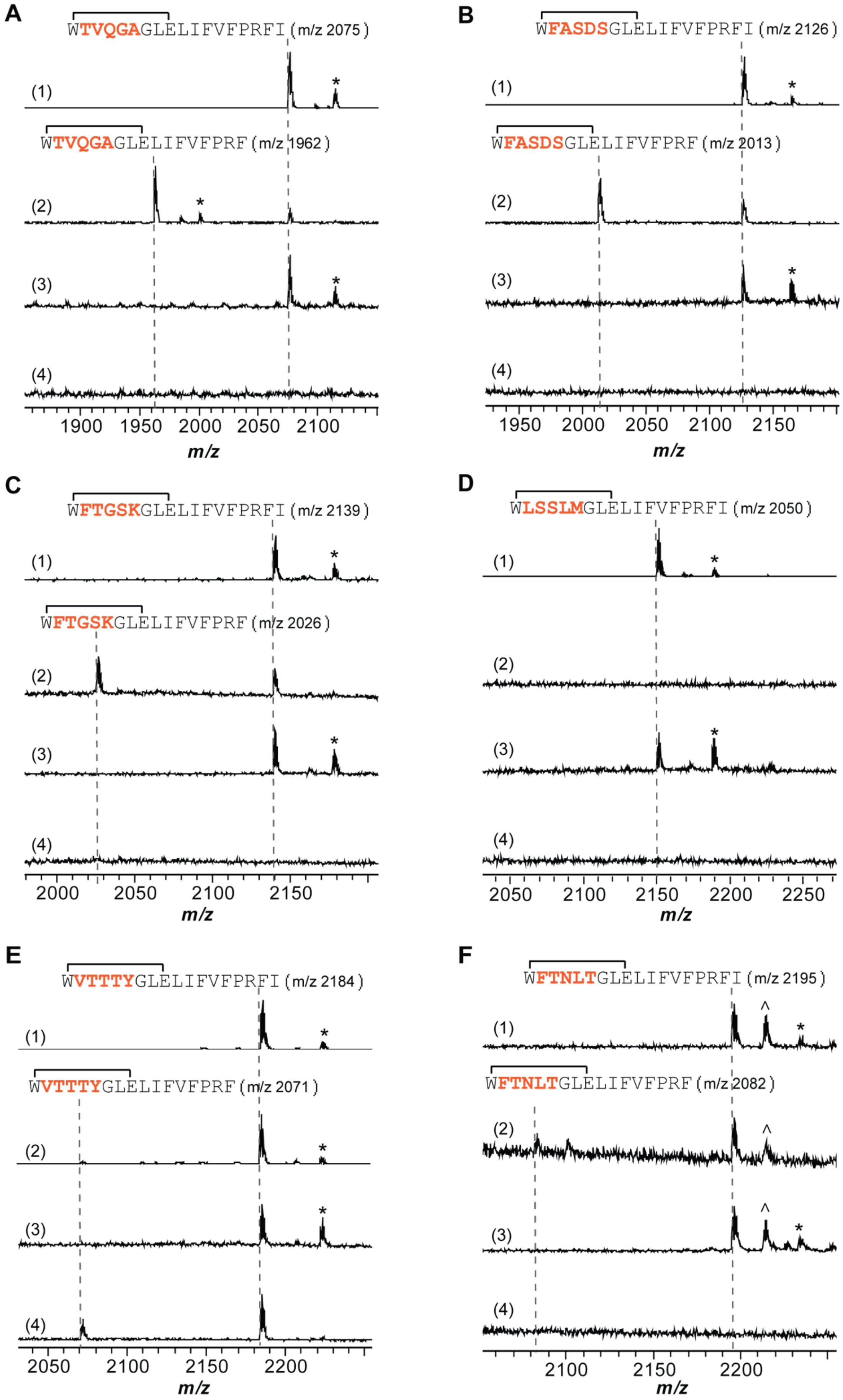
A
WTVQGAGLELIFVFPRFI (m/z 2075)
WTVQGAGLELIFVFPRF (m/z 1962)
(1)
(2)
(3)
(4)
1900 1950 2000 2050 2100
m/z
B
WFASDSGLELIFVFPRFI (m/z 2126)
WFASDSGLELIFVFPRF (m/z 2013)
1950 2000 2050 2100 2150
m/z
C
WFTGSKGLELIFVFPRFI (m/z 2139)
WFTGSKGLELIFVFPRF (m/z 2026)
2000 2050 2100 2150
m/z
D
WLSSLMGLELIFVFPRFI (m/z 2050)
2050 2100 2150 2200 2250
m/z
E
WVTTTYGLELIFVFPRFI (m/z 2184)
WVTTTYGLELIFVFPRF (m/z 2071)
2050 2100 2150 2200
m/z
F
WFTNLTGLELIFVFPRFI (m/z 2195)
WFTNLTGLELIFVFPRF (m/z 2082)
2100 2150 2200
m/z

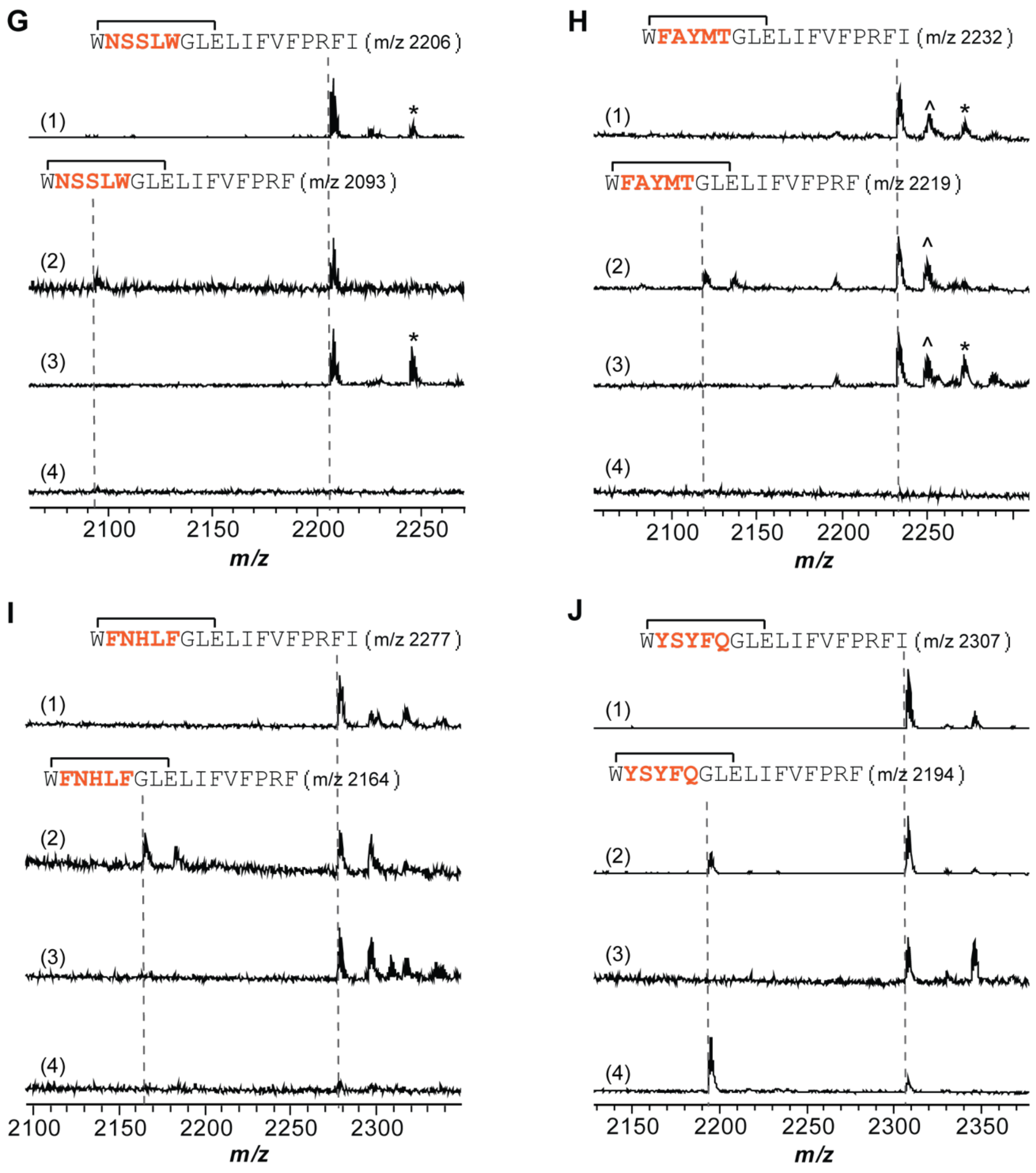

**Figure S29. Carboxypeptidase Y digestion for fusilassin ring variants.** FusA ring variants are produced through CFB (1), and the corresponding lasso peptide was treated under 3 different conditions: carboxypeptidase Y digestion for 18 h (2); heat treatment at 95 °C for 2 h (3); heat treatment at 95 °C for 2 h followed by carboxypeptidase Y digestion for 18 h at room temperature (4). The sequence of each FusA ring variant is shown in the graph and the variation sites are orange-colored. (A) TVQGA variant; (B) FASDS variant; (C) FTGSK variant; (D) LSSLM variant; (E) VTTTY variant; (F) FTNLT variant; (G) NSSLW variant; (H) FAYMT variant; (I) FNHLF variant; (J) YSYFQ variant. The absence/presence of intact lasso peptide upon treatment under condition (4) and the presence of intact lasso peptide under condition (2) and (3) indicate the threaded topology of lasso peptide.

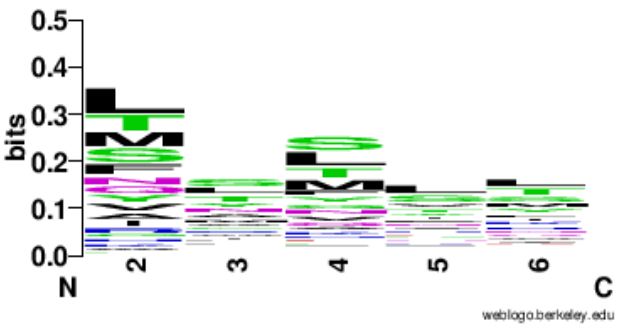

**Figure S30. Sequence logo of sequence-confirmed FusC substrates.** The logo was created using 280 sequence confirmed FusC substrates in WebLogo version 2.8.2.[18]

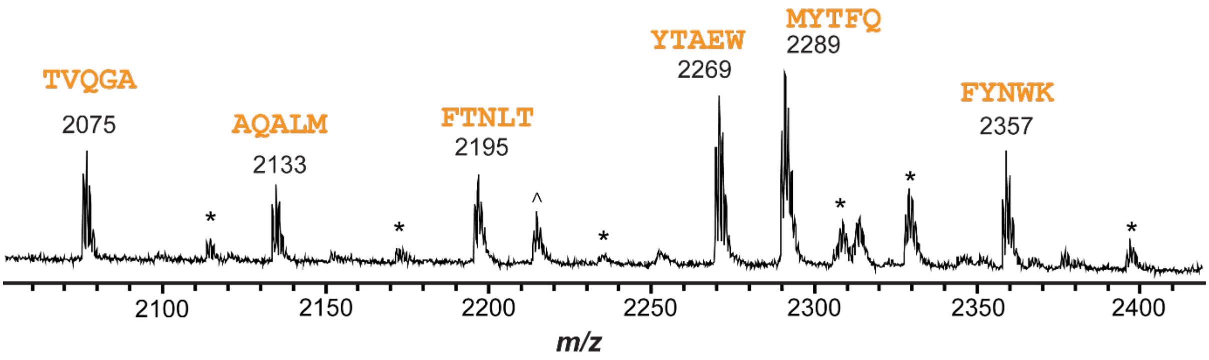

**Figure S31. Simultaneous production of multiple fusilassin variants using CFB.** MALDI-TOF mass spectrum of six fusilassin ring variants produced in a single CFB reaction. The sequences of the varied region (residues 2-6) are highlighted in orange. The mass label corresponds to the $[M+H]^+$ ion of the lasso peptide. * indicates the $[M+K]^+$ ion for the lasso peptide. ^ indicates the uncyclized linear core peptide.

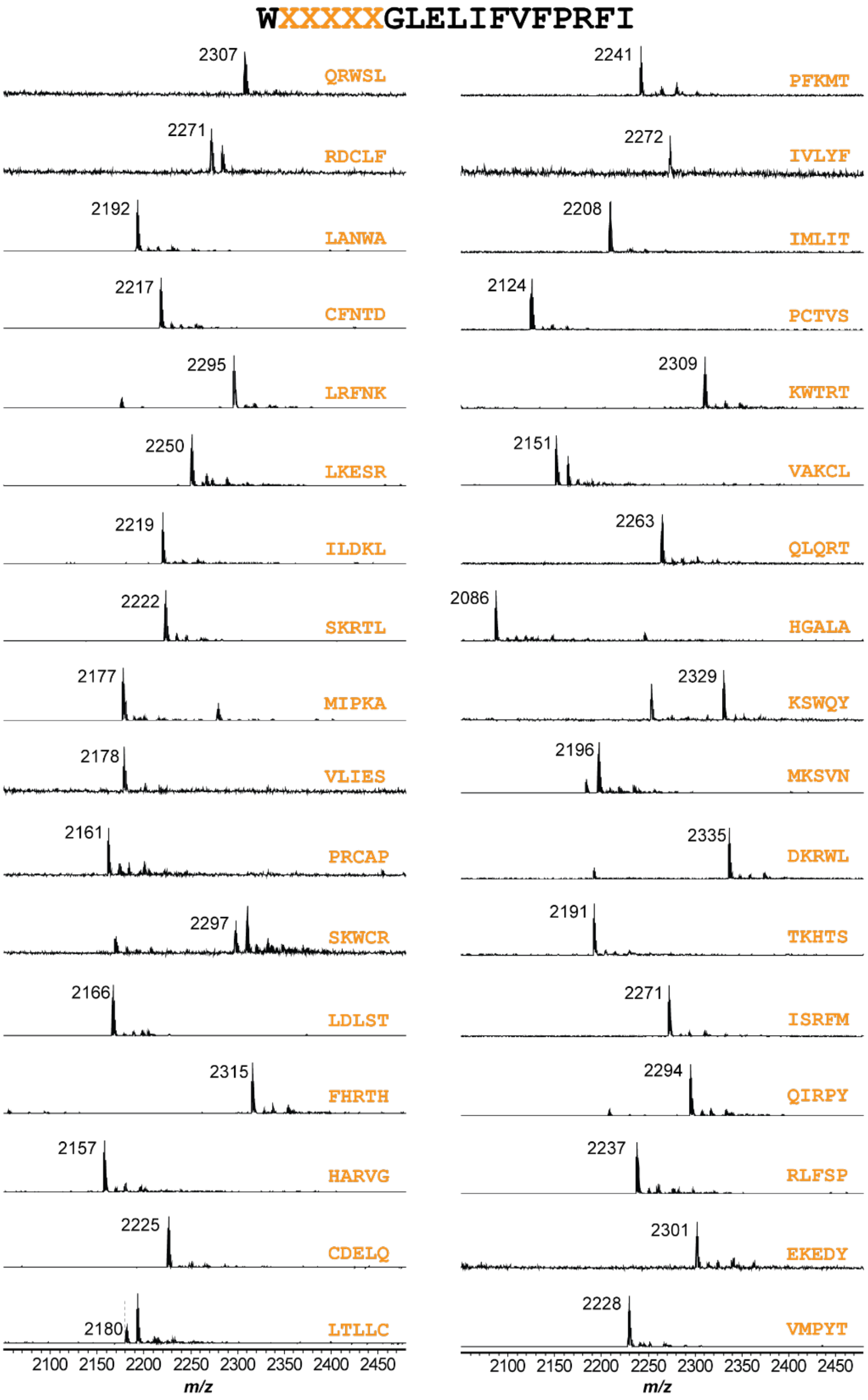

**Figure S32. MALDI-TOF-MS spectra of the uncyclized linear core peptides of the FusC non-substrates produced from PURExpress.** 154 FusC non-substrates that showed no signal from CFB were sequenced, and their corresponding linear core peptides were detected by MALDI-TOF-MS from the PURExpress reaction, in which purified FusB, C, and E were added, indicating that FusB/E was able to cleave the leader peptide, while FusC failed to cyclize the cleaved core. Orange depicts the varied sequence of positions 2-6 of the FusA core region. The mass label corresponds to the $[M+H]^+$ ion of the uncyclized linear core peptide. Figure continues onto the next 2 pages.

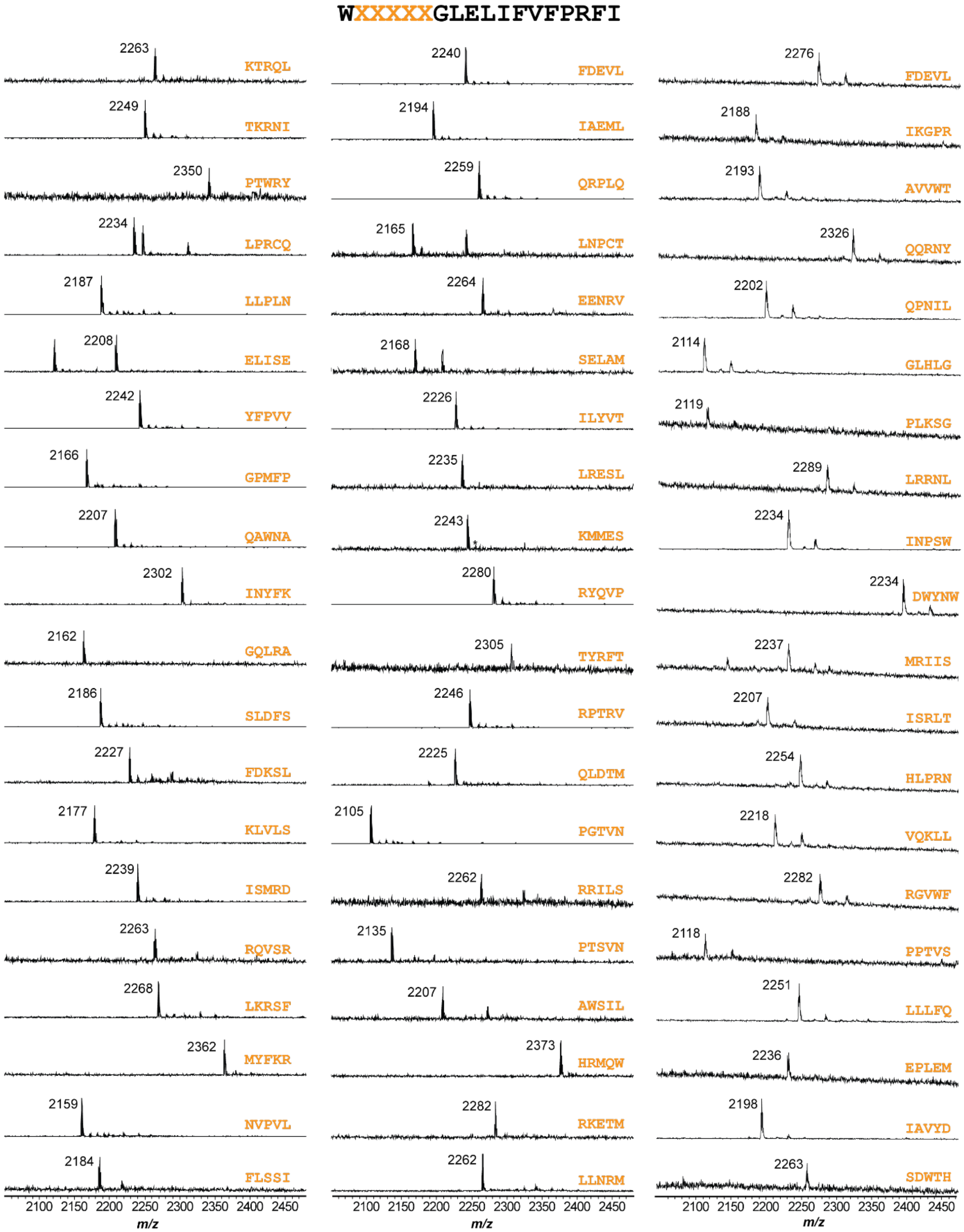

**Figure S32.** *Continued from previous page (2 of 3)*

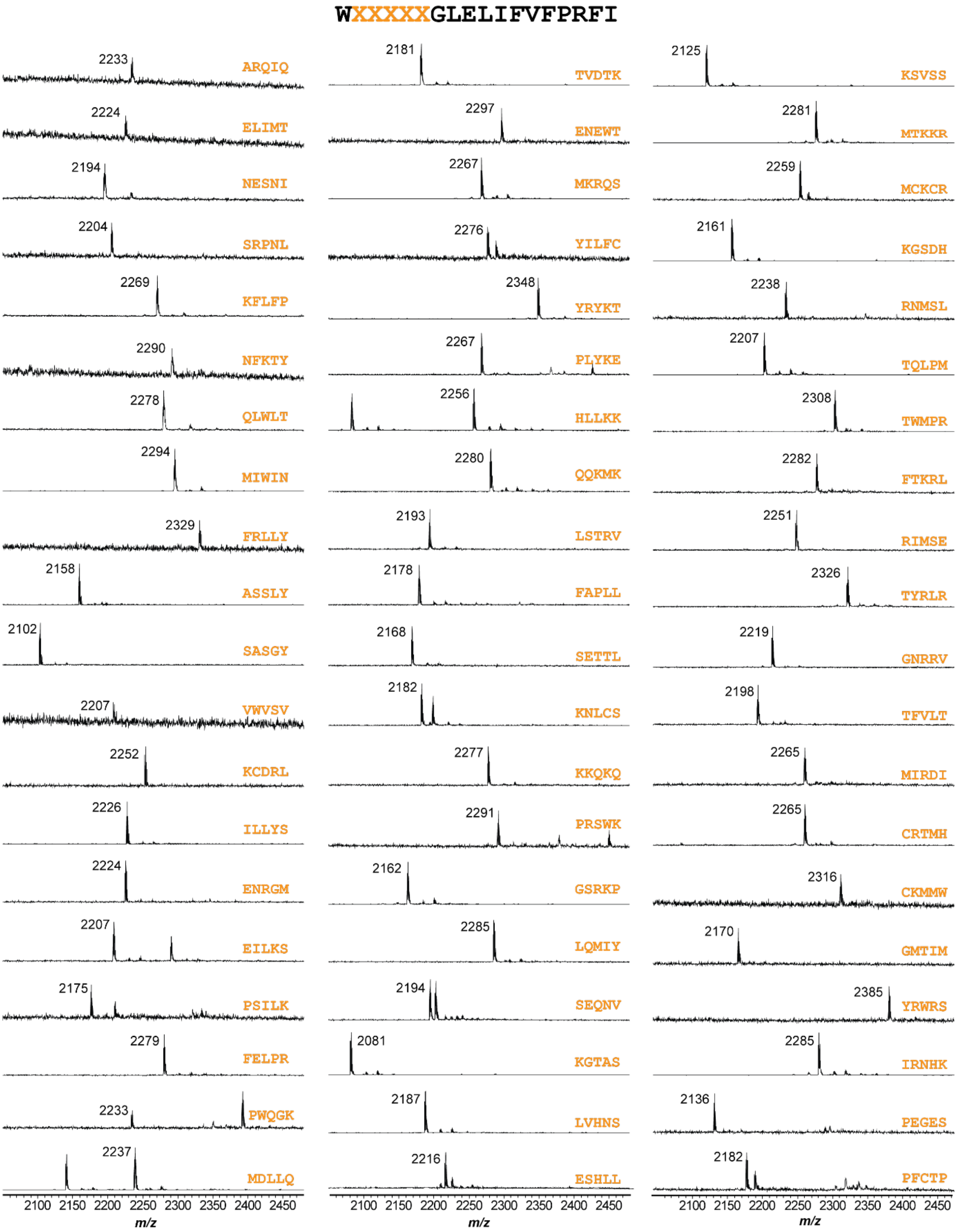

**Figure S32.** *Continued from previous page (3 of 3)*

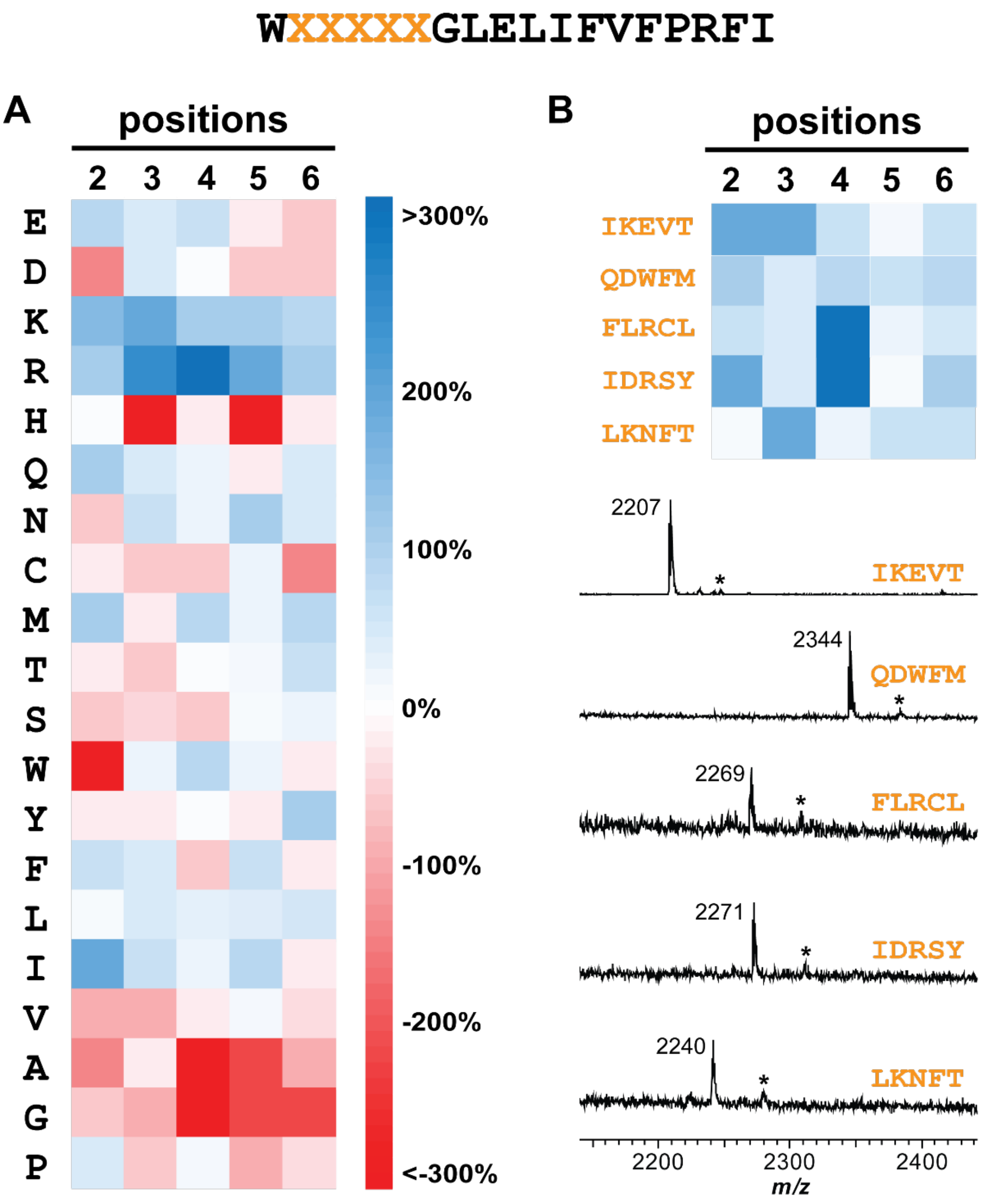

**Figure S33. Heatmap analysis of FusC non-substrates. (A)** The heatmap shows the percent difference between the observed occurrence of each residue on each position with expected occurrence based on the NNK codon. The analysis was conducted based on 154 sequence-confirmed non-substrates. The values are indicated on the color scheme. Blue-colored residues appeared more than expected in FusC non-substrates, while red-colored residues appeared less than expected. **(B)** Anticipated FusC non-substrates based on the heatmap. All five residues in designed sequences are predicted to be favored in FusC non-substrates. No signal was detected in CFB. The formation of their corresponding linear core peptides was detected by MALDI-TOF-MS from the PURExpress reaction, in which the precursor peptides were synthesized and reacted with purified FusB, C, and E. Orange depicts the varied sequence of positions 2-6 of the FusA core region. The mass label corresponds to the $[M+H]^+$ ion of the uncyclized linear core peptide. * indicates the $[M+K]^+$ ion of the linear core peptide.

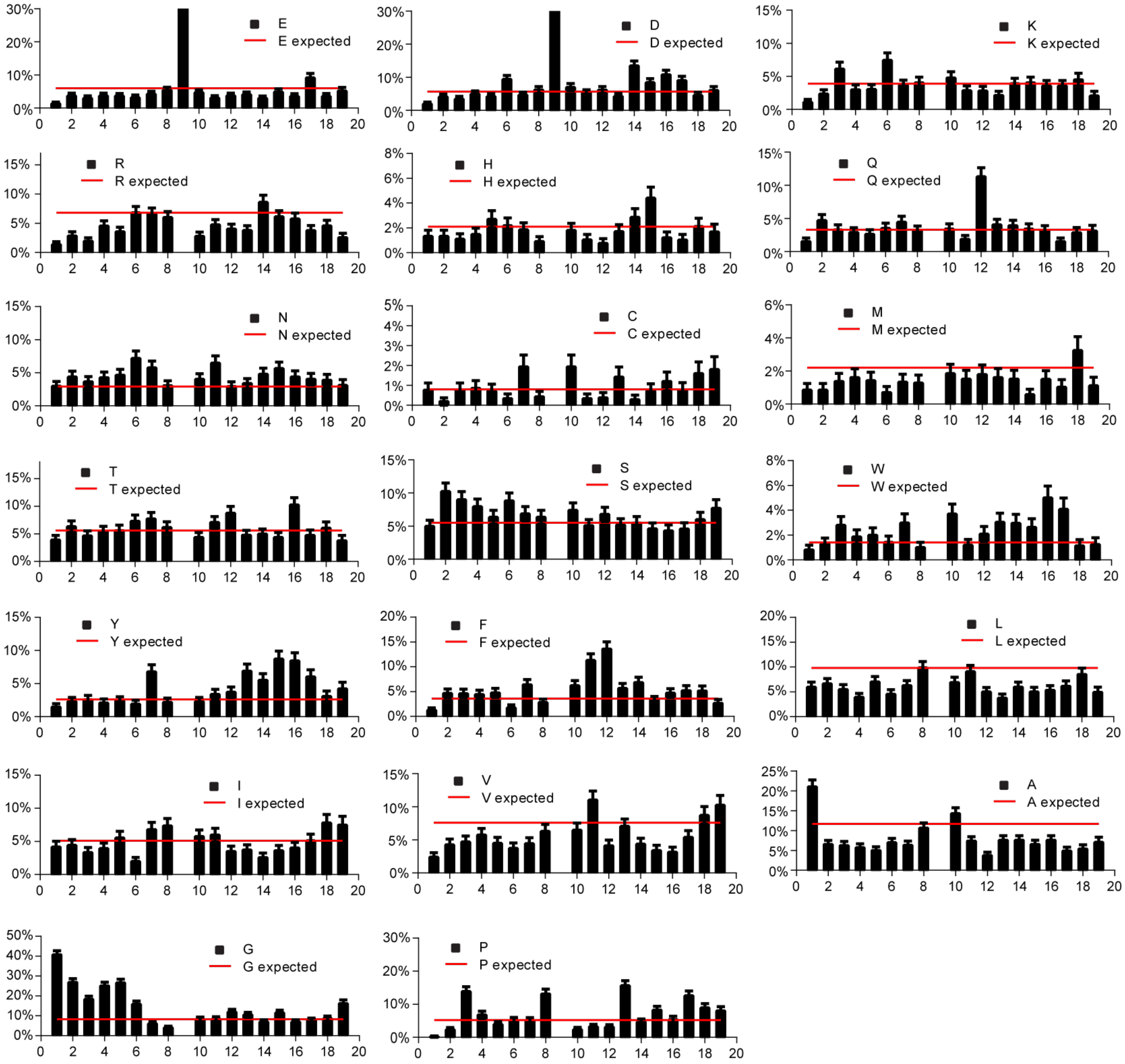

**Figure S34. Position-specific amino acid analysis for lasso peptides.** Shown are the observed frequencies (Y-axis) of the indicated amino acids within the set of unique lasso peptide core sequences (identical cores are only counted once). These values are compared to the expected frequency (red line), derived from the weighted average of the codon frequencies used by the most frequent genus that are predicted to encode lasso peptides (**Table S7).** Error bars represent the 95% confidence interval. X-axis values represent the position in the lasso peptide core region (positions 1 and 9 form the macrolactam).
